# DnaB and DciA: Mechanisms of Helicase Loading and Translocation on ssDNA

**DOI:** 10.1101/2024.11.09.622779

**Authors:** Nicholas Gao, Daniele Mazzoletti, Adele Peng, Paul Dominic B. Olinares, Castrese Morrone, Andrea Garavaglia, Nourelhoda Gouda, Sergey Tsoy, Albert Mendoza, Ahammad Chowdhury, Antonio Cerullo, Hrutvik Bhavsar, Franca Rossi, Menico Rizzi, Brian T Chait, Riccardo Miggiano, David Jeruzalmi

## Abstract

Graphical Abstract
The *Vibrio cholerae* (*Vc*) DnaB replicative helicase structure bound to single-stranded (ss) DNA is depicted in the ribbon (top left) and sphere-cylinder representation (top right). In the bottom center is a native mass spectrum showing Vc DnaB helicase loading onto single-stranded DNA (ssDNA).

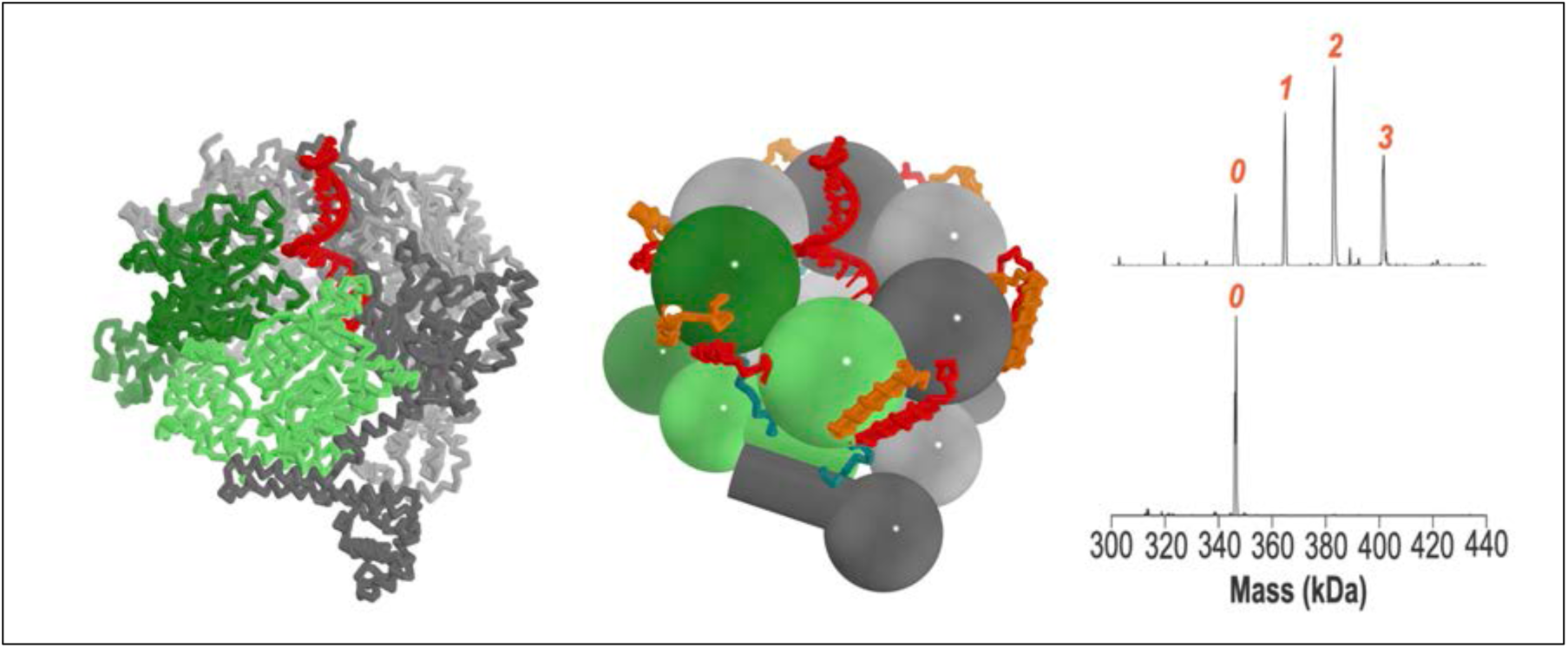

Replicative helicases are assembled on chromosomes by helicase loaders before initiation of DNA replication. Here, we investigate mechanisms used by the bacterial *Vibrio cholerae* (*Vc*) DnaB replicative helicase and the DciA helicase loader. Structural analysis of the ATPγS form of the *Vc*DnaB-ssDNA complex reveals a configuration distinct from that seen with GDP•AlF_4_. With ATPγS, the amino-terminal (NTD) tier, previously found as an open spiral in the GDP•AlF_4_ complex, adopts a closed planar arrangement. Further, the DnaB subunit at the top of the carboxy-terminal spiral (CTD) tier is displaced by ∼25 Å between the two forms. We suggest that remodeling the NTD layer between closed planar and open spiral configurations and migration of two distinct CTDs to the top of the DnaB spiral, repeated three times, mediates hand-over-hand translocation. Biochemical analysis suggests that *Vc*DciA leverages its Lasso domain to contact DnaB near its Docking-Linker-Helix interface. Up to three copies of *Vc*DciA bind to *Vc*DnaB and suppress its ATPase activity during loading onto physiological DNA substrates. Our data suggest that DciA loads DnaB onto DNA using the ring-opening mechanism.

## Introduction

Replisomes in all domains of life rely on replicative helicases to melt chromosomal DNA into single-stranded templates for DNA replication (Figure 1)(1–6). In *E. coli*, the hexameric DnaB replicative helicase is a closed protein ring that encircles and translocates on one strand of DNA, excluding the second, as it prepares templates for DNA synthesis (Figure 1). Members of the DnaB family (7–20) (henceforth: DnaB) and (reviewed in (21–27)) feature a two-tiered architecture, termed the amino-terminal domain (NTD) and the carboxy-terminal domain (CTD); each tier circumscribes a central channel through which ssDNA will traverse. A short linker helix (LH) connects each NTD and CTD; the LH packs against a neighboring CTD’s docking helix (DH) in the complete hexamer. The translocating form of DnaB adopts a right-handed closed-spiral configuration and is suggested to operate using a hand-over-hand mechanism. Translocation occurs akin to a human using two hands to climb a rope, except that DnaB has six hands (10). ATP hydrolysis on a DnaB subunit at the bottom of the spiral (in the pose in Figure 2) initiates translocation. This subunit peels away from its DnaB partners and from ssDNA (10, 24, 28, 29), migrates to the top of the spiral, binds ATP, docks onto a DnaB subunit via its β-hairpin arginine finger, and binds ssDNA position 3ʹ from the original location. Translocation continues via additional subunit migrations from the bottom to the top of the DnaB spiral (10). Crucial support for this model emerged from a series of structures of the DnaB-family helicase Gp4 from phage T7 (24, 29).

**Figure 1.**
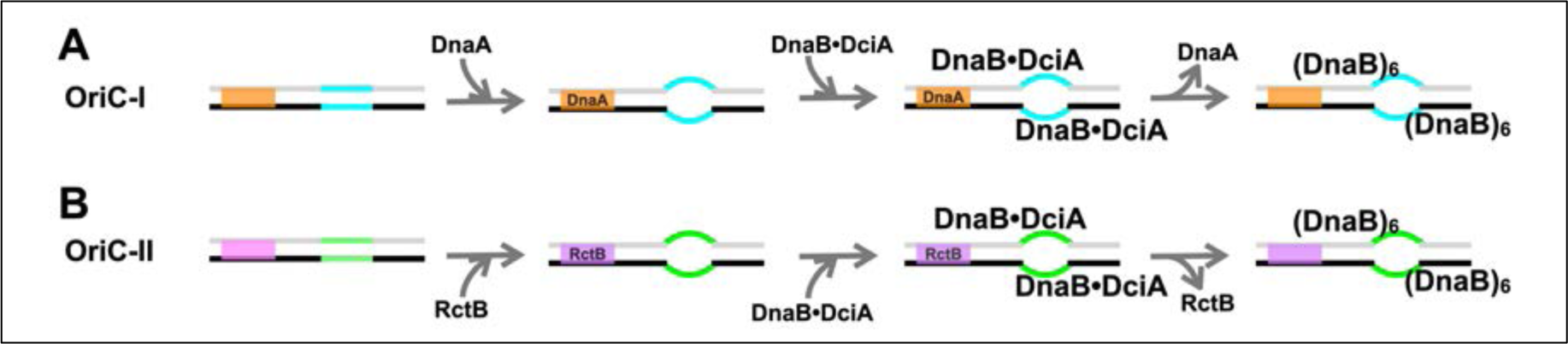
Initiation of DNA Replication in *Vibrio cholerae*. The primary (A) and secondary chromosomes (B) in *Vibrio cholerae* initiate replication via related but distinct mechanisms. Multiple copies of the replication initiator protein (OriC-I: *Vc*DnaA; OriC-II: *Vc*RctB) recognize sites on dsDNA (OriC-I: orange, OriC-II: purple) and oligomerize into a protein-DNA ensemble. Assembly of the initiator-origin DNA complex promotes the melting of an A-T-rich segment at the origin, termed the DNA unwinding element (DUE, OriC-I: cyan, OriC-II: green). The melted DUE segment provides the sites for the *Vc*DciA loader to load the *Vc*DnaB replicative helicase.

**Figure 2.**
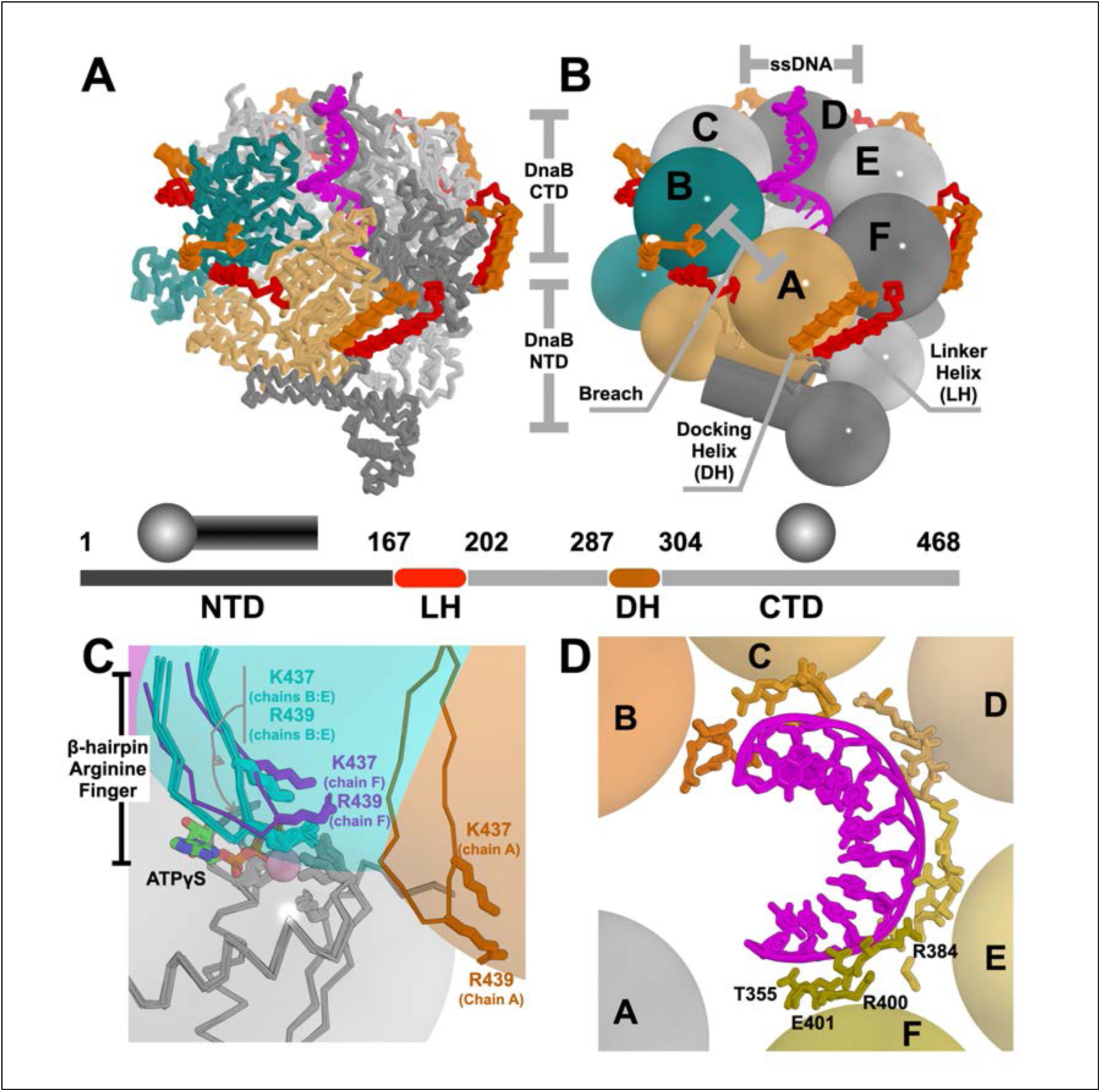
The *Vc*DnaB-ssDNA-ATPγS Complex and Its Interactions with Nucleotide and ssDNA. The *Vc*DnaB-ssDNA-ATPγS is drawn in a ribbon representation (A) or a schematic using a design language wherein the NTDs are depicted as a mushroom shape and the CTDs as spheres (B). In panels A and B, ssDNA, colored in magenta, is depicted in a cartoon and stick representation. The breach in the CTD layer is between chain A (light orange) and chain B (deep teal); other DnaB subunits are colored in alternating shades of gray. The Docking Helix (DH) and the Linker Helix (LH) are depicted as cylinders and ribbons colored orange and red, respectively. The primary sequence of *Vc*DnaB is depicted as a bar, with the salient domain boundaries indicated. C) Close-up of the ATP catalytic center at the interface between each pair of DnaB CTDs. ATP is depicted in the stick representation. The six DnaB CTD pairs, depicted here as spheres, are superimposed on the subunit (colored in grey) that harbors the Walker A and B motifs (not indicated). For clarity, only a portion of the superimposed CTD is shown as a ribbon; the grey sphere shows the full extent of the superimposed CTD. Spheres corresponding to the CTDs excluded from the superposition are colored in purple (chain F), cyan (chains B:E), and orange (chain A). The β-hairpin arginine fingers from all six non-superimposed CTDs are shown in the ribbon format. Although excluded from the superposition, the β-hairpin four corresponding to four chains (chains B, C, D, and E, cyan) superimpose closely. In contrast, chains F (purple) and A (orange) are found ∼3 Å and ∼19 Å, respectively. D) Schematic representation of the *Vc*DnaB-ssDNA interface. The DnaB CTDs are represented as spheres, labeled by the chain, and colored in shades of yellow/orange, except for chain A, which is colored gray. The ssDNA is drawn in a cartoon and stick representation and colored purple. Additional details on the *Vc*DnaB-ssDNA complex can be found in Supplementary Figures 7, 11, and Supplementary Table 4.

Assembly of DnaB on single-stranded DNA (ssDNA) requires distinct mechanisms and loading factors (Supplementary Figure 1). The loading factors include *Escherichia coli* (*Ec coli)* DnaC (30), bacteriophage λP (31), bacteriophage SPP1 G39P (32, 33), *Bacillus subtilis* (*Bst*) DnaI (34), T4 gene 59 (35), and phage P2 B (36). Surprisingly, the genomes of most bacteria lack these well-studied loaders.

Instead, most bacteria employ the structurally and perhaps mechanistically distinct DciA or DopE family members (37–41). Although the amino acid sequences of bacterial helicase loaders diverge significantly, they appear to mediate conserved mechanisms during helicase loading (11, 12, 14, 16, 19, 36, 42–48). Loaders display a conserved overall architecture: a globular domain adjacent to an intrinsically disordered segment that often features an alpha helix at a terminus, termed the lasso/grappling hook. Related arrangements can also be inferred for *Bst* SPP1 phage G39P (49), *Bst* DnaI (16, 50–52), and *Ec* phage T4 gp59 (53) helicase loaders. The DciA helicase loaders are organized around a type II K homology domain (KH) with Lasso elements at either the amino or carboxy terminus of a particular ortholog or both (12, 14, 40, 41, 54).

Cells apply three broad strategies to load hexameric helicases onto nucleic acid substrates: ring-opening, ring-forming, and ring-closing (22, 55, 56) (Supplementary Figure 1 and Supplementary Information). Although much is known about many bacterial loaders (11, 12, 14, 16, 19, 36, 42–46), little is known about DciA helicase loaders and the mechanisms they implement during helicase loading. DciA orthologs from *Vibrio cholerae* (*Vc*), *Mycobacterium tuberculosis* (*Mt*), and *Caulobacter crescentus* (*Cc*) are all essential proteins (37, 42, 57, 58), and they have been shown to bind to cognate DnaB helicases (12, 14, 42, 57, 58). Component KH and Lasso domains contact DnaB (12, 14, 58). The binding site on DnaB for DciA is proposed to encompass the DH-LH interface (12, 14, 58). DciA orthologs bind to DNA (11, 58, 59), a feature shared with other bacterial loaders (31, 35, 47, 60). Suppression of DnaB’s ATPase activity is a hallmark of the bacterial helicase loaders(43, 46, 60); however, *Cc*DciA has been reported to promote ATP hydrolysis (42). DciA activates cognate DnaB’s DNA unwinding activity by mediating assembly on ssDNA, though the underlying mechanisms have yet to be established (40, 41).

To gain insights into the mechanisms operating during the loading of DnaB by the DciA loader and subsequent translocation, we studied the orthologs from *Vibrio cholerae*. The single particle cryogenic electron microscopy (cryo-EM) structure of the *Vc*DnaB-ssDNA-ATPγS (ATPγS: Adenosine 5’-O-(3-thio)triphosphate) complex, the end state of helicase loading, implies that cycles of opening/closing of the NTD layer and CTD remodeling are significant features in the mechanism of translocation on ssDNA. We suggest that remodeling of the NTD layer and distinct migrations of two subunits to the top of the DnaB spiral, repeated three times, mediates hand-over-hand translocation. Native mass spectrometry (nMS) indicates that *Vc*DnaB is a hexamer, and that the stoichiometry of the *Vc*DnaB-DciA-ssDNA complex is 6:3:1. *Vc*DciA features a KH domain at its amino terminus and a disordered segment (lasso) at its carboxy terminus (12, 40, 41, 54, 58). Binding experiments show that the lasso segment specifies most of the binding energy in its interaction with DnaB. Furthermore, *Vc*DciA binds to DnaB at the interface between the docking helix (DH) and linker helix (LH) on adjacent subunits. *Vc*DciA suppresses the intrinsic ATPase activity of DnaB. We show that *Vc*DciA loads DnaB onto ssDNA and activates unwinding. Overall, our data suggest that *Vc*DciA is a ring-opening helicase loader.

## Materials and methods

### Protein Biochemistry

Standard techniques (61) were used to develop the nine proteins used in this study. Our studies encompassed efforts with full-length and designed truncations of DnaB and DciA from *Vibrio cholerae* (Supplementary Figure 2). The DnaB constructs included the full-length sequence (1-468), the DnaB amino-terminal domain (NTD; 1-170), the DnaB amino-terminal domain fused to the Linker Helix (NTD-LH; 1-208), and the Linker helix fused to the DnaB carboxy-terminal domain (CTD; 208-468). The DciA constructs included the complete sequence (1-157), the KH domain (1-98), and the Lasso domains (98-157). The supplementary methods section (Supplementary Methods and Tables 1 and 2) provides complete details on plasmid construction, bacterial expression, and protein purification.

Interactions between DnaB and DciA and their respective truncations were interrogated by co-lysing bacteria that separately expressed each protein, where only one of the proteins was histidine-tagged. The lysate was passed over NiNTA agarose (QIAGEN), washed of contaminants, and histidine-tagged proteins were eluted. Interactions were scored based on the propensity of the untagged protein in the experiment to be retained on Ni-NTA agarose (QIAGEN) compared to trials where the histidine-tagged protein was omitted. The supplementary methods section describes the ‘pull-down’ experiments more completely.

### Surface Plasmon Resonance

Dissociation constants (K_D_) were estimated using surface plasmon resonance (SPR) from measurements of the on and off rates between DnaB and DciA. SPR data were acquired using a Biacore X100 instrument (GE Healthcare/Cytiva), and a CM5 sensor chip functionalized with an anti-histidine antibody according to the manufacturer’s instructions. All measurements were conducted at 25 °C. Since the SPR response is proportional to molecular weight, we chose to immobilize the smaller C-terminal-His-tagged-DciA (MW = 18.4 kDa) while positioning the much larger DnaB (MW = 312.3 kDa) in the mobile phase. DciA was immobilized on the surface of a single channel of the CM5 sensor chip via the interaction between the histidine tag and the anti-histidine antibody. The appropriate ligand density (RL) on the chip was calculated according to the equation: 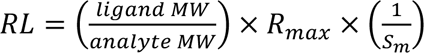 where R_max_ is the maximum binding signal, and S_m_ corresponds to the binding stoichiometry. With the immobilization procedure complete, non-specifically bound proteins were removed by washing with buffer HBS-EP+ (Cytiva) until the resonance signal reached a constant baseline. The reference flow cell provided a control surface for refractive index change and non-specific binding. DnaB solution was injected five times at increasing concentrations over the DciA-functionalized sensor surface in a single-cycle kinetics experiment. A long dissociation phase and a single regeneration step followed the last sample injection, without regeneration between each sample-injection.

The interaction between DnaB and DciA was assumed to display 1:1 kinetics; as such, the measured SPR data should display pseudo-first-order kinetics that is described by the following rate equation:

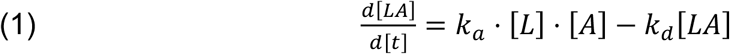

where k_a_ is the association rate constant, which refers to the rate of complex formation (units: M^-1^s^-1^); the dissociation rate constant k_d_ describes the stability of the complex, i.e., the fraction of complexes that dissociate in the time frame (units: M). [A] represents the analyte concentration; [LA] is the complex concentration measured directly as R in Resonance Units (RU); [L] is the ligand concentration on the surface.

To enable fitting of the SPR data, equation (1) was modified as follows:

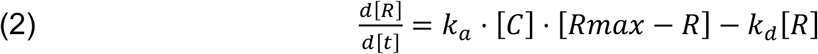

where Rmax is the maximum binding capacity, and Rmax-R is the free concentration of the analyte. Rmax and R were extracted from the experimental SPR curves. The fitted data proved estimates for k_d_ and k_a_ from which K_D_ = k_d_/k_a_ was calculated. Experimental binding kinetic data were fit to the ‘1:1 binding’ model implemented in the Biacore X100 Control Software (Cytiva).

### Native Mass Spectrometry

*Vc*DnaB and *Vc*DciA samples for native mass spectrometry (nMS) were exchanged into 500 mM ammonium acetate, pH 7.5, and 0.01% Tween-20 (nMS buffer) using a Zeba micro spin desalting column with a molecular cutoff of 40 kDa (ThermoFisher Scientific). Complexes containing DnaB and DciA in the above buffer were mixed in defined molar ratios before nMS measurements. To take data in the presence of nucleotide, ATP and magnesium acetate were added to final concentrations of 10 µM and 0.5 mM, respectively.

The list of DNA molecules used in the nMS study appears in Supplementary Table 2. Lyophilized synthesized ssDNAs (IDT) oligonucleotides were brought to 200 µM in HPLC-grade water and diluted to a working concentration of 3 - 6 µM in the above nMS buffer.

An aliquot (3 – 5 µL) of an experimental sample was loaded into a gold-coated quartz capillary tip prepared in-house. The sample was then electrosprayed into an Exactive Plus EMR instrument (ThermoFisher Scientific) using a modified static nanospray source (62, 63). Spectra were continuously recorded from a single sample-loaded capillary. For analysis of DnaB and DnaB-containing assemblies, the general MS parameters used include spray voltage, 1.2 – 1.3 kV; capillary temperature, 125 – 150 °C; S-lens RF level, 200; resolving power, 8,750 or 17,500 at *m/z* of 200; AGC target, 1 x 10^6^; number of micro scans, 5; maximum injection time, 200 ms; in-source dissociation (ISD), 0 – 10 V; injection flatapole, 8 V; interflatapole, 4 V; bent flatapole, 4 V; high energy collision dissociation (HCD), 200 V; ultrahigh vacuum pressure, 5 – 7 × 10^-10^ mbar; total number of scans, 100. For analysis of full-length *Vc*DciA, the MS parameters are similar as above except for capillary temperature, 100 °C in-source dissociation (ISD), 100 V; injection flatapole, 8 V; interflatapole, 7 V; bent flatapole, 6 V; higher-energy collisional dissociation (HCD), 0 V; ultrahigh vacuum pressure, 1.4 × 10^-10^ mbar. Mass calibration in positive EMR mode was performed using a cesium iodide solution. Raw nMS spectra were visualized using Thermo Xcalibur Qual Browser (version 4.2.47). Data processing and spectra deconvolution was performed using UniDec version 4.2.0 (64, 65) with the following general parameters: background subtraction (if applied), subtract curve, 10; smooth charge distribution, enabled; peak shape function, Gaussian; degree of Softmax distribution (beta parameter): 2 – 20. PDBO measured all mass spectrometric data in the laboratory of BTC.

### FRET-based DNA Strand Displacement Assay

DNA unwinding by *Vc*DnaB and its stimulation by *Vc*DciA were measured using a fluorescence resonance energy transfer (FRET) helicase strand displacement assay adapted from reference (20). The DNA substrates for the assay, FRET-DNA-fork-top-strand, FRET-DNA-fork-bottom-strand, and FRET-DNA-fork-capture-oligo, are identical to those described in reference (20). The top strand is labeled with Cy3 at its 5ʹ end; the bottom strand is labeled with the BHQ_2 quencher at its 3ʹ end. The capture oligo is unlabeled (Supplementary Table 2).

Strand displacement was measured by mixing annealed and fluorescence-quenched substrates at a concentration of 100 nM with 200 nM DnaB and DciA concentrations ranging from 0 nM to 6400 nM. An excess of the capture oligo (400 nM) was included to prevent the reannealing of the fork after strand displacement. FRET data were taken at 37 °C in the following buffer: 20 mM K-HEPES pH = 7.5, 5 mM Mg acetate, 50 mM K-glutamate, 5% glycerol, and 4 mM dithiothreitol (DTT). Reaction components were mixed into wells (volume: 100 µL/well) of a Corning black plate (p/n: 3880) with 96 half-area clear flat bottom wells. The working concentration of NaCl in the assay is 50 mM. The reaction was initiated by adding 1 mM ATP to each well; the plate was subsequently inserted in a SpectraMax M5 plate reader (Molecular Devices). FRET signals were recorded for 30 min at 37 °C. To avoid the crosstalk between Cy3ʹs closely spaced excitation and emission signals, an excitation wavelength of 520 nm was used; at this wavelength, excitation is reduced to 50% of its maximal value (https://www.thermofisher.com/us/en/home/life-science/cell-analysis/fluorophores/cy3-dye.html); however, this wavelength does not overlap with the fluorescence emission peak, measured at 570 nm. Relative Fluorescence Unit (RFU) data taken in this way were exported for analysis in Microsoft Excel and GraphPad Prism.

### FRET-based Replicative Helicase Loading Assay

To measure helicase loading by DciA, we adapted the unwinding assay described above; instead of a fork substrate with 5ʹ and 3ʹ unpaired segments, we employed substrates that featured single-stranded ‘bubbles’ (Supplementary Table 2) whose sequences are taken from (#894 and #895, reference: (42)). Bubble substrates favor DnaB assembly on ssDNA instead of threading the free end of ssDNA into the DnaB central chamber. In our first bubble substrate, the top strand was singly labeled at the 5ʹ end with Cy3, and the bottom strand with the Iowa Black RQ quencher (IDT). To account for the possibility that DnaB helicase could, in principle, be loaded for translocation in either direction, we employed a second substrate wherein fluorophore – quencher pairs were installed at both ends of the ‘bubble’ (Supplementary Information); such a substrate could inform on the loading and translocation on both strands and in both directions. As with the strand displacement assay, unlabeled capture strands forestalled the reannealing of the quencher-containing bubble strand.

FRET signals from helicase loading were recorded and analyzed as described for the strand displacement assay.

### Single Particle Cryogenic Electron Microscopy

Samples for cryo-EM were prepared and imaged, and the images were processed using standard techniques and software (66–69). Supplementary Methods, Supplementary Figures 3 and 4, and Supplementary Table 3 provide detailed descriptions.

### Model Building and Structure Refinement

The DeepEMhancer-sharpened map H (Supplementary Figure 3) was selected to build a molecular model of the *Vc*DnaB-ssDNA complex. Map H was submitted to ModelAngelo (70) to construct the initial protein and DNA model. The quality of the resulting model was high, even as it contained numerous chain breaks. Next, we superimposed the AlphaFold predicted structure of *Vc*DnaB, which has been split into three component domains (N-terminal Domain (NTD, residues 1:170), Linker Helix (LH, residues 171:211), and C-terminal Domain (CTD, residues 212:468)), onto the ModelAngelo output to create a model of hexameric *Vc*DnaB with excellent stereochemistry. The program COOT (71) was used to connect domains belonging to each monomer and to remove the following DnaB residues (chain A, B: 1-21 and 463:468, C-E-F: 1:21 and 464:468, D: 1:22 and 464:468) because of missing map density. Although *Vc*DciA was present in a four-fold excess in our cryo-EM sample, we found no EM density that the *Vc*DnaB model did not explain. As such, we conclude that *Vc*DciA is not present in our reconstruction.

The EM density in map H revealed an 11-mer ssDNA within the internal chamber of DnaB. This was sufficiently resolved to distinguish the DNA backbone from the nucleobases; however, it was not so resolved to permit assigning the nucleotide sequence to the map. In many places, purines could be distinguished from pyrimidines; however, we could not unambiguously position the 60-mer ssDNA in our cryo-EM sample. An 11-mer ssDNA molecule (TCCAGATACAC) was constructed in COOT (71, 72) and placed in density using its ‘real space refine’ function. In the lower resolution map E0 (6.24 Å, Supplementary Figure 3), we observed density extending away from DnaB along the axis where the 11-mer had been built. The density quality in this map did not permit the DNA backbone to be distinguished from the nucleobases. However, model building suggested that an additional 11 nucleobases could explain this density. Since this density provided no unique nucleobase information, we built this segment as polyadenine. The extra 11 DNA bases visible in Map E0 were refined in COOT against this map. The complete 22-mer ssDNA was refined against the high-resolution map H.

The *Vc*DnaB-ssDNA-ATPγS model was refined with Phenix (73–80). We used the ‘real_space_refine’ protocol, encompassing global minimization, morphing, NQH flips, and atomic displacement parameters (ADP). The refinement leveraged three rigid bodies (DnaB residues 1:171, 171:211, 212:468, 501 for each chain; ssDNA 1:11), Ramachandran, and secondary structure restraints. Refinement was carried out against the DeepEMhancer sharpened 3.37 Å density map H. A second round of real-space refinement encompassing global minimization, atomic displacement parameters, NQH flips, with Ramachandran and secondary structure restraints completed the model (Supplementary Table 3). The final model includes Chain A (DnaB residues 22:461), Chain B (DnaB residues 22:462), Chain D (DnaB residues 23:463), Chains C-E-F (DnaB residues 22:463), six ATPγS•Mg^2+^ complexes, one per chain (chains A-B-C-D-E-F residue 501), and ssDNA (chain G, residues 1-22).

### Model Analysis and Visualization

Structural analysis and visualization of the *Vc*DnaB-ssDNA complex were performed in the CCP4 software package (81), Coot (71, 72), the Uppsala software suite (82–85), UCSF-CHIMERA (86), Phenix (73–80), PyMOL (87), Nucplot (88), and as described previously (44). Distances and angles within and between protein structures were calculated using the PyMol Python scripts titled “pairwise_dist” (https://pymolwiki.org/index.php/Pairwise_distances) and “angle_between_domains” (https://pymolwiki.org/index.php/Angle_between_domains). Molecular graphics and figures were generated using UCSF-CHIMERA (86) and PYMOL (87). The software used in this study was sourced from the SBGrid Consortium (89, 90).

## Results

### Cryo-EM Structure of *Vc*DnaB Bound to ssDNA and ATPγS

To understand the loading of the DnaB helicase by DciA, we applied single-particle cryogenic electron microscopy (cryo-EM) to the *Vc*DnaB helicase bound to ATPγS, *Vc*DciA, and ssDNA derived from the *Vc*OriC-I replication origin. In assembling the complex, we relied on results from our nMS study (below). Cryo-EM analysis (Supplementary Methods, Supplementary Figures 3, 4, and Supplementary Table 3) produced a 3.37 Å structure of hexameric *Vc*DnaB bound to ATPγS and complexed to a 22-nucleotide ssDNA molecule (Figure 2 and Supplementary Figures 5, 6, 7, and 8). The resolution of the EM density corresponding to one of the chains (chain A) was lower than that of the other chains. Notwithstanding the fourfold excess in the cryo-EM sample, our maps displayed no density corresponding to the DciA loader. Our *Vc*DnaB-ssDNA complex likely represents the end-state structure after DciA-mediated loading or self-threading.

Compared to the *Bst*DnaB-ssDNA complex, determined with GDP•AlF_4_ (PDB: 4ESV (10)), the ATPγS form of the *Vc*DnaB-ssDNA complex presents a distinct configuration (Figure 3). Both share the trimer of dimers and pseudo-sixfold architecture of the NTD and CTD tiers. With diameters of ∼55 Å and ∼25 Å, the NTD and CTD layers in the *Vc* ATPγS form are dilated and constricted (8), respectively (Figures 2, 3, and Supplementary Figures 8 A-B-C-D). However, the six-chain NTD tier from the *Vc* ATPγS structure adopts a closed triangular planar and symmetric arrangement, resembling the NTD tier from other structures (PDB: 2R5U (91), Root Mean Square Deviation (RMSD) = 2.6 Å over 558 Cα atoms; PDB: 7T20 (92), RMSD = 1.5 Å over 825 Cα atoms). This configuration differs from the open spiral in the GDP•AlF_4_ form of the Bst-DnaB-ssDNA complex (10). As with other DnaB structures, the *Vc* NTD assembles around weak (head-to-head) and strong interfaces (tail-to-tail), defined based on the buried surface area (7); the NTD tier in the *Bst* structure is breached on the weak head-to-head interface. In both nucleotide forms, the constricted CTD tier is arranged as a closed spiral; however, the spirals are distinct (Supplementary Figure 9). The six DnaB subunits in the *Vc* and Bst structures are labeled A-F-E-D-C-B around the spiral; the breach in the CTD spiral lies between chains A and B, which are at the bottom and top of the spiral (as in the pose in Figures 2 and 3), respectively. Five of the six subunits of each structure (PDB: 4ESV: residues 181:441 and chain C, D, E, F, and A on *Vc*DnaB: residues 204:441 and chain B, C, D, E, and F) chains could be superimposed with an RMSD of 1.7 Å (1016 Cα atoms). The most significant difference in CTD configurations is the distinct position occupied by *Vc*DnaB chain A, which maps to chain B in the GDP•AlF4 4ESV structure; we address this change in greater detail below. We also note that in the *Vc* ATPγS form, chain A is located further away from its neighbors (distances to chain B: ∼39.6 Å, to chain F: ∼34.6 Å) than the average distance separating the remaining four subunits to their neighbors (Chains C, D, E, and F: ∼33.4 Å). By contrast, in the GDP•AlF4 4ESV structure, Chain A is distant from one immediate neighbor (distance between chains B and A: 42.6 Å), while other chains are 32.5 Å from their neighbors. Lastly, in the *Vc* ATPγS complex, the six Linker Helices (LH) occupy three positions relative to the NTD and CTD. The LH elements of chains B, C, D, and E occupy one position (LH1), with chains F (LH3) and A (LH2) diverging from this position by rotations/displacements of ∼16.5°/∼7.1Å and ∼29.6°/12.3 Å, respectively. In the GDP•AlF_4_ 4ESV complex, the LH elements occupy two positions, one by chains B, C, D, E, and F (LH1) and a second position (LH4) by chain A (Supplementary Figure 10).

**Figure 3.**
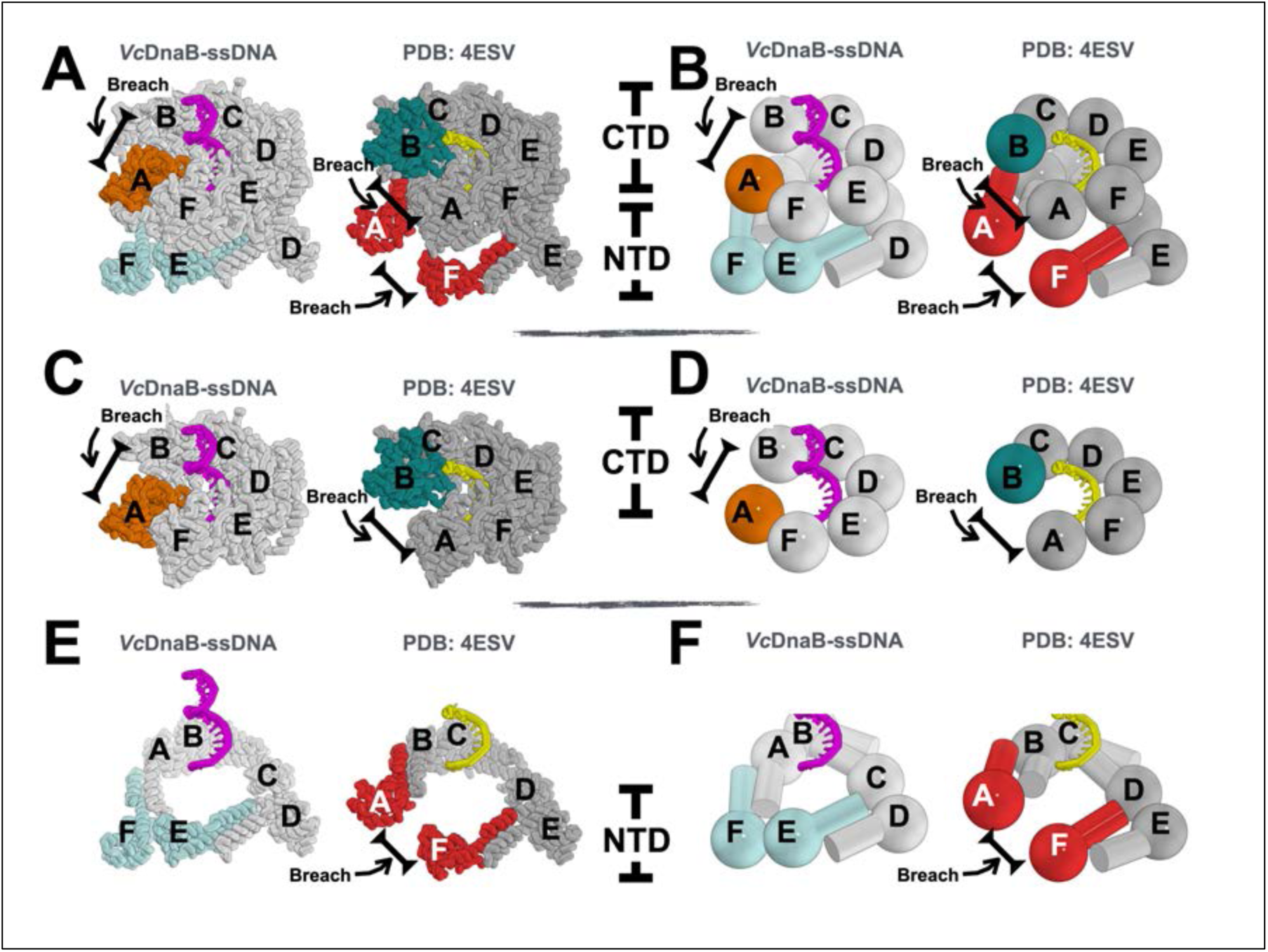
Nucleotide-Dependent Conformational Differences in the *Vc*DnaB-ssDNA-ATPγS and *Bst*GDP•AlF_4_ complexes. The two complexes are superimposed in all panels, as described in the main text. For clarity, the two entities have been translated with respect to each other, allowing each to be visible. A) Ribbon representation of both complexes. *Vc*DnaB NTD components chain F and E in the closed planar and triangular configuration are colored in pale cyan. *Bst*DnaB NTD components chain A and F in the open spiral configuration are colored in red. *Vc*DnaB chain A is colored orange. 4ESV chain B is colored deep teal. The ssDNA in the *Vc* and *Bst* complexes is depicted in both cartoon and stick representations, colored purple and yellow, respectively. All other chains of DnaB are colored in light gray (*Vc*) and dark gray (*Bst*). The breaches in the NTD and CTD layers are indicated. Chains corresponding to the N-terminal domain (NTD) and C-terminal domain (CTD) components are labeled. B) The depiction is identical to that in panel A, except that the NTD and CTD layers are represented using the sphere and mushroom-like shape design language. C) The exact representation as panel A, except that only the CTD layer is shown. D) Identical depiction as panel C except that the DnaB CTD layers are depicted as spheres. E) The exact representation as panel A, except that only the NTD layer is shown. F) Identical depiction as panel E except that the DnaB NTD layers are depicted using mushroom-like shapes.

ssDNA in the complex traces a spiral path that passes through the internal chamber of *Vc*DnaB (Figure 2). 11 of the 22 ssDNA nucleotides are bound to subunits of the *Vc*DnaB hexamer; the remaining 11 are outside the DnaB perimeter (Supplementary Figures 7 and 11). In contrast to the GDP•AlF_4_ 4ESV structure, where each subunit contacts ssDNA, only five DnaB subunits in the current *Vc* structure make contacts (less than 4.0 Å); chain A does not contact ssDNA. An inventory of protein-DNA contacts reveals the involvement of seven *Vc*DnaB residues, though not every contact is made by each subunit (Supplementary Table 4).

Six ATPγS/Mg^2+^ complexes are found at composite catalytic centers formed between adjacent *Vc*DnaB subunits, wherein the Walker A/P-loop and Walker B motifs are provided by one subunit and the β-hairpin arginine finger by a second (Figure 2C and Supplementary Figures 6 and 8E). In four nucleotide sites (chain F-E, E-D, D-C, and C-B), the β-hairpin/arginine finger makes a close approach to nucleotide on a neighboring subunit, with distances between the P-loop lysine (K234) and the arginine (R439) finger residues of ∼11.3 Å. The corresponding distance at the A-F subunit interface is slightly longer (∼11.9 Å), and the arginine finger has shifted by ∼3 Å relative to other subunits. The β-hairpin/arginine finger at the nucleotide site that lines the breach (B-A) exhibits a considerably longer (∼24.5 Å) distance. In the *Bst* GDP•AlF_4_ complex, 5/6 sites (Chains A-F, F-E, E-D, D-C, and C-B) feature an average P-loop lysine (K216) to arginine (R420) finger distance of ∼11.4; the chain B-A site, which lines the breach, has a considerably longer distance at ∼32.8 Å.

Taken together, the significant changes between the GDP•AlF_4_ and ATPγS forms of DnaB, including distinct NTD configurations, the offset of a single DnaB CTD, and diversity in LH element conformations in sequential subunits around the ring, imply that our structure has captured a distinct intermediate in the translocation reaction.

### Remodeling of the NTD and CTD Tiers Accompanying Translocation

To place the distinct ATPγS form of DnaB in the translocation reaction, we compared it to the *Bst* GDP•AlF_4_ (10) entity (Figure 3) and a series of DNA complexes (PDB: 6N7I, 6N7S, 6N7T, and 6N7V) of the DnaB-family helicase Gp4 from phage T7 (24, 29) (Comparisons to other *Vc*DnaB structures appear in the Supplementary Information). Aligning the *Vc* ATPγS and *Bst* GDP•AlF_4_ structures by superimposing 5/6 CTDs (RMSD: 1.7 Å, a similar alignment could be obtained by aligning on the ssDNA (RMSD: 0.64 Å) reveals that *Vc* chain A, excluded from the alignment, is displaced by 25 Å and rotated by ∼22° relative to the corresponding subunit in 4ESV (Chain B). Similar displacements/rotations are seen in the HelF subunits in the 6N7T and 6N7V DnaB-family Gp4 helicase structures, shifting positions by ∼24 Å and ∼13° (24, 29). Further, the DnaB alignment juxtaposes all the components of the NTD layer save for the chains at the top (4ESV: chain A) and bottom (4ESV: chain F) of the *Bst* spiral. These two chains are offset owing to the distinct planar vs. spiral configurations adopted by the NTD tier of the *Vc* ATPγS and *Bst* GDP•AlF_4_ complexes. Unexpectedly, the breaches in the two structures do not align (Figure 3). Instead, they are offset by one subunit.

Given the distinct states captured by each structure, we reasoned that the *Vc* ATPγS structure might precede the transition state *Bst* complex in the DNA translocation reaction coordinate. Ordering the structures in this way specifies a direction for the observed structural changes. Accordingly, the NTD of chain F and chain E in the *Vc* ATPγS complex will transition from the closed triangular planar configuration to the open spiral of the GDP•AlF_4_ complex (mapped to 4ESV chains: A and F); concomitantly, the CTD of *Vc* chain A will move upward (as in the pose in Figure 3) to assume the configuration seen in chain B of 4ESV (Supplementary Movie 1). With the migration of the *Vc* chain A CTD to the position specified in the 4ESV structure, the breach in the CTD spiral moves counterclockwise by one subunit. The LH element moves from the LH3 to the LH4 position in 4ESV chain A. Several lines of evidence in the *Vc* ATPγS complex support the proposal that the chain A CTD migrates (Figure 3). First, the arginine finger/β-hairpin element on chain F has peeled away by ∼3 Å, relative to its position in chains B, C, D, and E, from the P-loop on chain A (Figure 2C). Second, chain A is farther away from its neighbors than any other DnaB subunit. Third, chain A makes no contact with ssDNA. Lastly, of all the CTDs, the EM density for chain A is the weakest. Together, these findings point to the disengagement of the chain A CTD from its CTD neighbors and ssDNA as it poises to transition. Of course, the chain A CTD remains linked to the DnaB hexamer through its NTD.

Due to the NTD layer’s asymmetry, the next subunit’s motion, chain F, must follow a different trajectory. We propose that on ATP hydrolysis, chain F, now located at the bottom of the spiral, will also peel away from DNA and migrate upwards to a new binding site on ssDNA. In contrast to the motion of chain A, which results in the opening of the NTD layer, the motion of chain F CTD must be accompanied by the closure of the NTD ring from the open spiral to the closed planar configuration; this motion will also result in the adjustment of the position of the LH element, from the LH3 position to the LH1. Together, these two changes re-establish the NTD configuration in the *Vc* ATPγS structure. Applying the above to chains E and D first and then to chains C and B allows each DnaB subunit to complete a catalytic cycle of ATP hydrolysis amid translocation on ssDNA. These findings highlight the importance of opening and closing the NTD layer, which had previously been suggested to specify the direction of unwinding (93) in the translocation reaction.

### Oligomeric State and Stoichiometry of *Vc*DnaB, DciA, ssDNA complexes

Next, we used high-resolution native mass spectrometry (nMS) to determine the oligomeric states and the stoichiometry of complexes in the DnaB-DciA system assembled from purified proteins (Supplementary Figure 2). We first investigated the oligomeric states of DnaB complexes from *Vc*. To provide context, the hexameric *E. coli* (*Ec*) DnaB (22) and monomeric *B. subtilis* (*Bst*) DnaB-family species (34) were included. nMS analysis showed that the *Vc* DnaB helicase is hexameric (Figure 4A and Supplementary Table 5). We observed a sizeable population of lower oligomeric states for *Ec* and *Vc*DnaB (DnaB_5_, DnaB_4_, DnaB_3_, and DnaB_2_) in samples without Mg•ATP. The addition of a small amount of ATP (10 µM ATP for a 10 µM DnaB monomer sample) before nMS analysis yielded a predominant peak for the hexameric DnaB with minimal subcomplexes (Figure 4A and Supplementary Table 5), indicating that ATP stabilizes the *Vc*DnaB hexamer as previously suggested for the *E. coli* ortholog (94, 95). All subsequent experiments with DnaB-containing samples were supplemented with 10 µM ATP and 0.5 mM magnesium acetate before nMS analysis. Close-up inspection of the nMS spectrum for the *Vc*DnaB sample incubated with Mg•ATP showed a peak distribution corresponding to several Mg•ADP-bound states, indicating ATP binding and hydrolysis (Figure 4B and Supplementary Table 5).

**Figure 4.**
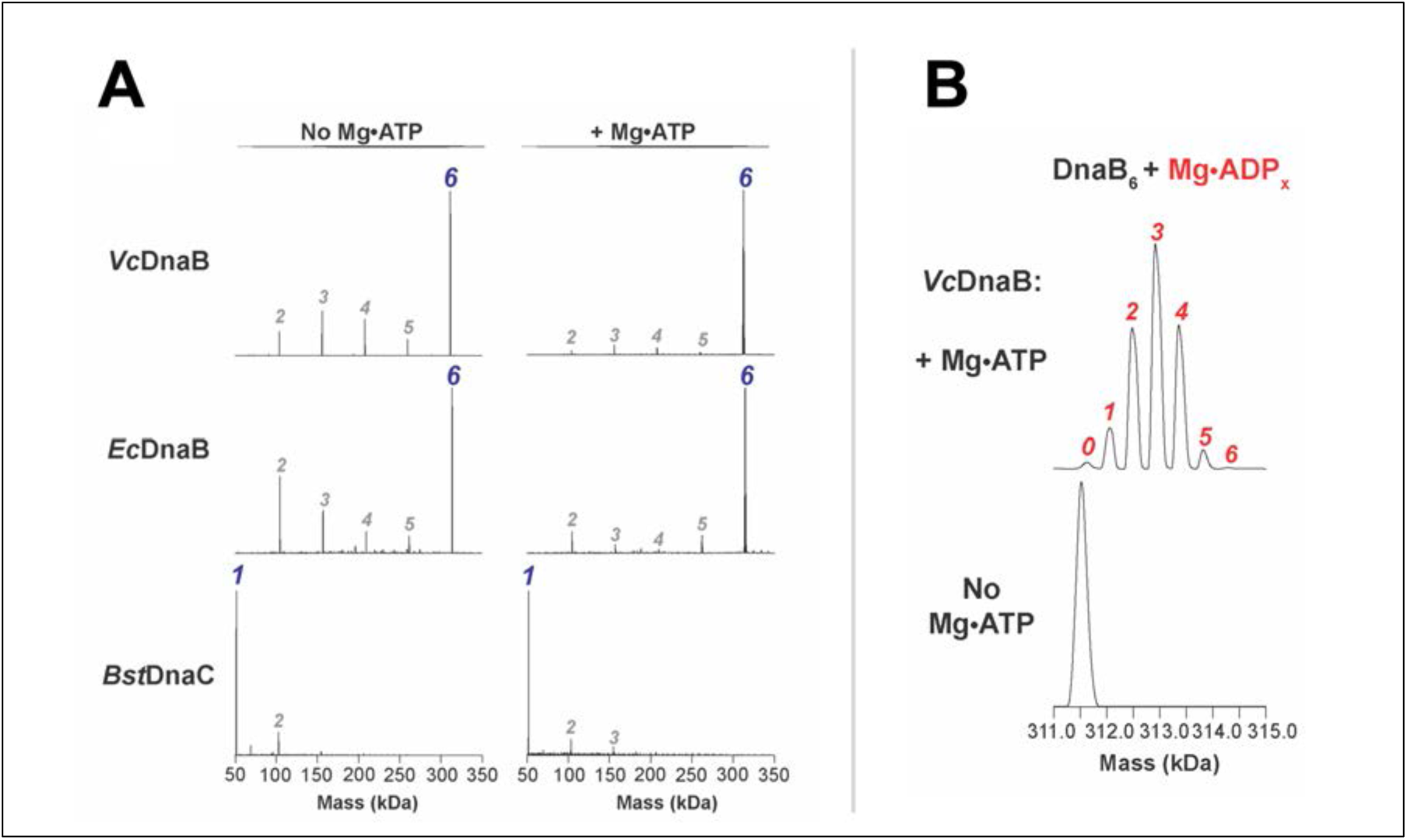
Although highly conserved, certain DnaB helicase orthologs feature distinct oligomeric states. (A) Deconvolved nMS spectra of replicative helicases from *V. cholerae* (*Vc*), *E. coli* (Ec), and *B. subtilis* (*Bst*) with and without Mg•ATP. Unlike the *B. subtilis* ortholog, which is monomeric (with or without ATP), *V. cholerae* and *E. coli* DnaB are hexamers. Each measurement was performed with a monomeric protein concentration of 10 µM. The *Bst*DnaC sample refers to the DnaB family helicase in *B. subtilis*. (B) Close-up of panel A peaks corresponding to the hexameric *Vc*DnaB complex. The average mass difference between adjacent peaks is 444 ± 20 Da; this value more closely matches the mass of ADP (Mass = 427 Da) than ATP (Mass = 507 Da). As such, we suggest that ATP hydrolysis has occurred, resulting in distinct Mg•ADP-occupied states. The numbers in panels A and B refer to the oligomeric state of *Vc*DnaB (A, blue) and of Mg-ADP molecules bound to *Vc*DnaB (B, red). A list of mass assignments of the various nucleotide-bound peaks in panels A and B appears in Supplementary Table 5.

Next, we examined complexes between *Vc*DnaB, *Vc*DciA, and ssDNA derived from the OriC-I chromosomal origin. Our nMS analysis showed that *Vc*DciA is monomeric (Supplementary Figure 12). Incubation of 10 µM (as monomer) *Vc*DnaB with 7 µM full-length *Vc*DciA, in the absence of ssDNA, revealed that the DnaB hexamer bound 1 – 2 copies of DciA, with small amounts of a third copy (Figure 5A). We attempted to analyze samples with higher relative amounts of DciA (> 10:7 DnaB: DciA) but encountered significant disassembly of *Vc*DnaB complexes with *Vc*DciA upon initiation of electrospray during nMS analysis (Supplementary Figure 13). Stoichiometries of 6:3 or 6:6 for the *Vc*DnaB-*Vc*DciA complex have been estimated from small-angle X-ray scattering and the X-ray crystal structure (12, 14).

**Figure 5.**
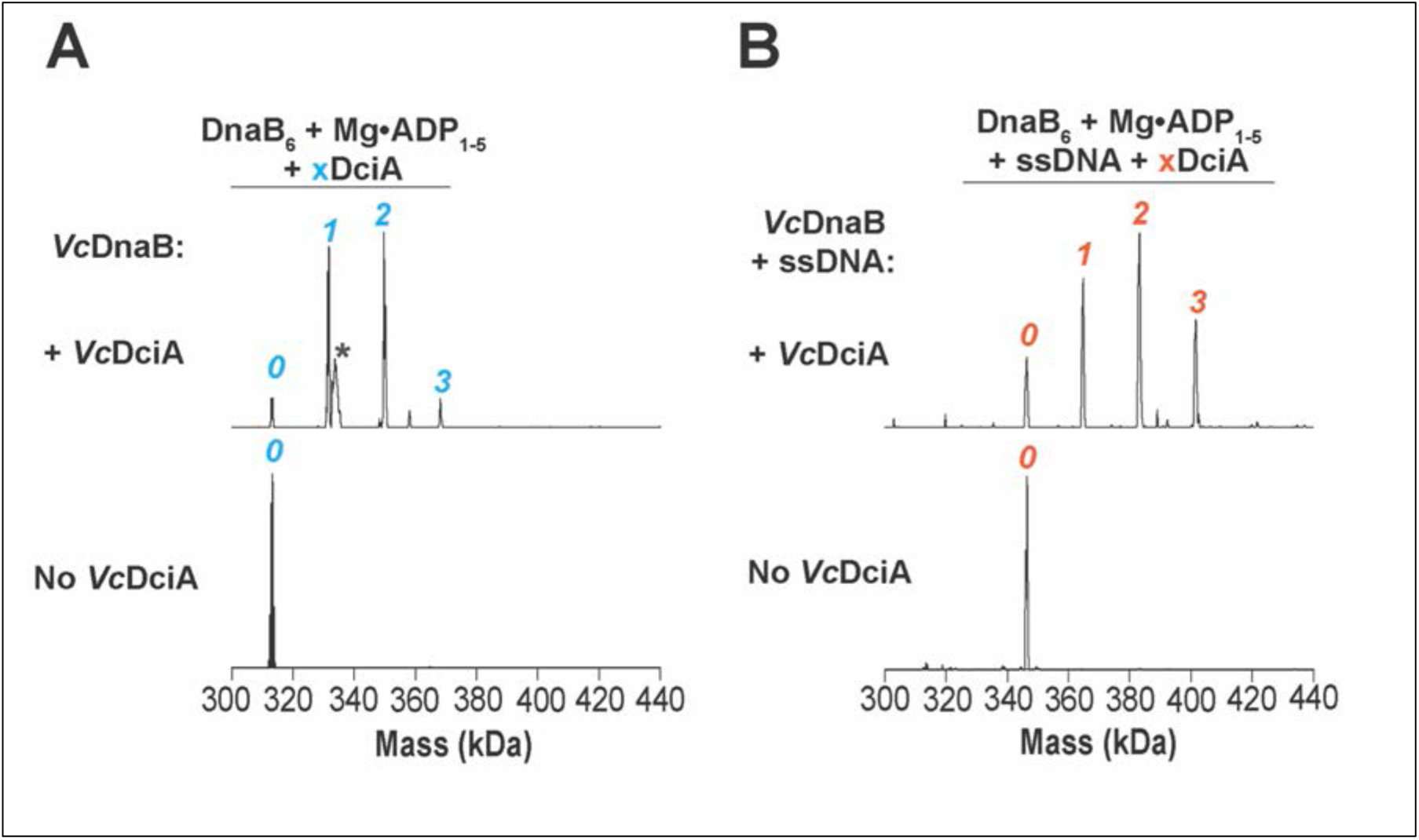
Up to Three Copies of *Vc*DciA Bind to Isolated Hexameric *Vc*DnaB and *Vc*DnaB Complexed to Replication Origin-Derived ssDNA. Deconvolved nMS spectra of 10 µM *Vc*DnaB (measured as the monomer) without and with 7 µM full-length *Vc*DciA (A) and in the absence or presence of 3 µM origin-derived ssDNA (O4-*Vc*-F) (B). The number of *Vc*DciA molecules observed bound to *Vc*DnaB is indicated in blue (A) or red (B). The expected masses include DnaB_6_ = 311.5 kDa, DciA = 18.4 kDa, and ssDNA = 32.5 kDa. The broad peak (*) is from an unknown low-level protein contaminant introduced during or after buffer exchange for this sample. All samples were exchanged into 10 µM ATP, 500 µM magnesium acetate, 500 mM ammonium acetate, and 0.01% Tween-20 before nMS analysis.

DnaB accepts ssDNA into its central chamber, the basis for the fork DNA unwinding assay(12, 42, 96). Accordingly, we found that *Vc*DnaB alone could bind OriC-I-derived ssDNA (Figure 5B). An nMS analysis of a mixture comprising *Vc*DnaB, *Vc*DciA, and the ssDNA showed that up to three copies of *Vc*DciA could be associated with ssDNA-bound DnaB. Interestingly, there is a marked increase in the relative peak intensity for the 6:3 DnaB: DciA assembly when ssDNA is bound compared to the ssDNA-free sample, suggesting that ssDNA association promotes the formation or stabilization of this higher-order ensemble.

We also investigated the binding of specific *Vc*DciA domains to *Vc*DnaB. nMS analysis of *Vc*DnaB titrated with increasing concentrations of the lasso domain of DciA showed a progressive increase in the number bound to the DnaB hexamer, with 1 – 4 copies bound when both proteins are present at equimolar amounts, to 2 – 6 copies bound at four-fold excess of DciA-Lasso (Supplementary Figure 14A). In contrast, only one copy of the DciA KH domain was found associated with DnaB_6_ at ∼10% relative peak intensity even at six-fold molar excess, indicating that the KH domain of DciA weakly binds to DnaB (Supplementary Figure 14B). Incubation of *Vc*DnaB with the two separate DciA domains at the highest concentrations studied did not change the maximum number of bound copies observed (i.e., one and six for the DciA KH and Lasso domains, respectively, Supplementary Figure 14B). Contrary to what was observed with DnaB and full-length DciA, no DnaB helicase disassembly was observed during the electrospray nMS analysis when either or both DciA domains were used in excess. Moreover, the distribution of DnaB: DciA-Lasso assemblies did not change (1 – 4 copies of DciA-Lasso remained bound to DnaB_6_ at equimolar amounts of both proteins) in the absence or presence of ssDNA (Supplementary Figure 15).

Our nMS experiments revealed the stoichiometry of the isolated *Vc*DnaB helicase and its complex with DciA and OriC-I-derived ssDNA; both estimates are crucial to deciphering the loading mechanism. The hexameric nature of *Vc*DnaB implies that it is a closed protein ring and that the DciA helicase loader is likely a ring-breaker. The 6:3:1 stoichiometry of the *Vc*DnaB-DciA-ssDNA complex suggests that although six DciA binding sites may exist on DnaB, only three are used; this value will be crucial to understanding how the DnaB-DciA complex will interact with other factors during replication initiation.

### *Vc*DciA Binds to Cognate DnaB Helicases with High Affinity

Next, we used orthogonal approaches to show that DciA binds to cognate DnaB and that DciA likely binds near the DH—LH interface (Figure 6). First, we conducted NiNTA ‘pull-down’ experiments to examine the binding between full-length and truncated constructs of DciA and DnaB (Figures 6A and Supplementary Figure 16). We find that full-length *Vc*DciA binds to full-length *Vc*DnaB. Moreover, DnaB constructs that span the LH and CTD domains also retain binding to full-length DciA (a shorter construct spanning the CTD could not be expressed). No other DnaB construct (NTD or NTD-LH) showed binding to full-length DciA. Parallel experiments with constituent KH (*Vc*: 1:98) and Lasso domains (*Vc*: 98:157) revealed binding to full-length cognate DnaB, although the *Vc*DciA KH domain bound weakly. No other truncated DnaB construct (NTD or NTD-LH) showed any interaction with either the KH or Lasso domains. These data point to the LH interface as a strong candidate for part of the DciA binding site, with another part comprised of the CTD itself.

**Figure 6.**
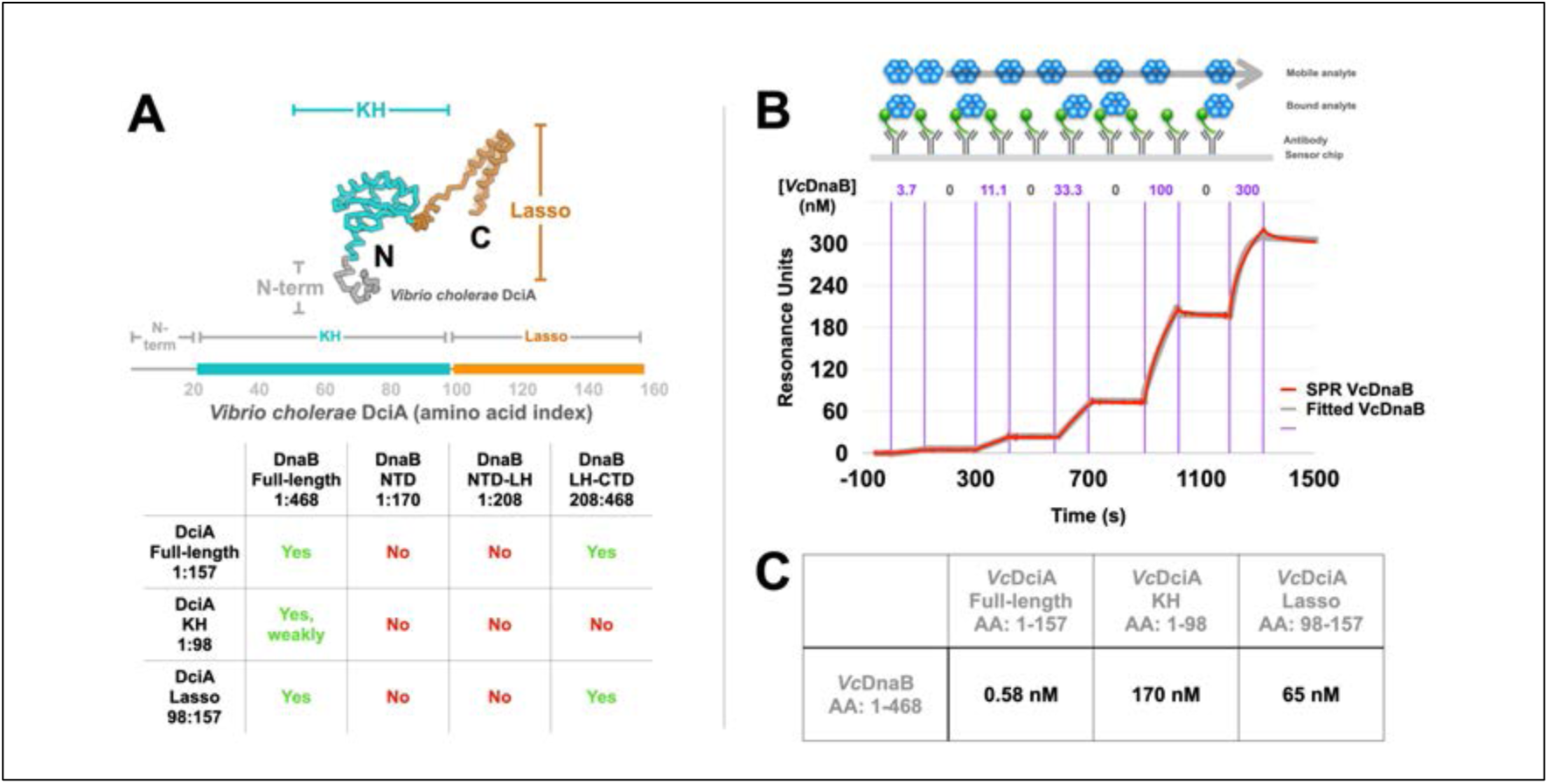
The *Vc*DciA loader Binds Tightly to *Vc*DnaB helicase. A) The primary sequence of *Vc*DciA is arranged as a bar, and the locations of the KH and Lasso domains are indicated. The table summarizes the NiNTA ‘pull-down’ experiments, wherein the green ‘yes’ and red ‘no’ indicate that the pull-down assay was scored to report binding or no binding, respectively (Supplementary Figure 15). B) The dissociation constants (K_D_) between DnaB and DciA were evaluated through single-cycle kinetic analyses *via* Surface Plasmon Resonance (SPR). On and off rates of full-length DnaB at incremental concentrations (3.7, 11.1, 33.3, 100, 300 nM) were measured by immobilizing cognate C-term His-tagged full-length *Vc*DciA (C; K_D_ = 0.59 nM; Chi^2^ = 4.33; R_max_ = 312.3) on the SPR sensor chip. A schematic of the SPR experiment accompanies the raw (red) and fitted ( grey) SPR curves. Experimental data were analyzed as described in the methods. The same approach was applied to measure K_D_s for the KH and Lasso constructs (Supplementary Figure 17). C) The K_D_ values emerging from the SPR analysis are summarized for the *Vc* system in tabular form.

Next, we applied surface plasmon resonance (SPR) to quantify the interaction between *Vc*DnaB and *Vc*DciA. We immobilized full-length DciA, the isolated KH, and Lasso domains to the SPR chip, measured on and off rates for full-length DnaB, and extracted dissociation constants (Figures 6B, 6C, and Supplementary Figure 17). As our pull-down binding experiments anticipated, *Vc*DnaB bound tightly (K_D_ = 0.58 nM) to full-length *Vc*DciA. We found the isolated *Vc* Lasso domain bound to DnaB with a K_D_ = 65 nM, but the KH domain exhibited a weak affinity (K_D_ = 170 nM) (Supplementary Figure 17).

Our data reveal that the most significant contacts between DciA and DnaB are found at the LH and CTD domains of DnaB, with the Lasso domain being the locus of many of these contacts. Given that two other loaders, *E. coli* DnaC and phage λP, bind at the DH-LH interface of two adjacent DnaB subunits (19, 43, 44, 48), the finding that the LH constitutes the DciA binding establishes this site as a conserved location of significance for DnaB helicase loading.

### Suppression of DnaB’s ATPase Activity by DciA

One of the hallmarks of a bacterial helicase loader is the suppression of ATPase activity of the cognate helicase (43, 46, 60). To evaluate the effect of *Vc*DciA on the ATPase activity of *Vc*DnaB, we measured the influence of full-length and truncated constructs of DciA against control measurements encompassing wild-type and the Walker B mutant E259A (Figure 7A). Our preparations of full-length and truncated DciA lacked ATPase activity. Titration of full-length *Vc*DciA revealed potent suppression of *Vc*DnaB’s ATPase activity (untagged DnaB: IC_50_ ≈ 26 nM, NHis-DnaB: IC_50_ ≈ 26 nM, Figure 7B). A construct encompassing the *Vc* KH domain (1:98) exhibited an IC_50_ ≈ 443 nM (Supplementary Figure 18A); the lower inhibitory constant is congruent with the weak interaction signaled by our nMS study and the NiNTA-based binding studies. Even though we found that the *Vc* Lasso domain (98:157) binds to full-length *Vc*DnaB, titration revealed no effect on *Vc*DnaB’s ATPase activity at low DciA concentrations (25 – 400 nM). However, at very high concentrations (800 – 6400 nM) of *Vc*DciA, we observed reproducible stimulation (up to 2.8 x at 4800 nM) of DnaB’s ATPase activity (Supplementary Figure 18B).

**Figure 7.**
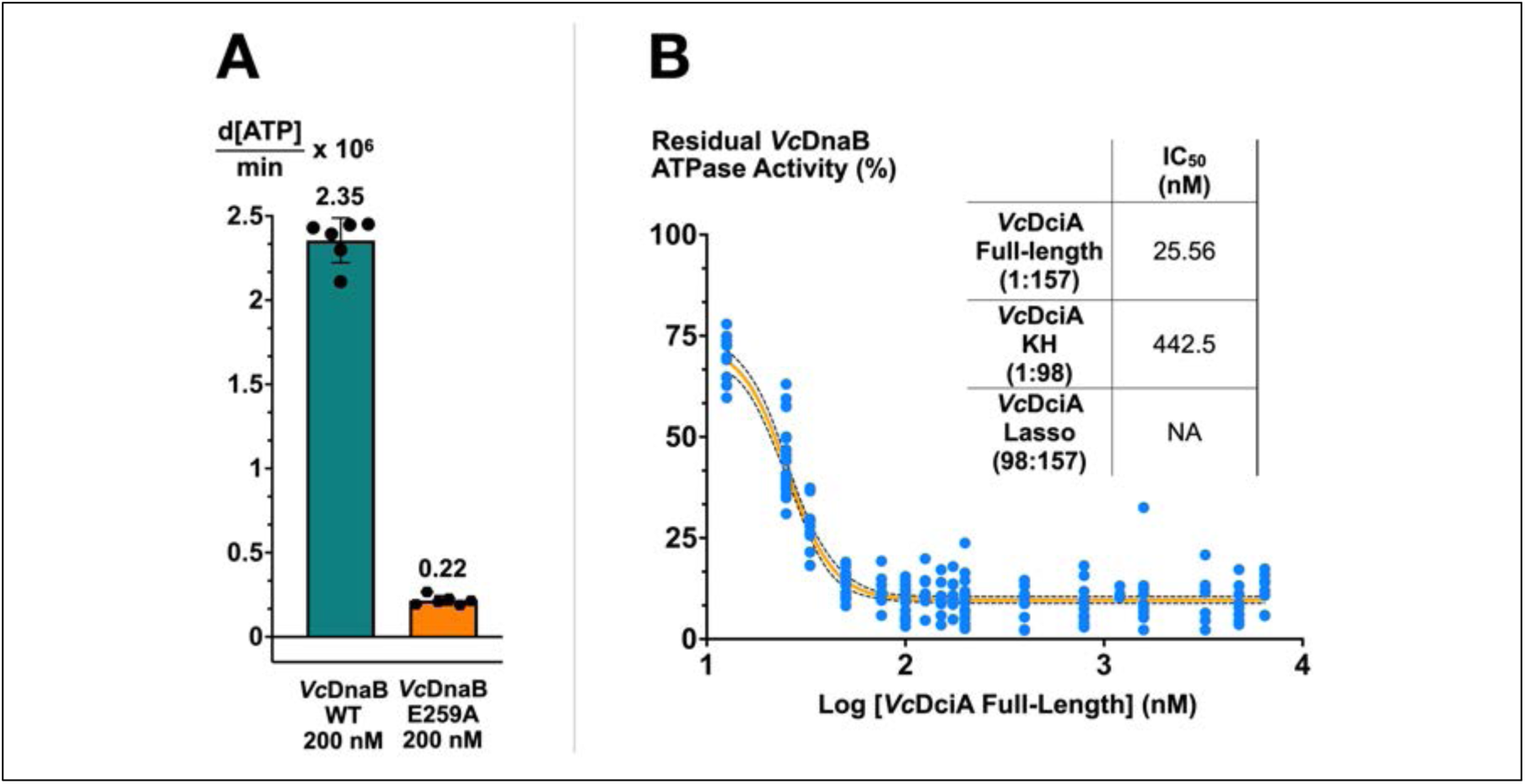
The full-length DciA loader suppresses VcDnaB’s ATPase activity. A) ATPase rate of 200 nM wild type (teal) and E259A mutant *Vc*DnaB (orange). The points represent the measured rates; the height of the bar represents the average of the measured points; the vertical bracket visually represents one standard deviation. Each experiment in panel A was performed two times in triplicate. B)The effect of titrating *Vc*DciA (0 - 6400 nM, log scale) on the residual ATPase activity of *Vc*DnaB (200 nM) is plotted. For each experiment, the residual activity is the quotient of the ATPase rate of a particular DciA concentration and the rate for 0 nM DciA. The blue points represent seven experiments with different batches of *Vc*DnaB and *Vc*DciA measured in triplicate on various days. The light-orange line represents the non-linear fit of the residual activity points (Sigmoidal, 4PL, X is log (concentration) model in Prism. The dotted lines represent the 95% confidence interval. The fit encompassed 210 points, yielded an R^²^ value of 0.91, and resulted in an IC₅₀ value of 25.56 nM (95% CI = 23.74 to 26.94 nM). Parallel analysis with the KH domain on four different batches revealed a 16-fold higher IC_50_ of 442.5 nM (R^²^ = 0.9693; 95% CI: 380.5 to 520.0 nM). The Lasso domain was not found to inhibit; however, we observed activation of DnaB’s ATPase at very high Lasso concentrations. The inset table summarizes the IC_50_ results. Control measurements with the E259A *Vc*DnaB Walker B mutant exhibited no ATPase activity (panel B, orange).

### DciA Loads DnaB on Physiological DNA Bubble Substrates

The nMS binding data and DNA unwinding experiments (Supplementary Information and Supplementary Figures 19, 20, 21, and 22) implicate DciA as a helicase loader since DciA enhances both underlying activities. However, the substrates employed in the above assays (ssDNA, 5ʹ and 3ʹ ssDNA tailed fork) do not permit self-threading into DnaB’s central chamber to be distinguished from loading/assembly around ssDNA. Physiologically melted DNA by DnaA at a replication origin has no free termini, and, as such, assembly around ssDNA, rather than threading, is expected to be the predominant form of loading *in vivo*.

To distinguish the DciA-mediated assembly of *Vc*DnaB on ssDNA from self-threading, we measured unwinding from a DNA bubble substrate, which features no free ends, unlike a tailed-fork substrate. Our assay employed DNA molecules #894 and #895 previously described (42), which we converted into a FRET-based readout by adding a Cy3 fluorophore at one 5ʹ end and the Iowa Black RQ quencher at the corresponding 3ʹ end (Figure 8). We also measured DnaB loading onto a doubly labeled DNA bubble (Supplementary Information and Figures 23, 24, 25C, and 25D).

**Figure 8.**
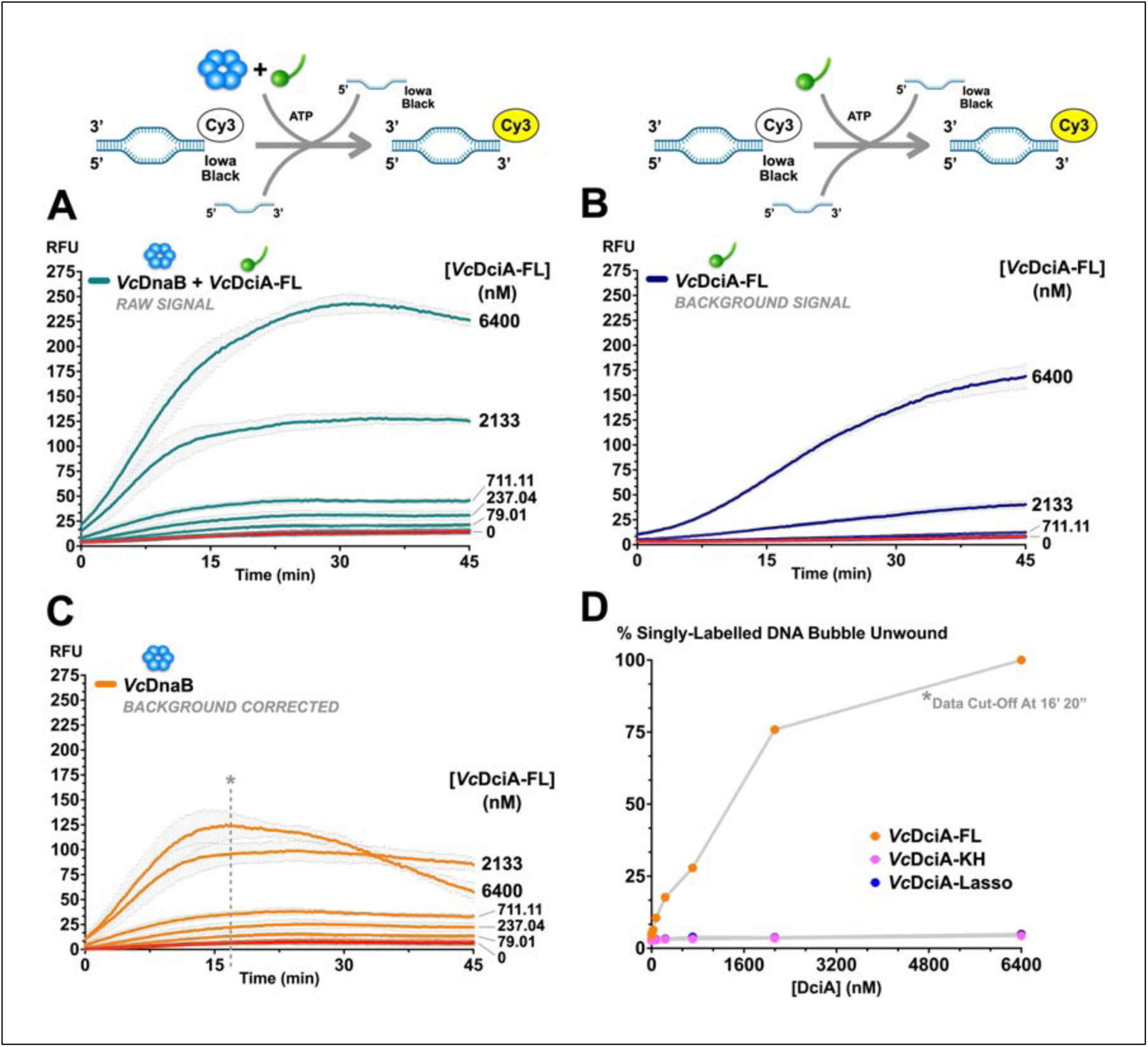
*Vc*DnaB is loaded onto a physiological DNA bubble substrate by the full-length *Vc*DciA loader. Helicase loading was measured using a DNA substrate that mimics unwound DNA at a replication origin, as is diagrammed in the schematic above. A) Loading of full-length *Vc*DnaB by full-length *Vc*DciA. Loading is read out by measuring fluorescence (in relative fluorescence units, RFU) over 45 minutes at 570 nm from a DNA substrate after it had been unwound by DciA-loaded DnaB; loading relieves the quenching to provide a signal. Data were taken by titrating *Vc*DciA-FL (0.11-6400 nM) against *Vc*DnaB (200 nM). Each labeled curve represents the average of 6 independent measurements taken from different batches of proteins on different days. The lack of loading activity in the curve that omits *Vc*DciA (red curve) implies that DnaB’s self-threading activity has been suppressed with the replication-origin mimicking DNA bubble substrate. The standard deviation of each point is plotted in light gray. B) Same as Panel A except that *Vc*DnaB is omitted. Envisioned as a control, this measurement teaches that *Vc*DciA appears to have the capacity to unwind bubble DNA on its own. C) Point-by-point subtraction of the curves in panels A and B estimates the bubble unwinding signal that arises solely from *Vc*DnaB. The downward phase of the 6400 nM DciA curve after 16 minutes arises from unwinding rate differences between *Vc*DnaB and *Vc*DciA. These data suggest that *Vc*DciA stimulates the loading of *Vc*DnaB by ∼25-fold, ranging from 5 to 125 RFU. D) The unwinding data in panel C is converted to a percentage unwound by dividing by the maximum RFU point and plotting the resulting values against *Vc*DciA concentration. Parallel measurements reveal that the isolated *Vc*DciA KH and Lasso domains (Supplementary Figure 25A and 25B) exhibit little (less than 5%) to no loading activity.

No fluorescence signal was observed when DciA was omitted (Figure 8, red curve), leading us to conclude that self-threading of ssDNA into DnaB had been suppressed with the bubble substrate. Titration of full-length *Vc*DciA (0.11 to 6400 nM) revealed a 10-fold (∼15 to ∼150) increase in fluorescence at the highest concentration (6400 nM). The increase in fluorescence strictly depended on ATP since the *Vc*DnaB E259A mutant showed no activity (Supplementary Figure 24). Substitution of the two DciA truncations – the NTD KH and the CTD Lasso – for full-length *Vc*DciA revealed essentially no loading activity (less than 5%) (Supplementary Figures 25A and 25B).

To better understand *Vc*DciA’s loading activity, we repeated the above titration without *Vc*DnaB. Surprisingly, titration of DciA against the DNA bubble substrate revealed a substantial fluorescence signal (Figure 8B); these data imply that *Vc*DciA has a previously undiscovered activity: to unwind dsDNA on its own. This activity did not require ATP. The finding that isolated *Vc*DciA produced a fluorescence signal ∼66% of that seen in the *Vc*DnaB-DciA titration implies significance, though establishing biological relevance will require further experimentation. DciA’s apparent unwinding activity is not observed when the DNA bubble is substituted with the fork substrate (Supplementary Figure 22), indicating a role for DNA structure in this activity.

We subtracted the two signals to separate the unwinding activities of *Vc*DnaB-DciA and isolated *Vc*DciA (Figure 8C). The calculated signal from *Vc*DnaB, when acted upon by *Vc*DciA, showed a 25-fold (∼5 to ∼125 RFU, Figure 8C) increase in fluorescence at the highest concentration (6400 nM) (Figure 8D). These data strongly argue that *Vc*DciA is a helicase loader for *Vc*DnaB.

The congruence of the DNA unwinding and helicase loading curves during titration with full-length *Vc*DciA suggests that, to a large degree, helicase loading rather than threading occurs with full-length DciA in both assays (Figure 8D and Supplementary Figure 19B). On the other hand, dissimilarity in the curves corresponding to the *Vc*DciA truncations implies that loading is not retained in either construct or that the observed DNA unwinding activity in the truncated constructs may arise solely from self-threading. Our data establish a helicase-loading biochemical activity for the *Vc*DciA protein. Moreover, the finding that loading onto the bubble substrate does not occur without DciA supports a closed protein ring configuration for DnaB, which agrees with our nMS results (Figure 4A).

## Discussion

Our analyses of the bacterial DciA loader and the DnaB helicase from *Vibrio cholerae* shed new light on the underlying helicase-loading and activation pathway (Figure 9). In bacteria, two copies of the DnaB replicative helicase are assembled onto single-stranded DNA introduced into the replication origin by the DnaA initiator protein. It is now clear that most bacteria lack an ortholog for the well-studied DnaC helicase loader; instead, the DciA helicase loader appears likely to be the predominant loading factor (37, 40, 41). *Vc* binds to cognate hexameric DnaB with a low nM K_D_. The unstructured Lasso segment mediates significant binding energy to the interaction with DnaB, and an essential part of the DciA binding site lies at the nexus of DnaB’s DH and LH elements. Prior studies suggested stoichiometries of 6:3 or 6:6 for the *Vc*DnaB-*Vc*DciA complex and a binding site at the DH-LH nexus (12, 14). We find up to three copies of *Vc*DciA and one copy of replication-origin-derived ssDNA bound to *Vc*DnaB. The stoichiometry of the DnaB-DciA system distinguishes it from the *E. coli* DnaB-DnaC and DnaB•λP complexes, which feature 6:6 and 6:5 stoichiometries, respectively (19, 43, 44, 48). The unwinding activity of DnaB is significantly stimulated by DciA on fork and replication origin-mimicking DNA bubble substrates. The finding that isolated *Vc*DnaB is a hexamer under physiological concentrations of ATP (Figure 4A) and exhibits essentially no self-loading on the replication-origin mimicking DNA bubble substrate (Figure 8) leads us to conclude that DciA is a ring-opening-type helicase loader.

**Figure 9.**
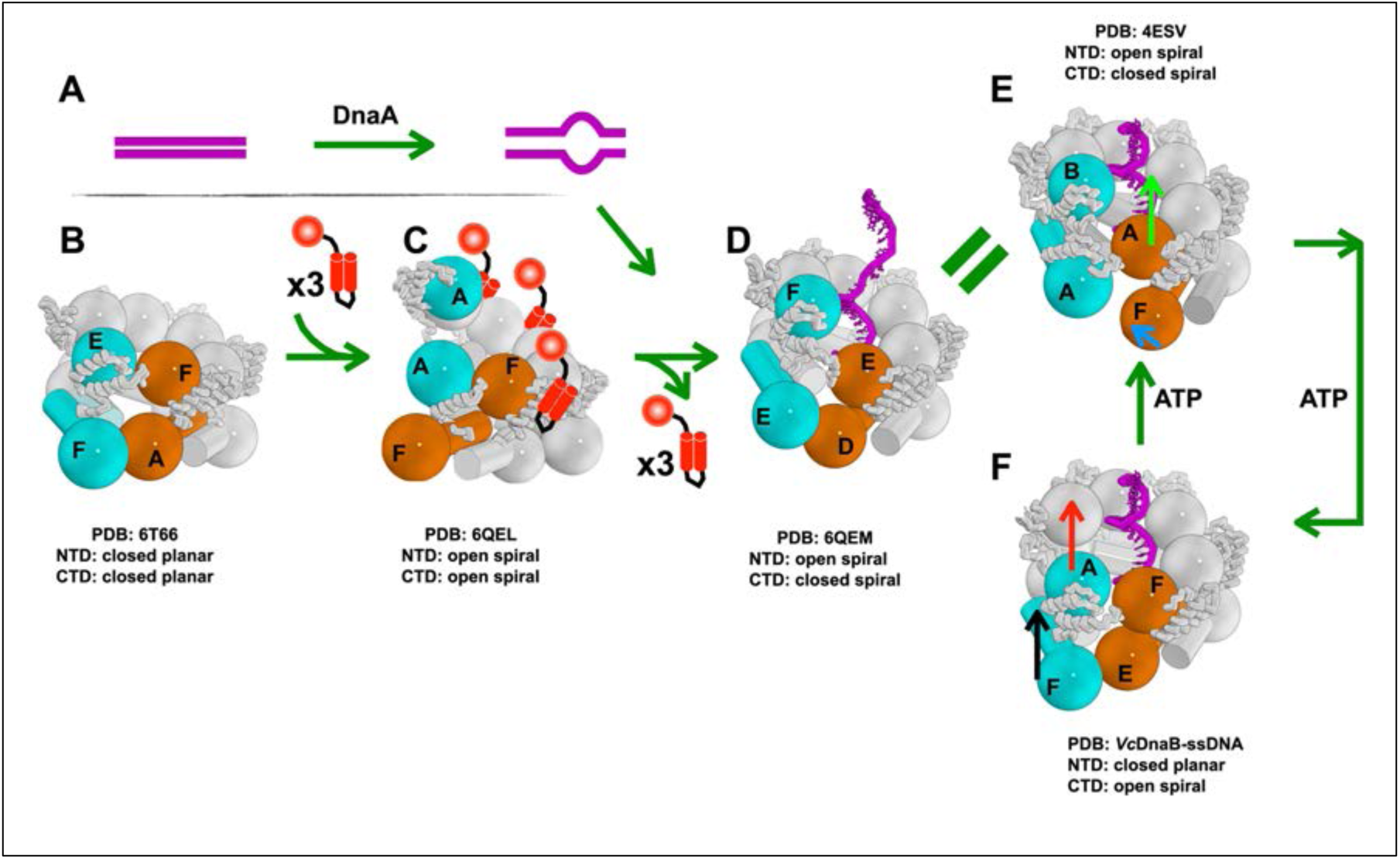
Model for Loading of DnaB by DciA and Translocation of DnaB on ssDNA. A) The DnaA protein binds to the replication origin and promotes melting into a DNA bubble. B) The hexameric closed planar DNA-free *Vc*DnaB (PDB: 6T66 (12)) provides a ground state model for the loading pathway. C) Our data suggest that up to 3 DciA loaders (red) bind to *Vc*DnaB; we suggest that DnaB adopts an open spiral configuration analogous to that seen with the DnaC/λP complexes (19, 43, 44, 48). D) Remodeling of this complex accompanies the entry of DnaA-generated ssDNA into the central chamber of DnaB. The current structures suggest that the D and E conformations are identical (19). We suggest that DnaB populates the two states captured in the GDP•AlF_4_ (E) and ATPγS (F) conformations during translocation. The first translocation step (panel F to panel E) involves opening the NTD tier and the migration of a CTD in the ATPγS complex to its position in the GDP•AlF_4_ complex. The ATPγS-chain F NTD shifts (black arrow) in position to the location of the GDP•AlF_4_-chain A. Opening of the NTD layer is accompanied by upward migration by 25 Å of the cyan CTD chain A in the ATPγS complex (panel F, red arrow) to the position occupied by chain B in the GDP•AlF_4_ complex (panel E); this represents the first translocation step. In the second step (panel E to panel F), the GDP•AlF_4_-NTD layer (chain F) returns (blue arrow) to the ATPγS-closed planar configuration (chain E). This change is accompanied by a 25 Å upward migration of the orange CTD chain A in panel E to a new position (chain F) in panel F (green arrow). Repetition of these two steps around the DnaB enables each subunit to translocate. We suggest that the DciA-loaded DnaB-ssDNA complex, which resembles the GDP•AlF_4_ conformation, enters this cycle and then transitions to the ATPγS state. The DnaB models in panels B-F are colored in white except for certain blue and orange CTDs and NTDs that line the various breaches, except for the closed planar entity.

Our structural analysis did not reveal the architecture of the *Vc*DnaB-*Vc*DciA-ssDNA complex. Nevertheless, the structure of the ATPγS form of the DnaB-ssDNA complex provides new insights into the mechanism of translocation. The topologically closed NTD and distinctly configured CTD layers distinguish the ATPγS state from a structure previously determined bound to the transition state ATP analog GDP•AlF_4_ (10) and provide a first view of nucleotide-dependent conformational change during ssDNA translocation. Our model for ATP-dependent translocation by DnaB extends the hand-over-hand mechanism (10) by integrating cycles of opening/closing of the NTD layer, reconfiguration of the LH element, and disengagement of individual CTDs from ssDNA and neighboring subunits, followed by migration from the bottom to the top of the DnaB spiral to new ssDNA binding sites. Migration to the top of the spiral is facilitated by the engagement of its β-hairpin arginine finger with a freshly bound ATP by the DnaB subunit that precedes arrival at the top of the spiral. ATP binding also stabilizes contacts to ssDNA. We suggest that translocation encompasses three identical sub-steps. Each sub-step involves migration of the chain A CTD (or chain E or chain C) with the simultaneous opening of the NTD layer into an open spiral, as seen in the GDP•AlF4 4ESV structure. Notably, the open spiral state of the NTD tier would likely influence contacts with the DnaG primase (9, 97), whose complete binding site on DnaB requires a closed NTD tier. In the second step, migration of the next CTD (chain F, chain D, or chain B) in sequence, newly at the bottom of the spiral, is accompanied by the closure of the NTD layer to the triangular planar arrangement seen in the *Vc* ATPγS structure. Our model specifies that two DnaB subunits move in sequence and that this program of motion is repeated thrice around the hexamer.

The precise mechanisms associated with ATP dynamics in ring-shaped oligomeric ATPase are of intense interest (10, 22, 28, 29, 98–102). Substantial evidence has established sequential ATP hydrolysis during rotary catalysis by the ATP synthase (103, 104). Within the AAA+ family of ATPases, the picture is far less clear, with studies pointing to sequential, stochastic mechanisms, both simultaneously (reviewed in reference (105)) or a concerted mechanism (106). Our proposal that two adjacent DnaB subunits undergo nucleotide dynamics and motion in sequence is compatible with sequential or stochastic mechanisms, as it disfavors a concerted scheme. Additional studies will be required for a more precise answer.

To link the significant findings of our work in DciA-mediated loading and subsequent translation by DnaB, we looked to another ring-opener, DnaC, to provide models for other steps in the helicase loading pathway (Figure 9). The configuration of DnaB in the *E. coli* DnaB-DnaC•ssDNA (PDB: 6QEM) complex (19) closely resembles that of the 4ESV GDP•AlF_4_ state (10), with a closed spiral CTD and open spiral NTD and contacts to ssDNA by each subunit. DnaB in the 6QEM and 4ESV structures is nearly identical. Thus, on the departure of the loader, DnaB enters the two-subunit motion (Figure 9E-F) scheme proposed above. Notably, the departure of the DciA loader produces a DnaB configuration after it has taken the first of the two sequential steps described above (Figure 9D). We suggest that after the eviction of the loader, the subunit at the bottom of the spiral will take the second of these steps to migrate to the top of the spiral (Figure 9E-F). Repetition of the sequenced motions of two DnaB subunits around the ring accompanies further translocation along ssDNA.

A crystal structure of the 6:6 *Vc*DnaB-*Vc*DciA complex implies two distinct DciA binding modes and a nearly closed planar DnaB ((14) and Supplementary Information and Supplementary Figures 26 and 27). Additional studies will be required to sequence the binding modes in this structure and others that might be present in the helicase loading reaction.

An important unanswered question centers on the mechanism of expulsion of DciA from the DnaB-ssDNA complex. The DnaC (19, 60, 107) and λP (108, 109) systems feature biochemical programs to mediate loader dissociation. Our data reveal a tight interaction between helicase and loader (Figure 6) that suppresses DnaB’s ATPase activity (Figure 7). Yet, DciA exhibits a dose-dependent stimulation of DnaB’s unwinding activity (Figure 8 and Supplementary Figure 19), absent any other factor. These findings suggest that DciA may spontaneously dissociate to relieve inhibition of DnaB’s ATPase activity and enable unwinding activities. It is also possible that interactions with other replisome components (25, 107, 110) could contribute to the eviction of DciA. Future studies will have to clarify this question.

## Data Availability

Coordinates for the EM-derived model of the *Vc*DnaB-ssDNA-ATPγS complex are available from the RCSB Protein Data Bank under accession codes 9DLS. Cryo-EM maps are available from the EMBD under the accession code EMD-46984.

## Author Contributions

NGao, DM, RM, and DJ conceptualized the study. NGao designed and prepared all the proteins. NGao and DM performed the ATPase assays. NGao performed the pull-down assays. NGouda, ST, AM, AChowdhury, ACerullo, and HB assisted with various laboratory experiments. DM performed the fluorescence-based DNA unwinding and helicase loading assays. DM and CM performed the surface plasmon resonance analyses. All nMS analyses were performed by PDBO in the BTC lab. AP and NGao acquired the cryo-EM data and performed the initial data analysis. DM analyzed the cryo-EM data. DM built and analyzed the cryo-EM structure. NGao, DM, RM, and DJ drafted the manuscript, working with FR and MR.

## Acknowledgments

We thank the Jeruzalmi and Miggiano lab members, the City College biophysics group, the CUNY Advanced Science Research Center, and the New York Structural Biology Center for their scientific and technical advice. We also thank Professor Anuradha Janakiraman for excellent scientific discussions and technical assistance.

## Supplementary Information

### Assembly of Hexameric Helicases on Nucleic Acid Substrates

Cells apply three broad strategies to load hexameric helicases onto nucleic acid substrates: ring-opening, ring-forming, and ring-closing (1–3); the terms refer to the initial state of the helicase. These schemes encompass loading onto ssDNA, ssRNA, and dsDNA. In the ring-opening scheme, closed-ring hexamers, e.g., *E. coli* DnaB, are opened by helicase loaders, such as *E. coli* DnaC (4–6) or phage λP (7–9), and assembled on ssDNA formed at the cognate replication initiator– origin DNA complex. In the ring-forming scheme, the hexameric helicase is assembled on ssDNA from monomers, either spontaneously as with the LT-antigen or E1 helicases or through the activity of a helicase loader such as DnaI, which loads the DnaB family helicase in *B. subtilis* (10, 11). The ring-closing scheme is observed with the bacterial Rho and the eukaryotic MCM2-7 hexameric replicative helicases (reviewed in (1–3)). In the Rho helicase, ring closure is mediated by the entry of the nucleic acid substrate. In the MCM2-7 ensemble, dsDNA is deposited via the chaperone origin recognition complex and associated factors. The scheme(s) the DciA loader uses to assemble DnaB onto the origin-DNA are unknown.

### *Vc*DnaB Primarily Contacts the Phosphate Backbone of ssDNA

*Vc*DnaB subunits use seven residues (Asn353, Thr355, Asn383, Arg384, Arg400, Glu401, and Gly403) to contact the ssDNA backbone, though not every chain uses every residue; one contact is seen to contact a nucleobase (chain D Asn353 and N2 of G5). The segment of ssDNA within DnaB’s central chamber is arranged with 11-12 nucleotides per turn of ssDNA and a pitch of ∼32 Å, a value that compares well with the pitch (∼33.4 Å) displayed by the DnaB CTD spiral. The approximate match in pitch between protein and ssDNA spirals implies that corresponding DNA binding residues on each subunit make equivalent contacts with the ssDNA backbone (12) (Supplementary Figure 11 and Supplementary Table 4).

### *Vc*DciA Stimulated Unwinding by *Vc*DnaB Measured Using a Replication Fork-Mimic

Every bacterial helicase loader stimulates the DNA unwinding activity of its cognate DnaB helicase. To investigate the influence of DciA on cognate DnaB, we applied an ensemble FRET-based DNA unwinding assay wherein a DNA replication fork mimicking substrate (5) is fitted with a Cy3 labeled nucleobase Watson-Crick-Franklin paired with a nucleobase conjugated to the BHQ2 quencher; the unwinding of the fork substrate is readout by recording appearance of a fluorescent light signal at 570 nm (Supplementary Figure 19). This assay showed that 200 nM *Vc* DnaB exhibited measurable albeit low DNA unwinding activity. However, titration of full-length *Vc* DciA (0.11 to 6400 nM) revealed a 10-fold (15 to 150) increase in fluorescence at the highest concentration (6400 nM) (Supplementary Figure 19A). Unwinding depended on ATP and was significantly reduced in the *Vc*DnaB Walker B mutant E259A (Supplementary Figure 20). Parallel analysis of the isolated DciA KH and Lasso domains implied retention of about 30% of the activity of the full-length *Vc*DciA to activate the helicase. (Supplementary Figures 19B and 21).

### *Vc*DciA Stimulated Unwinding by *Vc*DnaB Measured Using a Doubly Labeled DNA Bubble Substrate

The bubble DNA substrate (Figure 8 and Supplementary Figure 23), in principle, permits the loading of DnaB onto both single strands, leading to unwinding in both directions; however, the bubble substrate above with the single FRET pair only reports on activity at one end of the DNA. To examine activity at both ends, we repeated the helicase loading assay with a bubble substrate that featured the above FRET pairs at both termini. Full-length DciA and the two truncated constructs displayed the same pattern of activity as seen with the singly labeled substrate (Supplementary Figures 23, 25C, and 25D). Full-length DciA produced 10% more fluorescence on the doubly labeled substrate instead of the expected doubling of the signal. This discrepancy may be due to unexpected quenching between the two ends of the molecule due to conformational switching that brings them closer together. Alternatively, loading two DnaB molecules may be more complex than previously thought.

### Comparison to other structures of *Vc*DnaB

We compared our cryo-EM *Vc*DnaB structure to three previously determined crystal structures: 6T66, 7QXM, and 8A3V. None feature ssDNA (6T66, 7QXM), and one includes DciA at a 6:6 ratio with DnaB (8A3V). We compared the structures for subunit proximity, ring breaches, nucleotide occupancy, the β-hairpin/arginine finger disposition, and LH position.

The 6T66 structure (13) presents as a planar hexamer in the crystallographic asymmetric unit with a dilated NTD layer and a constricted CTD tier (Supplementary Figure 26A). The DnaB subunits in this structure are labeled A-B-C-D-E-F, counterclockwise down the CTD axis. Two of the CTDs (A – F: 35.1 Å and D – E: 36.4 Å) are 2-3 Å further away from their neighbors than the disposition of other subunit pairs (A–B: 32.8 Å; B–C; 32.7 Å; C–D: 32.7 Å; E– F: 33.1 Å), implying two breaches in the CTD layer; the NTD layer is not breached. In the *Vc*DnaB-ssDNA complex, chain A, at the bottom of the spiral, is further away (A–B: 39.6 Å; F–A: 34.6 Å) from its neighbors than other pairs (average: ∼33.4 Å). The significance of two breaches in non-consecutive subunits in the CTD tier in the 6T66 structure remains to be understood.

The nucleotide-binding site of each subunit in 6T66 is populated with GDP•AlF_4_/Mg^2+^. However, the β-hairpin/arginine finger elements that complete these binding sites populate three positions. The distance between the P-loop lysine 234 and arginine finger 439 at the sites between chains D to E (18.7 Å) and F to A (16.0 Å) is considerably longer than the corresponding values for other subunit pairs (A–B: 10.9 Å; B–C: 11.2 Å; C–D: 11.1 Å; E–F: 11.3 Å). Naturally, longer distances are seen in CTDs that are further apart (Supplementary Figure 26C).

We next examined the positions adopted by the LH element in the 6T66 hexamer. Alignment of the CTD onto the chain A CTD, but omitting the DH and LH elements from the calculation (resid 204:465 and not (resid 287:304 or resid 178:205)) produces an average RMSD = 0.74 Å. In this alignment, the LH elements populate two positions (LH1 and LH2 above). The LH elements from chains A and E populate positions closely related to LH2; the rest are seen in the LH1 position (Supplementary Figure 26D). The subunit pairs with divergent dispositions and arginine fingers populate the LH2 position; the rest are in LH1.

The 7QXM *Vc*DnaB crystal structure features an unliganded planar hexamer in the asymmetric unit (14). The DnaB subunits in this structure are labeled A-F-E-D-C-B, counterclockwise down the CTD axis. The NTD layer is dilated and follows NTD layers from 6T66 and the *Vc*DnaB-ssDNA complex. The CTD layer is constricted, with individual CTD further away from each other than in 6T66 or the *Vc*DnaB-ssDNA complex (A–B: 33.7 Å; B–C: 37.0Å; C–D: 33.7 Å, D–E: 36.3; E–F: 33.6 Å; F–A: 37.9 Å). Likewise, the P-loop lysine to arginine finger distances are variable (A–B: 21.1 Å; B-C: 16.8 Å; C–D: 20.4 Å, D–E: 15.7 Å; E–F: 20.9 Å; F–A: 17.3 Å). The observed variability in the disposition of CTDs in 7QXM may reflect the underlying lack of nucleotide to organize and stabilize inter-CTD contacts via the β-hairpin arginine finger. The LH elements in 7QXM populate two positions, LH2 (Chains B, D, and F) and another position (Chains A, C, and E), which are unique to this structure.

The 8A3V *Vc*DnaB-DciA crystal structure has two DnaB subunits and two DciA molecules in the crystallographic asymmetric unit (15). Applying crystallographic symmetry generates a 6:6 DnaB: DciA entity, revealing a dilated NTD layer and a constricted CTD layer. The nucleotide site of each *Vc*DnaB CTD is filled with ADP. In the *Vc*DnaB_2_: *Vc*DciA_2_ asymmetric unit, the inter-CTD distance is ∼32 Å, and the P-loop to arginine finger distance is 12.3 Å. The LH elements in 8A3V occupy the LH1 position (Supplementary Figure 27).

Although parallels could be drawn between the 6T66, 7QXM, and 8A3V structures and the present *Vc*DnaB-ssDNA structure, additional studies will be required to sequence them precisely in the reaction coordinate.

### Structural Comparisons to *E. coli* DnaB-ssDNA Complexes

The RCSB PDB includes four unpublished entries corresponding to *E. coli* DnaB-ssDNA structures (PDB: 7T20, 7T21, 7T22, and 7T23 (16–19)). One structure was determined with the ATP analog AMP-PNP (7T20); the others harbor the ADP•AlF_4_ transition state analog (7T21, 7T22, and 7T23). The ATP analog structure maps closely to the *Vc* structure, while the ADP•AlF_4_ transition state DnaB structures map to the Bst complex. As such, conclusions made from comparisons between the *Vc* and Bst structures are fully compatible with the *E. coli* structures.

## Supplementary Methods

### Expression Plasmid Construction

Genes corresponding to full-length *Vibrio cholerae* (*Vc*) DnaB and *Vc*DciA were amplified using polymerase chain reaction (PCR) and genomic DNA from *Vibrio cholerae* N16961 (ATCC # 39315D-5), and variously, cloned into the following protein expression vectors (pET24a, pCDFDuet-1, or pRSFDuet-1). These efforts encompassed untagged, amino, or carboxy-terminal hexahistidine-tagged proteins. Point mutations were introduced using the Quickchange Site Directed Mutagenesis kit (Agilent).

*E. coli* DnaB was prepared as previously described of the *E. coli* DnaB•λP complex, except that λP was omitted (9).

### Protein Expression

For production-scale bacterial growths, colonies corresponding to bacterial transformants were picked and resuspended in LB media supplemented with the appropriate antibiotic concentration. Bacterial cultures were grown at 37 °C to an OD600 ≈ 0.6 – 0.7 (RPM = 180). Protein expression was induced by making the culture 0.5 mM in isopropyl-D-1-thiogalactopyranoside (IPTG), followed by a temperature shift of the culture to 16 °C; protein production was allowed to continue for 14-16 hours (RPM = 180). Cells were recovered by centrifugation, resuspended in 20mM Tris-HCl pH 7.4, 500mM NaCl, 5mM MgCl_2_, 10% sucrose, 5mM β-mercaptoethanol at a ratio of 5 mL buffer per gram of wet cells, and frozen at −80°C.

#### *Vc*DnaB

*Vc*DnaB-untagged (pNG066, 50 µg/mL kanamycin), *Vc*DnaB-NHis (pNG067, 50 µg/mL kanamycin), *Vc*DnaB-E259A-NHis (pNG072, 50 µg/mL kanamycin), *Vc*DnaB-NTD-untagged (pNG114, 50 µg/ml kanamycin), *Vc*DnaB-NTD_LH-untagged (pNG131, 50 µg/mL kanamycin) and *Vc*DnaB-LH_CTD-untagged (pNG134, 50 µg/mL kanamycin) were expressed using the above procedure.

#### *Vc*DciA

*Vc*DciA-FL-CHis (pNG057, 50 µg/ml streptomycin) and *Vc*DciA-CTD-CHis (pNG087, 50 µg/ml streptomycin) were expressed using the procedure above, except for three changes: a) the media was LB for pNG057 and SB (20, 21): for pNG087, b). after adding 0.5 mM IPTG to the culture, protein expression was allowed to continue for 3 hours at 37 °C, and c) cells were resuspended after harvesting in 20 mM Tris-HCl pH 8, 300 mM NaCl, 5% glycerol, 5 mM β-mercapoethanol.

*Vc*DciA-NTD-Chis (pNG078, 50 µg/ml streptomycin) was grown using the same procedure as full-length DciA, except that cells were resuspended in 20 mM Tris-HCl pH 8, 300 mM NaCl, 5% glycerol, and 5 mM β-mercapoethanol.

#### *Vc*DnaB-*Vc*DciA coexpression

*Vc*DnaB-untagged (pNG066, streptomycin: 50 ug/mL) and *Vc*DciA-FL-Chis (pNG057, Kanamycin: 50 ug/mL) were co-expressed using the above procedure altered to account for the co-expression.

### Protein Biochemistry

Unless otherwise indicated, standard protein biochemistry techniques were used to purify all the proteins in this study (Supplementary Figure 8). All manipulations were carried out at 4 °C.

Unless otherwise indicated, 90 - 160 mL of cells expressing the protein of interest were lysed using a Sonic dismembrator (Model 500, Fisher Scientific) at the following settings: 3 min at 70% amplitude, with alternating 30s on and off cycles and 15 min at 50% amplitude. The protein of interest was recovered in the soluble fraction by centrifugation (RPM=16,000, JA-17, Beckman-Coulter).

#### *Vc*DnaB

Untagged *Vc*DnaB was purified by combining ammonium sulfate (AS) precipitation and heparin, hydroxyapatite, and size exclusion chromatography. *Vc*DnaB was found in the supernatant after making the soluble fraction 20% of saturated AS (SAS), but in the pellet when treated with 45% SAS. The 45% SAS pellet was resuspended in heparin buffer A below and purified over a 50 mL Heparin Sepharose Fast Flow (Cytiva), with the following buffers: heparin buffer A: 20 mM Tris-HCl, pH = 7.4, 5 mM MgCl_2_, 5% glycerol, 5 mM β-mercaptoethanol; Heparin buffer B: A + 1 M NaCl and a linear gradient from 0% to 40% B. Heparin-purified *Vc*DnaB was purified by hydroxyapatite chromatography (65 ml, CHT, Bio-Rad) in the following buffers: A: 5mM potassium phosphate, pH = 7.5, 100 mM NaCl, 5 mM MgCl_2_, 5% glycerol, 5 mM β-mercaptoethanol; B: A + 250 mM potassium phosphate, pH = 7.5). CHT-purified *Vc*DnaB was concentrated by 45% AS precipitation, redissolved in 6 mL of 20 mM potassium HEPES, pH=7.5, 500 mM NaCl, 5 mM MgCl_2_, 5% glycerol, 5 mM β-mercaptoethanol), and applied to a 320 mL Superdex200 Prep Grade (Cytiva) column equilibrated in the above buffer. Size exclusion chromatography fractions free of nuclease activity were pooled, concentrated (Corning, 20 ml, 10 kDa molecular weight cutoff) to 55 mg/ml, aliquoted (∼10-20 µl aliquots), flash frozen in liquid nitrogen, and maintained at - 80 °C until use.

*Vc*DnaB-NHis and *Vc*DnaB-E259A-NHis were purified using the protocol employed with untagged *Vc*DnaB but with discrete changes. The 45% SAS pellet was resuspended in Ni^2+^-nitrilotriacetic acid (NiNTA) Buffer A (20 mM Tris-HCl, pH = 7.4, 50 mM NaCl, 5 mM MgCl_2_, 5% glycerol, 5 mM β-mercaptoethanol) and incubated with 15 mL of NiNTA agarose beads (Qiagen) according to the manufacturer’s instructions. The beads were washed with 180 mL of NiNTA Buffer B (20 mM Tris-HCl, pH=7.4, 50 mM NaCl, 5 mM MgCl_2_, 5% glycerol, 5 mM β-mercaptoethanol, 30 mM imidazole pH 8); *Vc*DnaB-NHis and was eluted from the beads with 300 mL of NiNTA Buffer C (20 mM Tris-HCl pH=7.4, 50 mM NaCl, 5 mM MgCl_2_, 5% glycerol, 5 mM β-mercaptoethanol, 250 mM imidazole, pH = 8). NiNTA-purified *Vc*DnaB-NHis was further purified over Heparin Sepharose Fast Flow as was done with untagged DnaB, except that the elution gradient was modified; 4 column volumes of Heparin buffer A (above), followed by a linear gradient from 5% to 100% Heparin buffer B over six column volumes. *Vc*DnaB-NHis fractions were recovered and pooled in the ∼80 mM to ∼600 mM NaCl positions on the gradient. The heparin pool was passed through a 23 mL SP Sepharose Fast Flow (Cytiva) equilibrated in Q buffer A: 20 mM Tris-HCl, pH=7.4, 100 mM NaCl, 5 mM MgCl_2_, 5% glycerol, and 5 mM β-mercaptoethanol; *Vc*DnaB-NHis was captured by a 50mL Q Sepharose Fast Flow column (Cytiva) attached in-line. After a 4-column volume wash with 10% Q buffer B: (20 mM Tris-HCl, pH = 7.4, 1 M NaCl, 5 mM MgCl_2_, 5% glycerol, and 5 mM β-mercaptoethanol), a linear gradient from 10% to 100% Q buffer B was used to elute *Vc*DnaB-NHis, which was recovered in the ∼250 mM to ∼800 mM NaCl positions on the gradient. The pooled *Vc*DnaB-NHis containing fractions were further purified over hydroxyapatite (Bio-Rad, CHT) and Superdex200 Prep grade (Cytiva) as described above. Size exclusion chromatography fractions were pooled, concentrated as described above, to 40 mg/ml, aliquoted (∼15-30 µL aliquots), flash frozen in liquid nitrogen, and maintained at −80 °C until use.

#### *Vc*DciA

*Vc*DciA-FL-CHis, *Vc*DciA-NTD-CHis, and *Vc*DciA-CTD-CHis were purified using nearly identical protocols. Cells were lysed as above, except that Protease Inhibitor Mini Tablet, EDTA-free (Pierce, ThermoFisher Scientific p/n 32955), was added to the *Vc*DciA-CTD-CHis sample at a ratio of one tablet for every 10 mL of lysate. All three *Vc*DciA constructs were recovered in the soluble fraction. Each fraction was applied to ∼10 mL of Ni-NTA agarose beads, incubated for 40 min, and washed with 80 mL of NiNTA Buffer A (20 mM Tris-HCl, pH = 8, 300 mM NaCl, 5% glycerol, and 5 mM β-mercaptoethanol), supplemented with 15 mM imidazole, pH = 8 (*Vc*DciA-NTD-CHis), or 30 mM imidazole, pH = 8 (*Vc*DciA-FL-CHis and *Vc*DciA-CTD-CHis). *Vc*DciA constructs were eluted from the beads with NiNTA Buffer B (20 mM Tris-HCl, pH = 7.4, 50 mM NaCl, 5 mM MgCl_2_, 5% glycerol, 5 mM β-mercaptoethanol, and 250 mM imidazole, pH = 8). Further purification was achieved by flowing the various *Vc*DciA constructs through a 50mL Q Sepharose Fast Flow column for capture on a 23mL SP Sepharose Fast Flow column attached in-line; both columns had been equilibrated in SP buffer A: (20mM Tris-HCl, pH = 7.4, 50 mM NaCl, 5% glycerol, and 5 mM β-mercapoethanol). After separating the two columns, weakly bound proteins were washed away with 4 columns volumes of 5% SP buffer B (20 mM Tris-HCl, pH = 7.4, 1 M NaCl, 5% glycerol, and 5 mM β-mercaptoethanol); this was followed by a linear gradient from 5% to 100% SP buffer B over 3 column volumes. Notably, we observed that *Vc*DciA-FL-CHis eluted at the ∼150 mM to ∼350 mM NaCl position in the gradient, *Vc*DciA-NTD-CHis between ∼120 mM and ∼700 mM NaCl, and *Vc*DciA-CTD-CHis between ∼180 mM and ∼300 mM NaCl. Pooled fractions were concentrated (*Vc*DciA-FL-CHis by 50% AS precipitation, *Vc*DciA-NTD-CHis by 70% AS precipitation, and *Vc*DciA-CTD-CHis using a 15 mL (2 kDa molecular weight cutoff) spin concentrator (Sartorius Vivaspin). Each construct was then chromatographed over a 320 mL Superdex200 Prep Grade column, equilibrated in 20 mM potassium-HEPES, pH = 7.5, 300 mM NaCl, 5% glycerol, and 5 mM β-mercaptoethanol, and processed for −80° storage as described above.

#### Coexpressed *Vc*DnaB-*Vc*DciA

A soluble fraction derived from coexpressed *Vc*DnaB-untagged - *Vc*DciA-FL-Chis, prepared as described above, was submitted to NiNTA and size exclusion chromatography (320 mL Superdex200 Prep Grade). The NiNTA chromatography was performed as above with the following buffers: NiNTA buffer A (20 mM Tris-HCl, pH = 7.4, 500 mM NaCl, 5 mM MgCl_2_, 5% glycerol, 5 mM β-mercaptoethanol, and 30 mM imidazole, pH = 8) and NiNTA buffer B (20 mM Tris-HCl, pH = 7.4, 500 mM NaCl, 5 mM MgCl_2_, 5% glycerol, 5 mM β-mercaptoethanol, and 250 mM imidazole, pH = 8). NiNTA-purified protein was concentrated by precipitation with 75% SAS and redissolved in 6 mL of the running buffer (below) for size exclusion chromatography. The size exclusion chromatography was carried out in (20 mM potassium HEPES, pH = 7.5, 500 mM NaCl, 5 mM MgCl_2_, 5% glycerol, and 5 mM β-mercaptoethanol).

### NiNTA pull-downs

Equal volumes (22.5 mL) of cells at similar densities expressing untagged *Vc*DnaB (full-length, NTD, NTD-LH, or LH-CTD, each in 20 mM Tris-HCl pH 7.4, 500 mM NaCl, 5 mM MgCl_2_, 10% sucrose, 5 mM β-mercaptoethanol) and his-tagged *Vc*DciA (full-length, Lasso, or KH, each in 20 mM Tris-HCl pH 8, 300 mM NaCl, 5% glycerol, 5 mM β-mercapoethanol), were mixed, diluted with 15 mL of buffer (20mM Tris-HCl pH 8, 300mM NaCl, 5% glycerol 5 mM β-mercapoethanol) were combined to a total volume of 60 mL. The mixture was lysed using a Sonic dismembrator (Model 500, Fisher Scientific) at the following settings: 2 min at 70% amplitude, with alternating 30s on and off cycles. Sonication settings were: 30s on, 30s off, 2 min total on time, 70% amplitude. The lysate was centrifuged at 16k RPM in a JA-17 rotor (Beckman-Coulter), and the supernatant was recovered.

The clarified lysate was incubated with Ni-NTA Agarose beads for 40 minutes, washed with 90 mL of NiNTA Buffer A (20 mM Tris-HCl pH 8, 300 mM NaCl, 5% glycerol, 5 mM β-mercaptoethanol, 15 mM Imidazole pH 8). Protein was eluted using 30 mL of NiNTA Buffer B (20 mM Tris-HCl pH 8, 300 mM NaCl, 5% glycerol, 5 mM β-mercaptoethanol, 250 mM Imidazole pH 8).

### ATPase Rate Measurements

An NADH-coupled microplate spectrophotometric assay was used to measure rates of ATP hydrolysis (22, 23) by *Vc*DnaB and how rates are affected by the presence of our suite of full-length and truncated constructs of *Vc*DciA. In all experiments, *Vc*DnaB was kept fixed at 200 nM, while titrations of DciA encompassed concentrations between 0 nM and 6400 nM.

NADH oxidation data were recorded in the presence of 2 mM ATP (Gold Biotechnology, Inc. p/n: A-081-1), 2 mM phosphoenolpyruvic acid monopotassium salt (PEP), 0.01 U/µL lactate dehydrogenase (Sigma-Aldrich, L1254), 0.002 U/μL pyruvate kinase (Sigma-Aldrich, p/n: P7768,) and 0.3 mM NADH in the following buffer: 50 mM potassium-HEPES – pH=7.5, 150 mM potassium acetate, and 8 mM magnesium acetate, 5 mM β-mercaptoethanol, and 0.25 mg/mL bovine serum albumin (Sigma-Aldrich p/n: A3912). The above components were combined, save for ATP, into wells of a Corning 96-well Clear Flat Bottom UV-Transparent Microplate (p/n: 3635), and the plate was warmed to 37 °C for 5 minutes. After the quick addition of 2 mM ATP, the plate was inserted into a SpectraMax M5 plate reader (Molecular Devices), and absorbance data were taken at 340 nm every 30 s over 30 min. Raw data were exported in text format and imported into Microsoft Excel for analysis. Each absorbance curve was manually inspected to identify 10 minutes during the experiment wherein the slope of the 340 nm absorbance vs time was constant. The values in the above period were divided by the (-) 6.22 mM^-1^ cm^-1^ extinction coefficient of NADH and the path length in cm calculated by the plate reader to result in the rate of ATP hydrolysis in mM/s; further division by the mg of DnaB in the assay provided the specific activity in mM/s/mg.

### Grid Preparation for Single Particle Cryo-EM

To prepare complexes for cryo-EM, untagged *Vc*DnaB-FL and *Vc*DciA-FL-CHis were diluted in 20 mM HEPES pH = 7.5, 200 mM NaCl, 5 mM MgCl2, 5 mM β-mercaptoethanol, and 5 mM ATPγS to reach a final concentration of 18 μM (DnaB monomer) and 12.7 μM (DciA). A ssDNA oligonucleotide bearing the origin of the replication sequence of *Vibrio cholerae* serotype O1 (Supplementary Table 4, oligo: O1b-*Vc*F) was added at a concentration of 5.4 μM. 3 μL of the *Vc*DnaB-DciA-ssDNA sample was applied to a Quantifoil UltrAuFoil R1.2/1.3 Au300 grid, previously plasma cleaned with a 3:1 ratio of Ar/O_2_ at 15 W for 10 seconds in a Gatan Solarus II glow discharger. Liquid absorption was allowed for 30 seconds at 4 °C and 100% humidity. The sample was then blotted for 4 seconds with a blot force of 4 and plunge-frozen into liquid ethane. The procedure was conducted on a Vitrobot Mark IV (Thermo Fisher). The grid was stored in liquid nitrogen until data collection.

### Cryo-EM Data Acquisition

Cryo-EM images were acquired on a Titan Krios (ThermoFisher) operating at 300 keV and fitted with a Gatan K3 direct electron detector. 12,265 movies were collected using Leginon (24, 25) at an 81,000 magnification, corresponding to a pixel size of 0.541 Å/pixel, and at a defocus of −1.57 μm. The total exposure dose was 51.37 e–/Å^2^, and the exposure time per movie was 2 seconds. Additionally, 20° and 40° tilted images were collected to mitigate the expected preferred particle orientation that would otherwise compromise the EM reconstruction.

### Cryo-EM Image Processing

The 12,265 images measured from the *Vc*DnaB-DciA-ssDNA-ATPγS complex were processed using two approaches (Supplementary Figure 3): a) template-based picking using a prior DnaB structure (PDB entry = 7T20 (17)) as the initial template, and b) blob-picking. For facile computing, the blob-picking approach initially targeted 10% of the images, which produced 2.5 million particles that were 2D classified and submitted to *ab initio* reconstruction with C1 symmetry enforced. This reconstruction generated two volumes, C0 and C1 (∼123K and ∼151K particles), which were judged to be of low quality and influenced our decision not to pursue the blob-picking approach further.

For the PDB structure-based template-picking approach, for computational ease, we selected 100 frames at random, binned them by a factor of ½, and submitted them for motion correction and Contrast Transfer Function (CTF) estimation as implemented in cryoSPARC (26–32). Unless otherwise indicated, the following steps were performed within cryoSPARC. An EM map was calculated from PDB entry 7T20 in EMAN2 (33) and used to derive templates for application to the above 100 images; this procedure netted 92,178 particles. From this collection, 79,820 particles were extracted using a box size of 360 pixels and 2D classification. Subsequently, *ab initio* reconstruction was performed with C1 symmetry, which produced two classes (A0, A1). Class A0 (18.5K particles) was carried forward to homogeneous refinement, which created an 8.72 Å map (B0). Inspection of Class A1 suggested a lower quality than A0. As such, further processing of Class A1 was not pursued.

The complete data set (12,265 images) was then submitted to motion correction (Fourier crop factor of ½) and CTF estimation using standard settings. These calculations enabled manual curation leveraging ice thickness (0.82-1.1) and CTF fit resolution (2.17-9.98) parameters to eliminate images with low overall resolution. The resulting 11,817 images were submitted to template-based picking using the B0 map; this calculation produced ∼4.3 million particles. These particles were extracted as above and submitted to 2D classification and *ab initio* reconstruction using C1 symmetry. The resulting volumes, D0 and D1 (∼437K and ∼291K particles), were submitted to heterogeneous refinement and produced two volumes, E0 and E1 (∼526K and ∼202K particles). Inspection of these two volumes implied that the 6.24 Å E0 volume was of higher quality.

Particles from the E0 Class were re-extracted using a box size of 360 pixels, with no Fourier crop, and submitted to non-uniform refinement, which yielded an EM map with a resolution of 3.36 Å (Volume F). To further improve the quality of our reconstruction, volume F particles were submitted to global CTF refinement. The final non-uniform refinement job produced map G at 3.37 Å, which was further sharpened with DeepEMhancer (34) to produce volume H; this volume embodied our best map and was used for model building (below). EM density was evident for the protein, nucleic acid, and nucleotides. Indeed, estimates of the local resolution ranged from 2.8 Å to 4.8 Å, with the highest resolution density spanning the DnaB-CTDs (Supplementary Figure 4C). The quality of the protein density was sufficient to allow unambiguous assignment of side chains to the map. However, no density could be assigned to the DciA loader.

Next, we made several efforts to improve the resolution of Map H in hopes of locating the DciA loader. 3D classification of the particles that generated Map F produced ten classes (group I). Classes I1 and I9 retained the highest number of particles, 82K and 77K, respectively, and showed high similarity to Map H, but with no features that could be assigned to *Vc*DciA. Non-uniform refinement was performed on volumes I4 (46K particles) and I6 (59K particles), producing maps at the resolution of 3.76 Å (class J4) and 3.78 Å (class J6). Automatic model building with ModelAngelo (35) on the sharpened map, once again, failed to show any evidence of *Vc*DciA. Homogeneous refinement of the particles that produced the E0 volume created class L, followed by two consecutive rounds of non-uniform refinements (classes M and N) with different settings, using volume E0 as the input map. This procedure generated Map N, followed by Map O after DeepEMhancer sharpening, with both at resolutions of 3.45 Å. Neither map showed evidence of density better than Map H, let alone proof of the presence of *Vc*DciA. Lastly, inspection of the 3D variability analysis of the particles that produced volume F (group K) produced no evidence of conformational changes in the *Vc*DnaB-ssDNA complex.

### Supplementary Figures

**Supplementary Figure 1.**
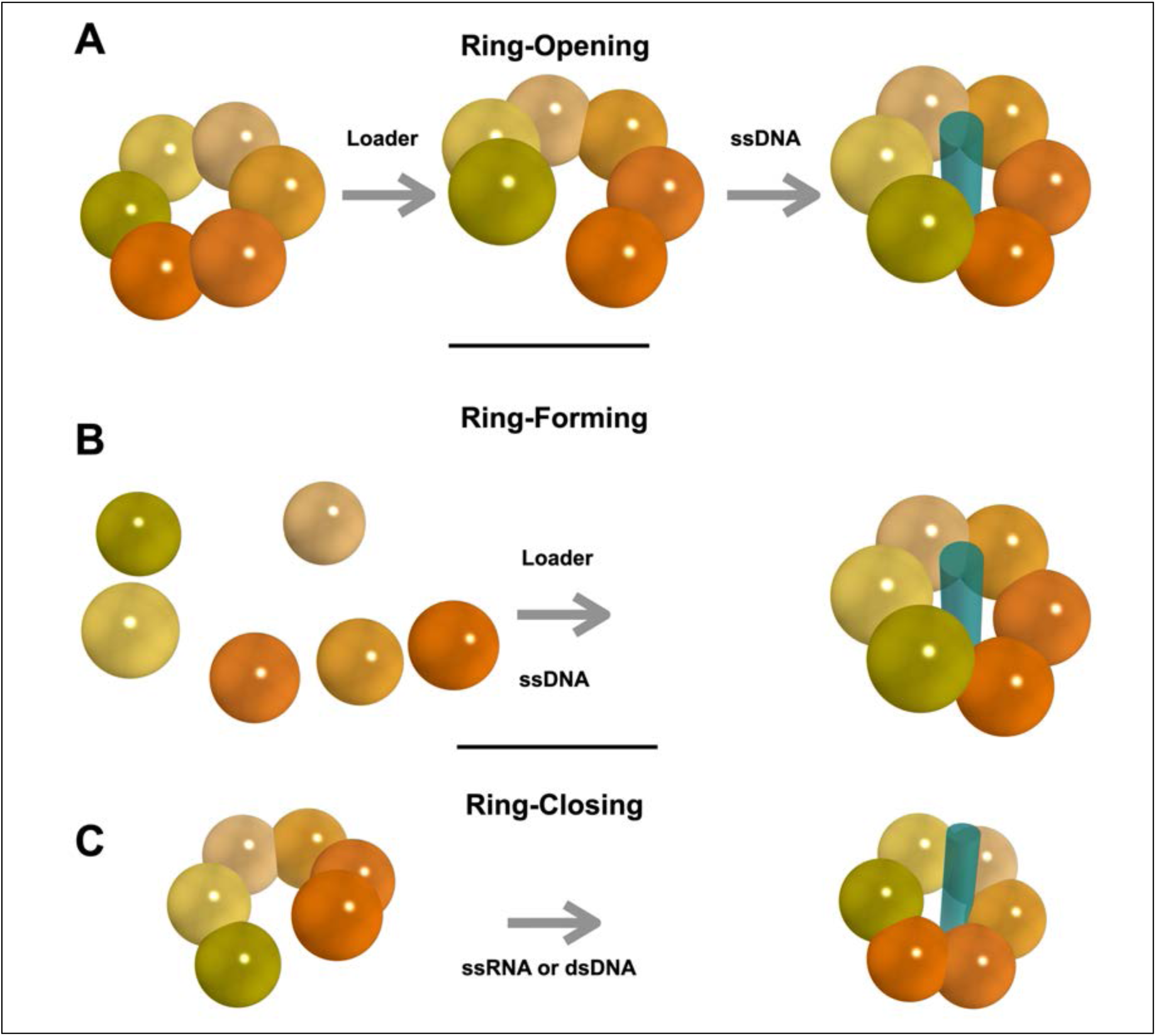
Mechanisms of Helicase Loading. Strategies used to load hexameric helicases onto nucleic acid substrates are ring-opening, ring-forming, and ring-closing (1–3); the terms refer to the initial state of the helicase. In the ring-opening scheme, closed-ring hexamers are opened by helicase loaders and subsequently assembled on ssDNA. The hexameric helicase is assembled on ssDNA from monomers using a ring-forming mechanism. Entry of the nucleic acid substrate and other factors mediate the ring-closing scheme. Hexameric helicase subunits are represented as spheres in shades of orange. For clarity, only the ATPase domains are shown. Nucleic acid substrates are depicted as deep teal cylinders.

**Supplementary Figure 2.**
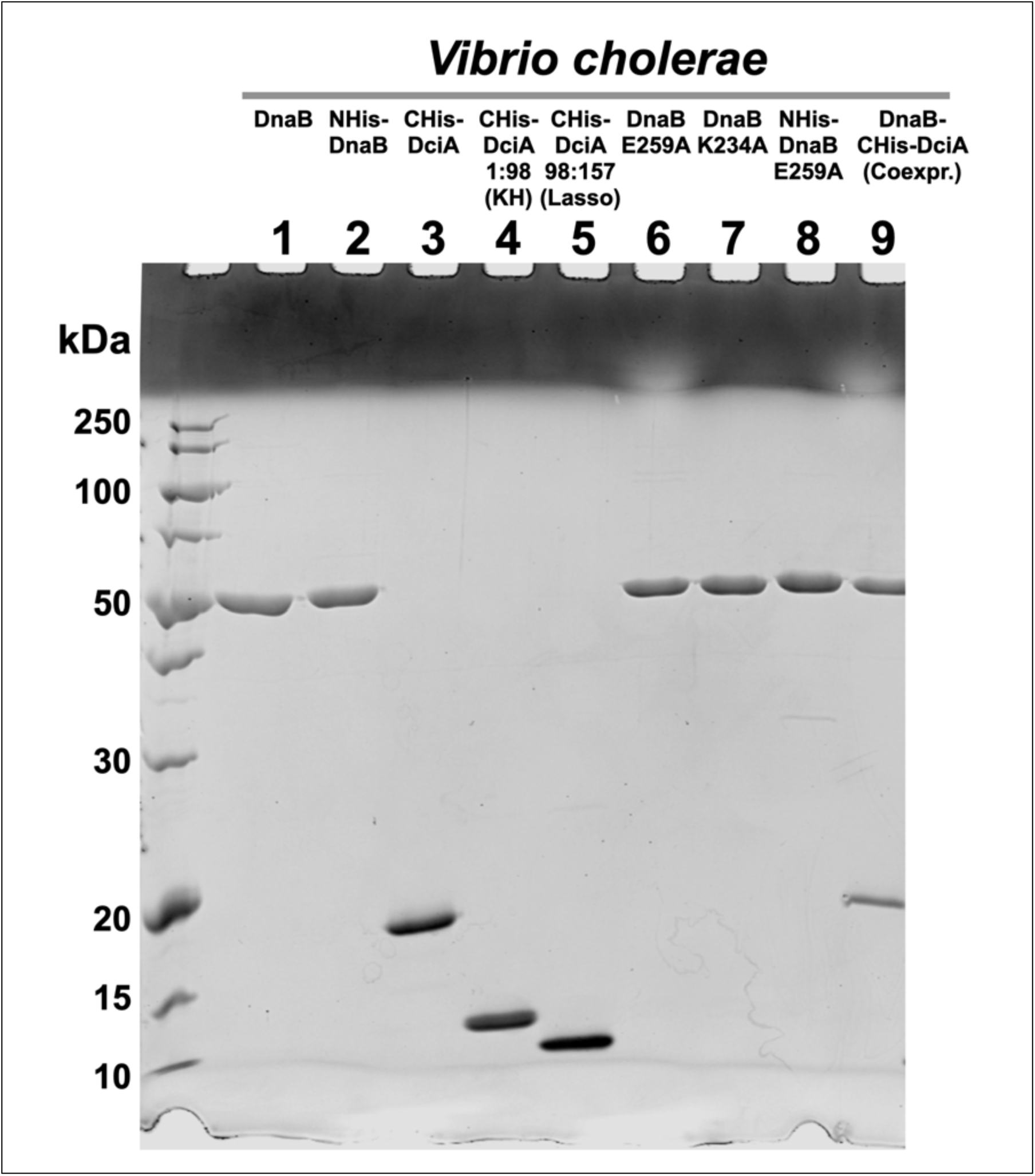
Purified *Vc*DnaB and *Vc*DciA proteins. A) 15% SDS-Gel electrophoretic analysis of purified *Vc* proteins. Lane 1: *Vc*DnaB untagged; Lane 2: N-terminal His-tagged *Vc*DnaB; Lane 3: C-terminal His-tagged *Vc*DciA full-length; Lane 4: C-terminal His-tagged *Vc*DciA KH domain (residues 1:98); Lane 5: C-terminal His-tagged *Vc*DciA Lasso domain (residues 98:157); Lane 6: Untagged *Vc*DnaB-E259A; Lane 7: Untagged *Vc*DnaB-K234A; Lane 8: N-terminal His-tagged *Vc*DnaB-E259A; Lane 9: Coexpressed *Vc*DnaB untagged + C-terminal His-tagged Full-length *Vc*DciA.

**Supplementary Figure 3.**
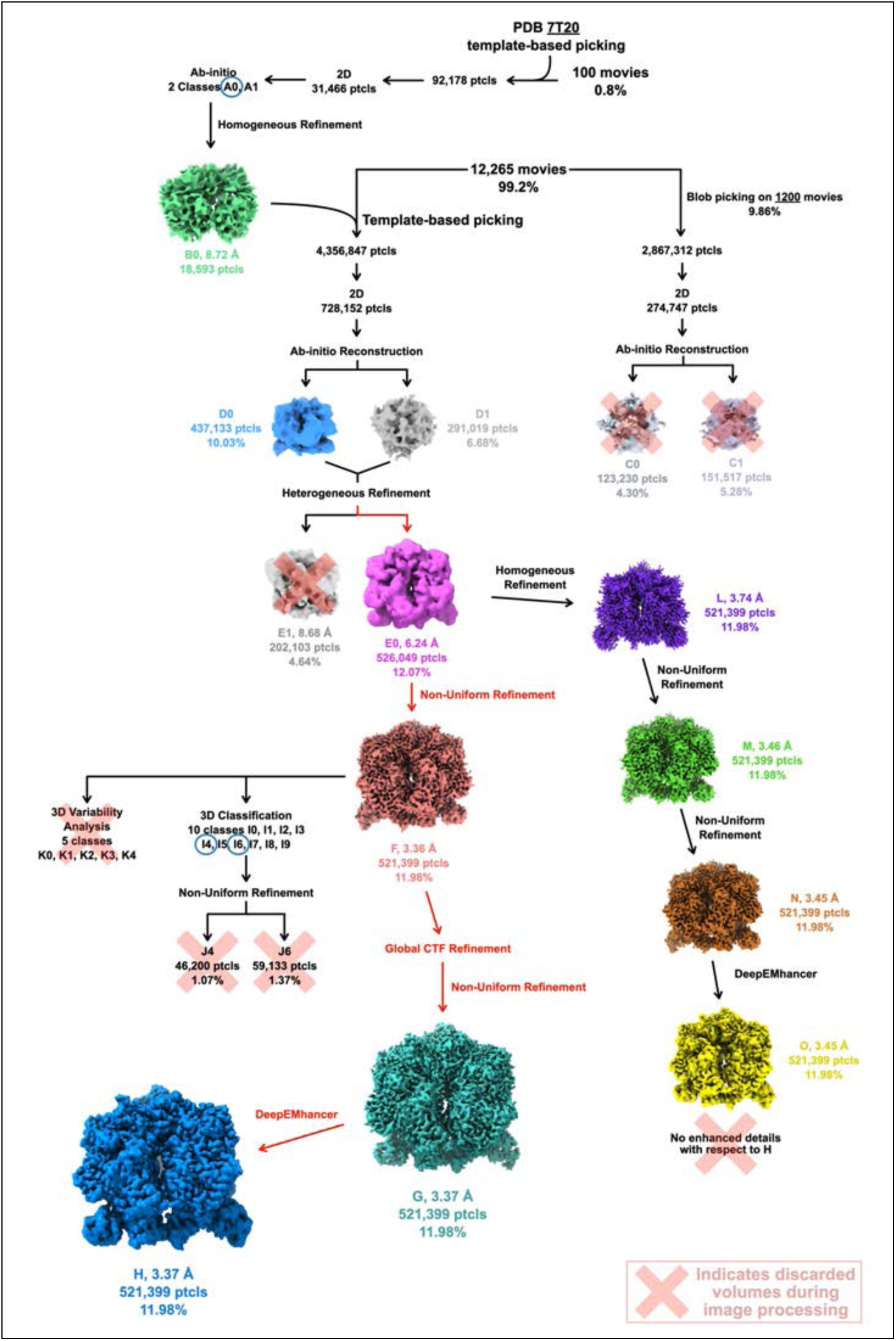
Workflow in the Processing of cryo-EM Data for the *Vc*DnaB-ssDNA-ATPγS complex. The steps in this workflow are described in the Methods section. The percentages reported alongside the volumes are calculated against the number of starting picked particles. The various volumes/maps in the workflow are depicted with unique colors. Map H represents the highest quality map we obtained, against which the atomic model was built and refined. The extended length of ssDNA was built into map E0.

**Supplementary Figure 4.**
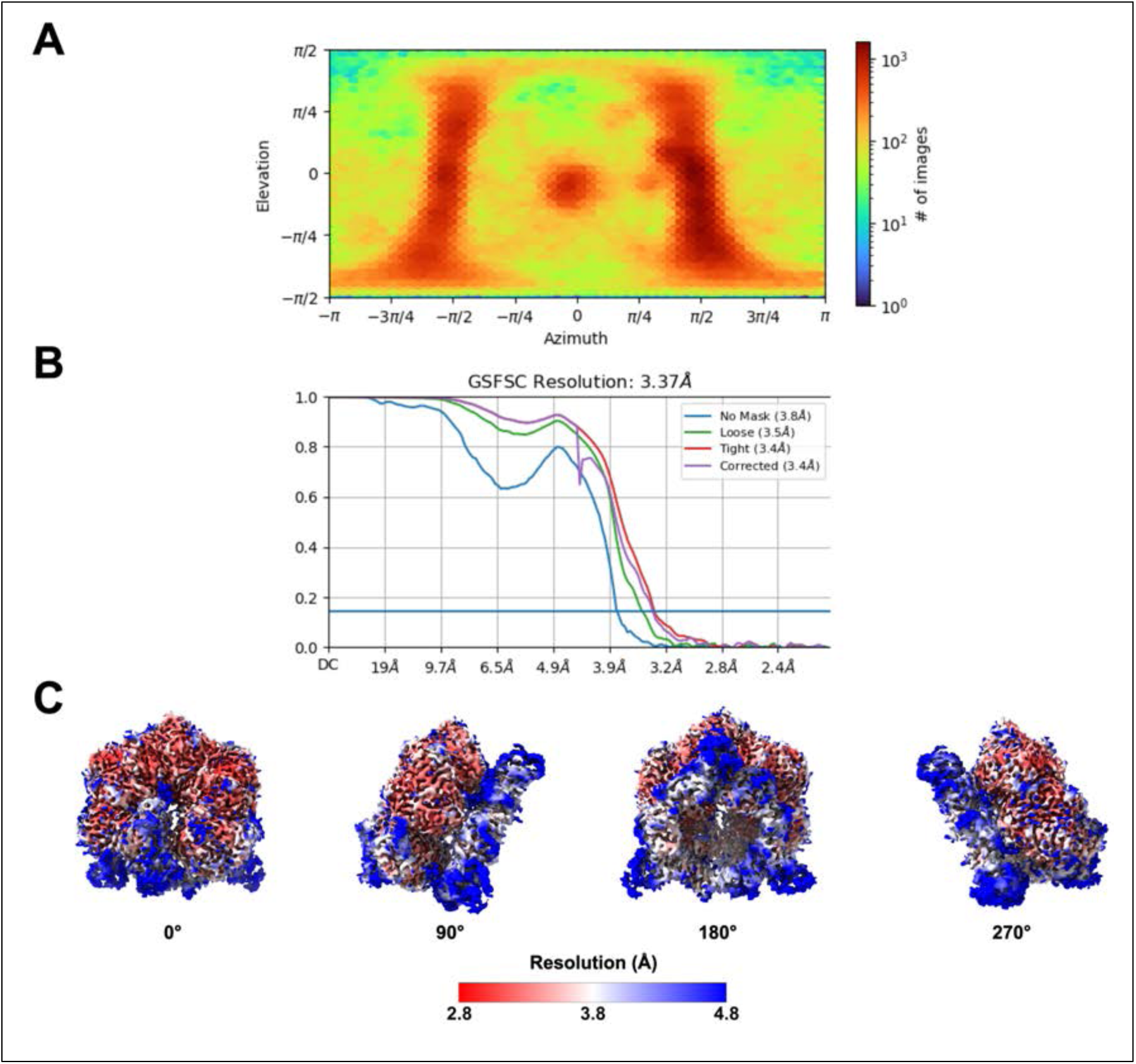
Statistics, Fourier Shell Correlation, and Local Resolution. A) CryoSPARC plot of particles’ azimuthal and elevation angles in the 3.37 Å reconstruction of *Vc*DnaB-ssDNA-ATPγS. The blue-to-red scale represents the number of particles, ranging from 10^2 to 10^^3^. The red clusters represent the preferred orientation of the particles in our data set; however, this distribution of particles did not impact the quality of the map reconstruction. B) Fourier Shell Correlation (FSC) plots taken from the non-uniform refinement calculation that produced the map H. Plots were generated by applying no mask, spherical mask, loose mask, tight mask, and corrected mask on the protein-DNA complex. The corrected tight mask plot shows a resolution of 3.37 Å at an FSC value of 0.143. C) The *Vc*DnaB + ssDNA EM density map is colored (red: 2.8 Å to blue: 4.8 Å) according to the local resolution. The density map is depicted in four poses that differ by 90° rotations along the Y axis.

**Supplementary Figure 5.**
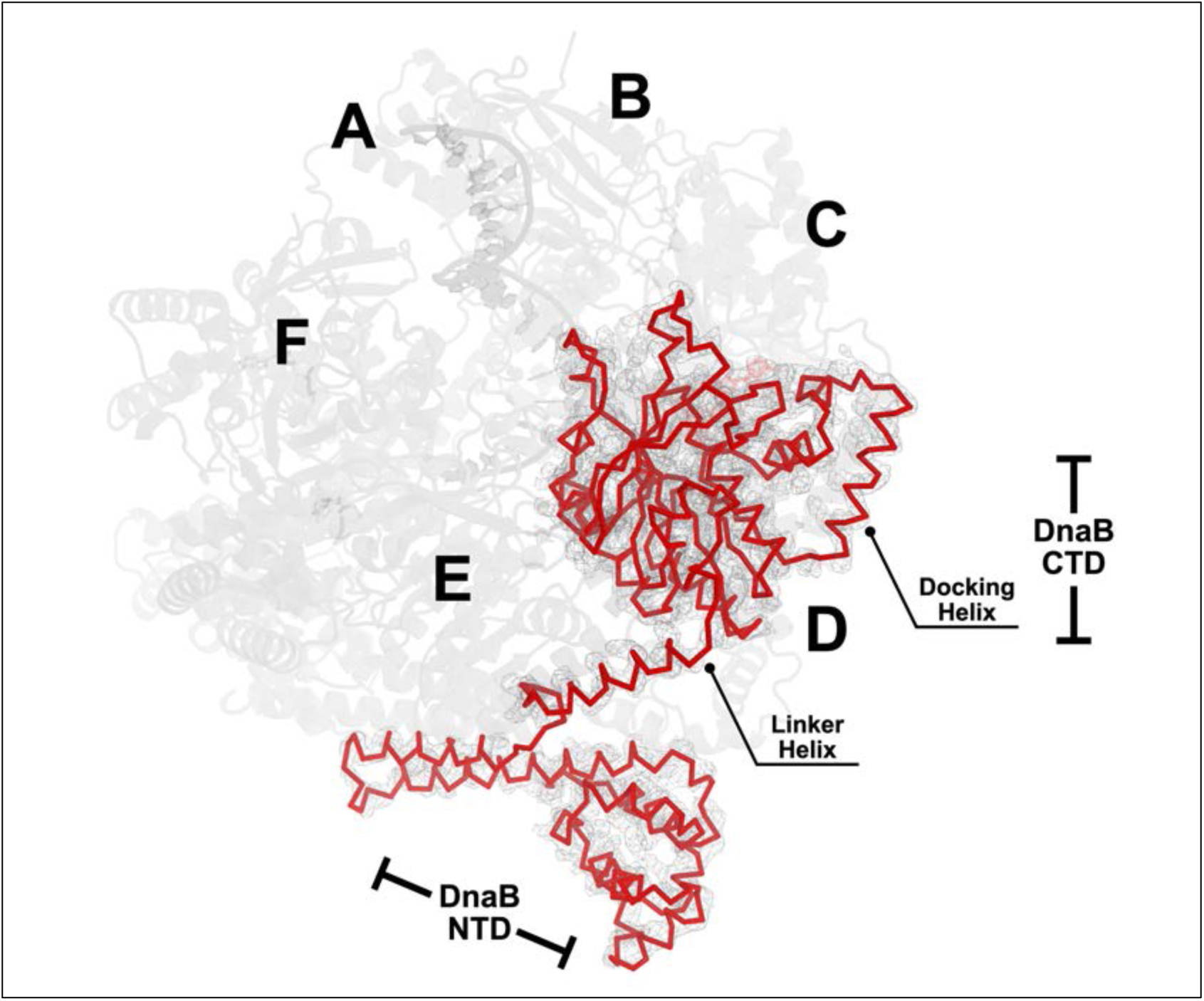
Cryo-EM Density Corresponding to one *Vc*DnaB chain. EM density is drawn as a grey mesh around chain D of the *Vc*DnaB-ssDNA-ATPγS structure. The density is drawn in PyMol using the following parameters: isomesh level: 10 and isomesh carve: 2.3. The *Vc*DnaB chain B ribbon model is superimposed on the EM density. The rest of the hexameric *Vc*DnaB-ssDNA-ATPγS structure is drawn in the cartoon format to provide context. Labels identify DnaB sub-domains and chains.

**Supplementary Figure 6.**
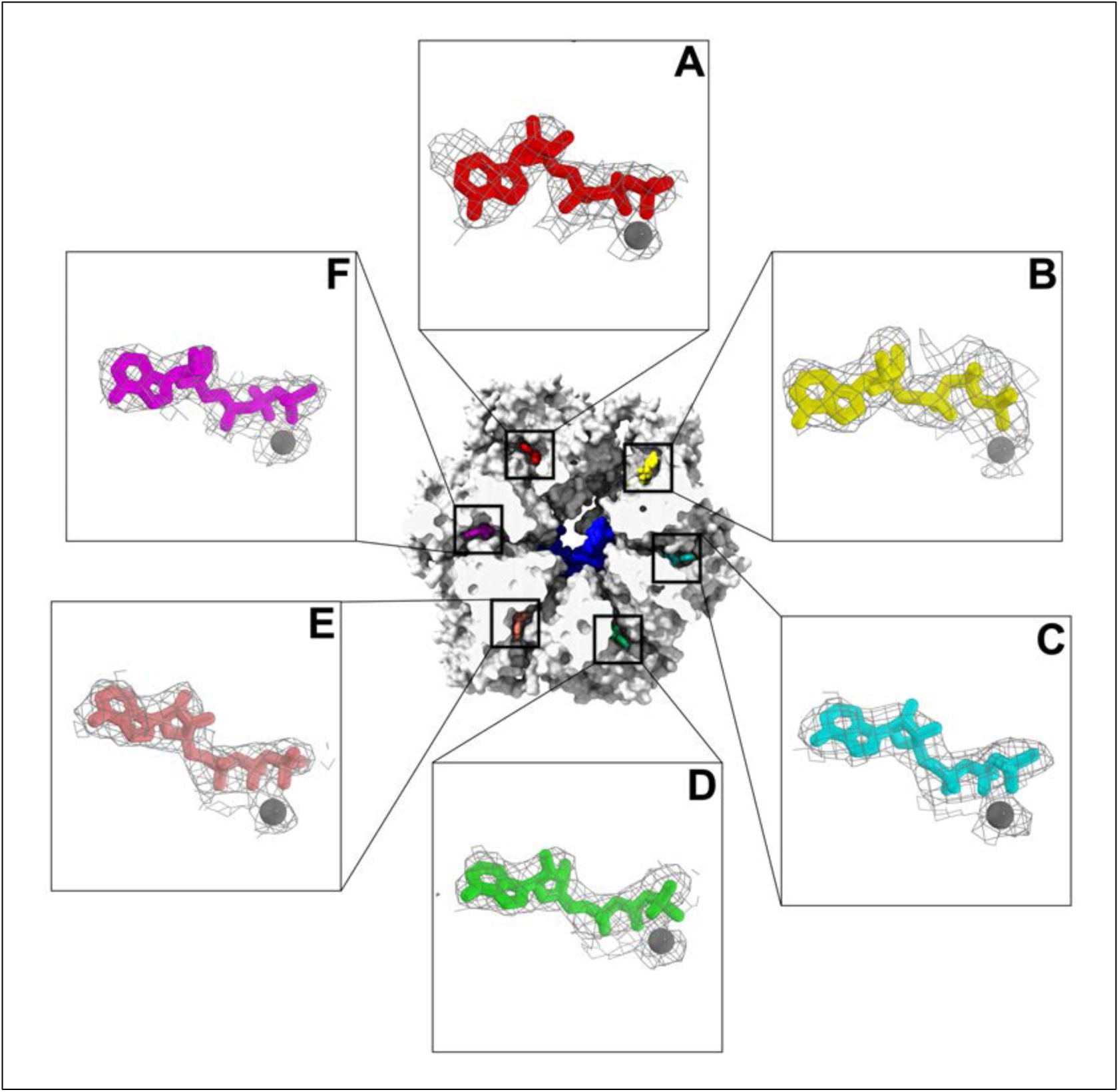
The six nucleotide binding pockets in the *Vc*DnaB-ssDNA structure are filled with ATPγS•Mg^2^. The β and γ phosphates of each ATPγS moiety are coordinated to a Mg^2+^ atom. EM density is drawn as a mesh around each nucleotide. The density is drawn in PyMol using the following parameters: isomesh level chain B: 2; chains A-C-D-E-F: 2, and isomesh carve: chains A-B-C-D-E-F: 5. The ATPγS molecules, each colored uniquely, are drawn in the stick representation, with the Mg^2+^ ion depicted as a grey sphere. The position of the nucleotide pockets is indicated on the surface representation of *Vc*DnaB, sliced to expose the six pockets. ssDNA within DnaB is colored dark blue.

**Supplementary Figure 7.**
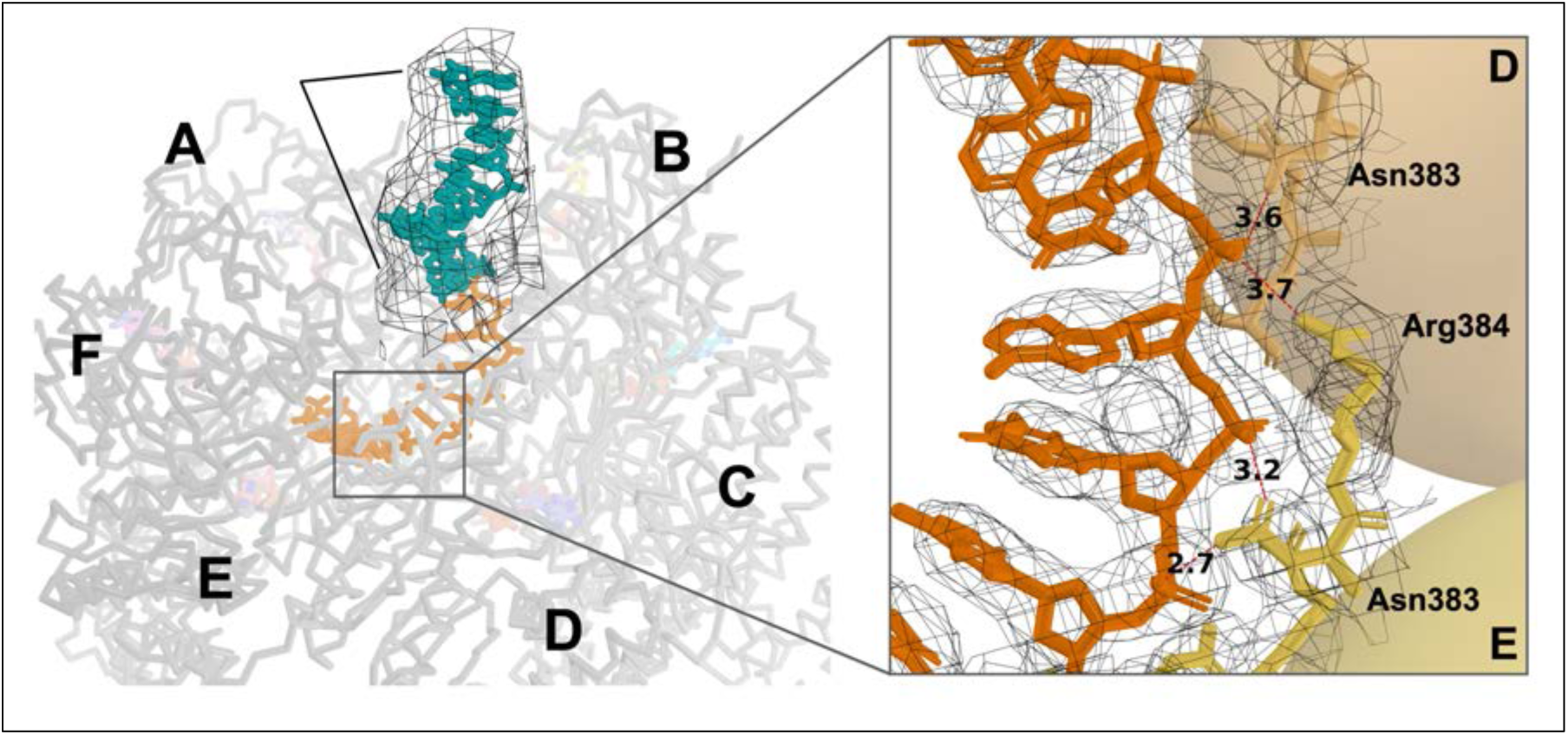
11 additional ssDNA positions outside the confines of *Vc*DnaB are seen in the cryo-EM density map. Close inspection of the 6.24 Å map E0 (Supplementary Methods and Supplementary Figure 3) enabled an ssDNA model to be constructed that was 11 positions longer than that seen in map H. The map into which this segment was built is shown as a mesh (isomesh level: 2.5). The extra positions are colored in teal; the ssDNA built into map H is colored orange. *Vc*DnaB is represented as a grey ribbon, and each monomer is labeled by chain (A-B-C-D-E-F). For reference, the six ATPγS and six Mg^2+^ atoms are displayed as sticks and spheres, respectively, with transparency applied. As noted in the methods, neither map H nor map E0 was sufficiently resolved to permit the assignment of parts of the input 60-mer sequence to the map. As such, the sequence in our final model was built as follows: TCCAGATACACAAAAAAAAAAA. Inset: close-up view of the DnaB-ssDNA interface superimposed on the cryo-EM density. The inset was rotated ∼90° counterclockwise with respect to the main panel and adjusted to align the nucleobases along the vertical axis.

**Supplementary Figure 8.**
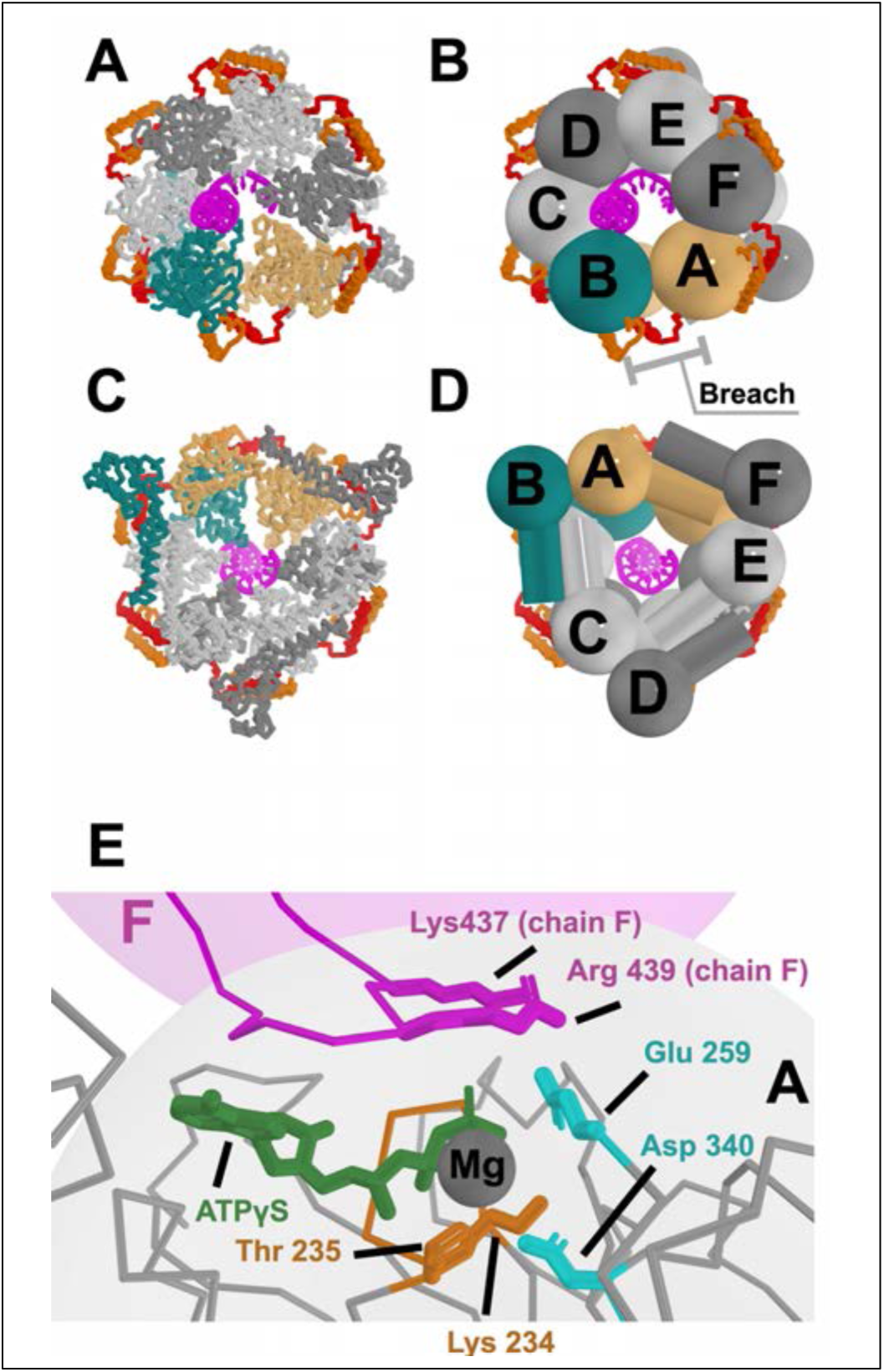
The *Vc*DnaB-ssDNA-ATPγS Complex. The *Vc*DnaB-ssDNA-ATPγS is drawn in a ribbon representation (A or B) or a schematic using a design language wherein the NTDs are depicted as a mushroom shape and the CTDs as spheres (B or D). ssDNA, colored in magenta, is displayed in a cartoon and stick representation. The breach in the CTD layer is between chain A (light orange) and chain B (deep teal); other DnaB subunits are colored in alternating shades of gray. The Docking Helix (DH) and the Linker Helix (LH) are depicted as transparent cylinders colored orange and red, respectively. The pose in panels A and B is along the path of DNA. Panels C and D are in a pose that is rotated approximately 180° from that in panels A and B. E) Closeup of the composite nucleotide catalytic center in *Vc*DnaB at the interface between subunits A (gray shape) and F (magenta shape). ATPγS (green) is bound at a composite site created by the Walker/P-Loop residues Thr 234 and Lys 235 (orange), Walker B residues Asp 340 and Glu 259 (cyan) from chain A, and the β-hairpin arginine finger residues Lys 437 and Arg 439 from chain F. Chains A (grey) and F (magenta) are depicted in the ribbon format.

**Supplementary Figure 9.**
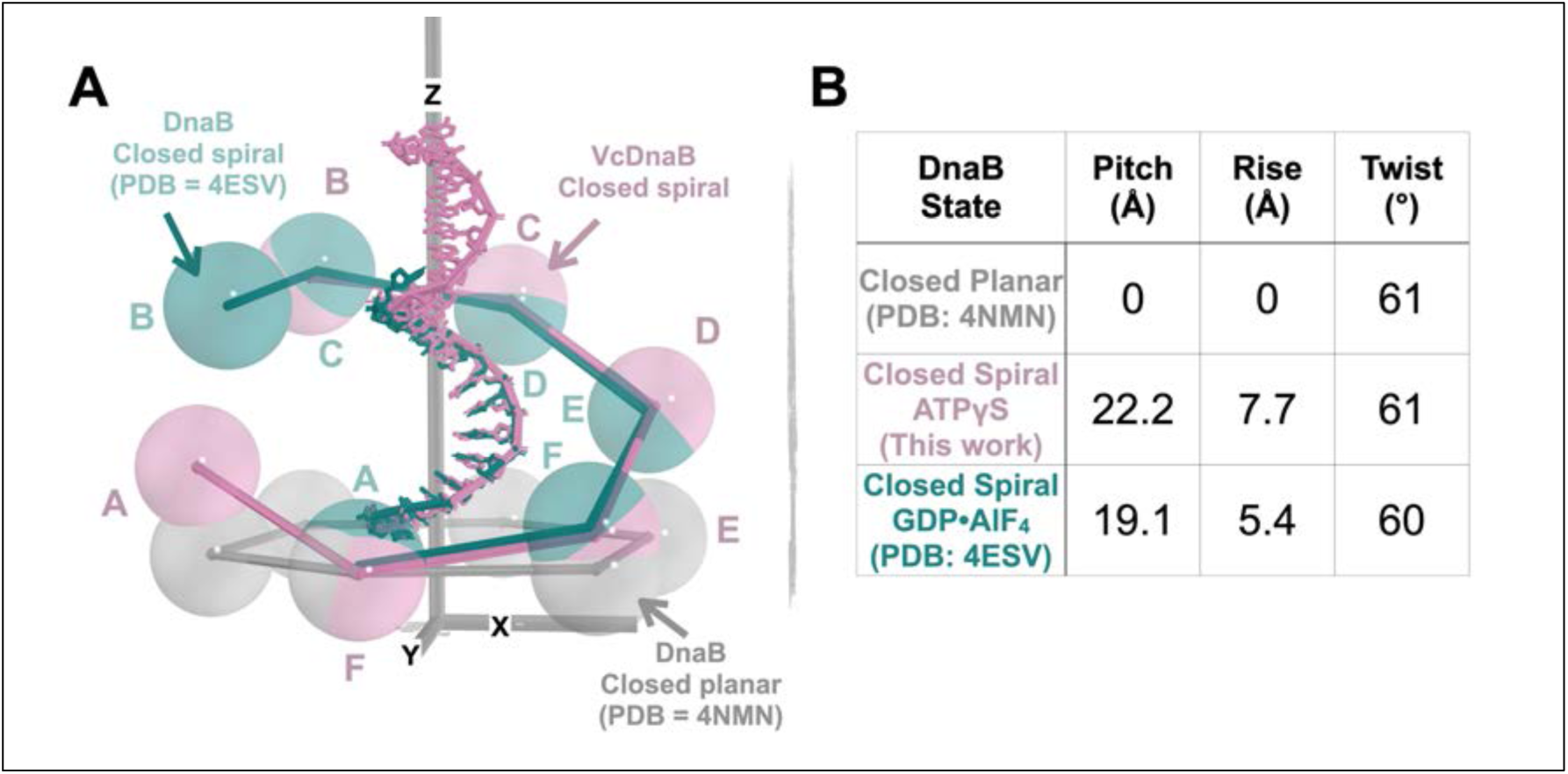
The CTD Tier of DnaB Populates Distinct States as a Function of Nucleotide. A) The CTD tiers are presented as interconnected spheres centered on the indicated DnaB structures. The closed spiral (pink) of the ATPγS *Vc*DnaB CTD layer is displayed against the corresponding closed spiral CTD layer from *Bs*t GDP•ALF_4_ (PDB: 4ESV (12), deep teal) and the closed planar form (PDB: 4NMN (36), grey). The CTD of chain A of the GDP•AlF4 4ESV structure was superimposed onto the CTD of chain A of 4NMN (RMSD = 1.0 Å over 210 Cα atoms). The CTDs of the VcDnaB structure (excluding chain A) were aligned to the CTDs of 4ESV (excluding chain B) by superimposing the ssDNA species (depicted above in the color of the parent structure) in each structure, resulting in an alignment of CTDs with an RMSD = 1.7 Å over 1016 Cα atoms. This alignment positions *Vc*DnaB-ssDNA chain F atop chain A of 4ESV. The 4NMN structure is aligned to the XY plane, with the Z-axis passing through the central chamber. The chains for the CTD spheres corresponding to the 4ESV and *Vc*DnaB structures are labeled in the color of the underlying structure. B) Helical parameters (pitch, rise, and twist) for the specified states of the DnaB CTD.

**Supplementary Figure 10.**
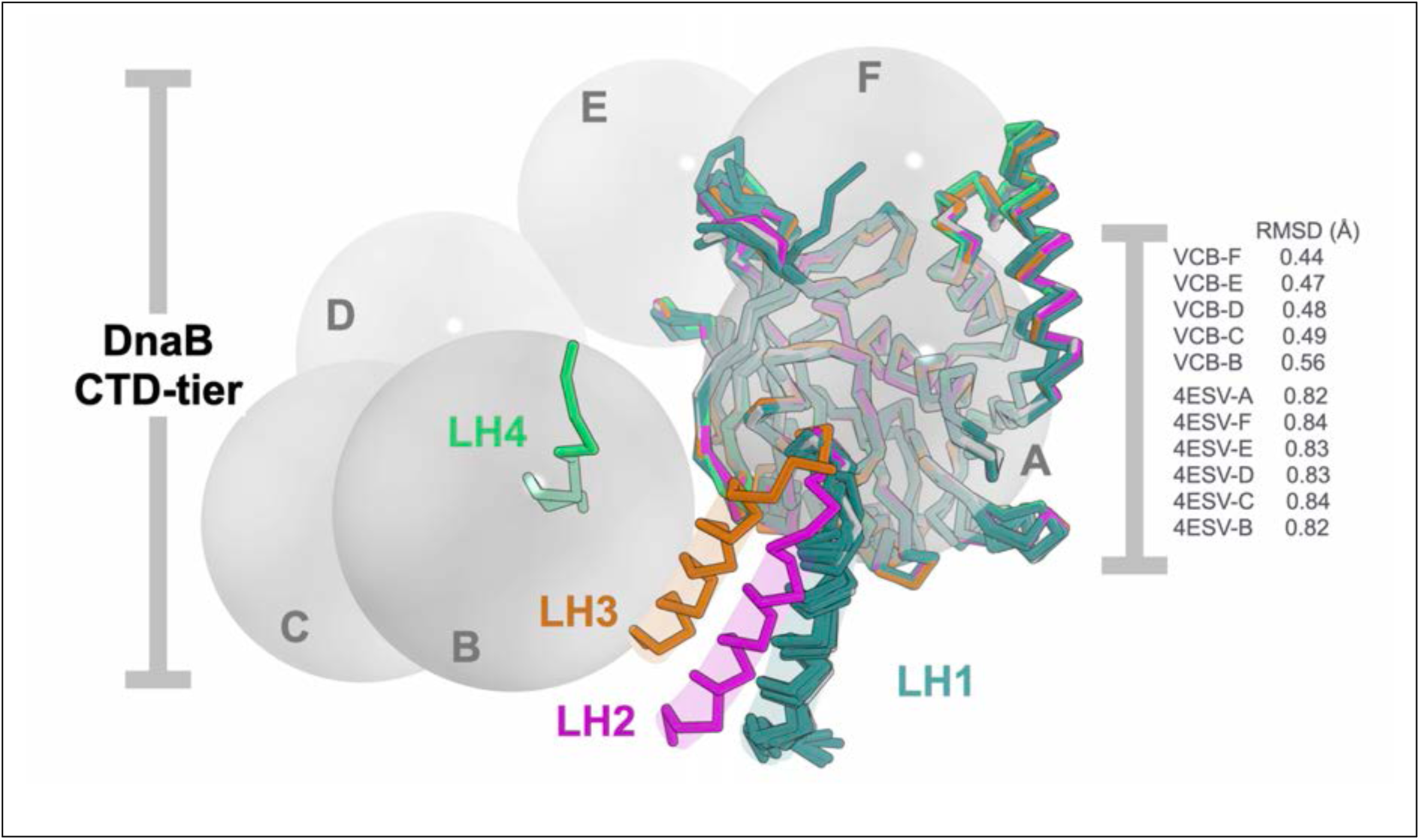
The DnaB Linker Helix (LH) Element is a Moving Part During Translocation on ssDNA. Superposition analysis of the Vc ATPγS and GDP•AlF_4_ structures reveals that the LH element of DnaB occupies at least four positions, termed LH1, LH2, LH3, and LH4. The CTDs (total of 11) of the *Vc* and Bst structures were superimposed on residues 204:465 of *Vc* chain A (residues corresponding to the LH 178:205 or DH 287:304 elements were omitted from the calculation). The four positions of the LH are colored in deep teal (LH1), magenta (LH2), orange (LH3), and lime-green (LH4). Except for chains A (LH3) and F (LH2), the *Vc* DnaB chains populate the LH1 position. Chain A of the Bst-4ESV structure occupies the LH4 position, while the others are observed in the LH1 position. The CTD layer of the *Vc* structure is shown as labeled spheres drawn at the centers of mass of each CTD. The superimposed CTDs are drawn in the ribbon representation, and the transparent cylinder indicates the LH.

**Supplementary Figure 11.**
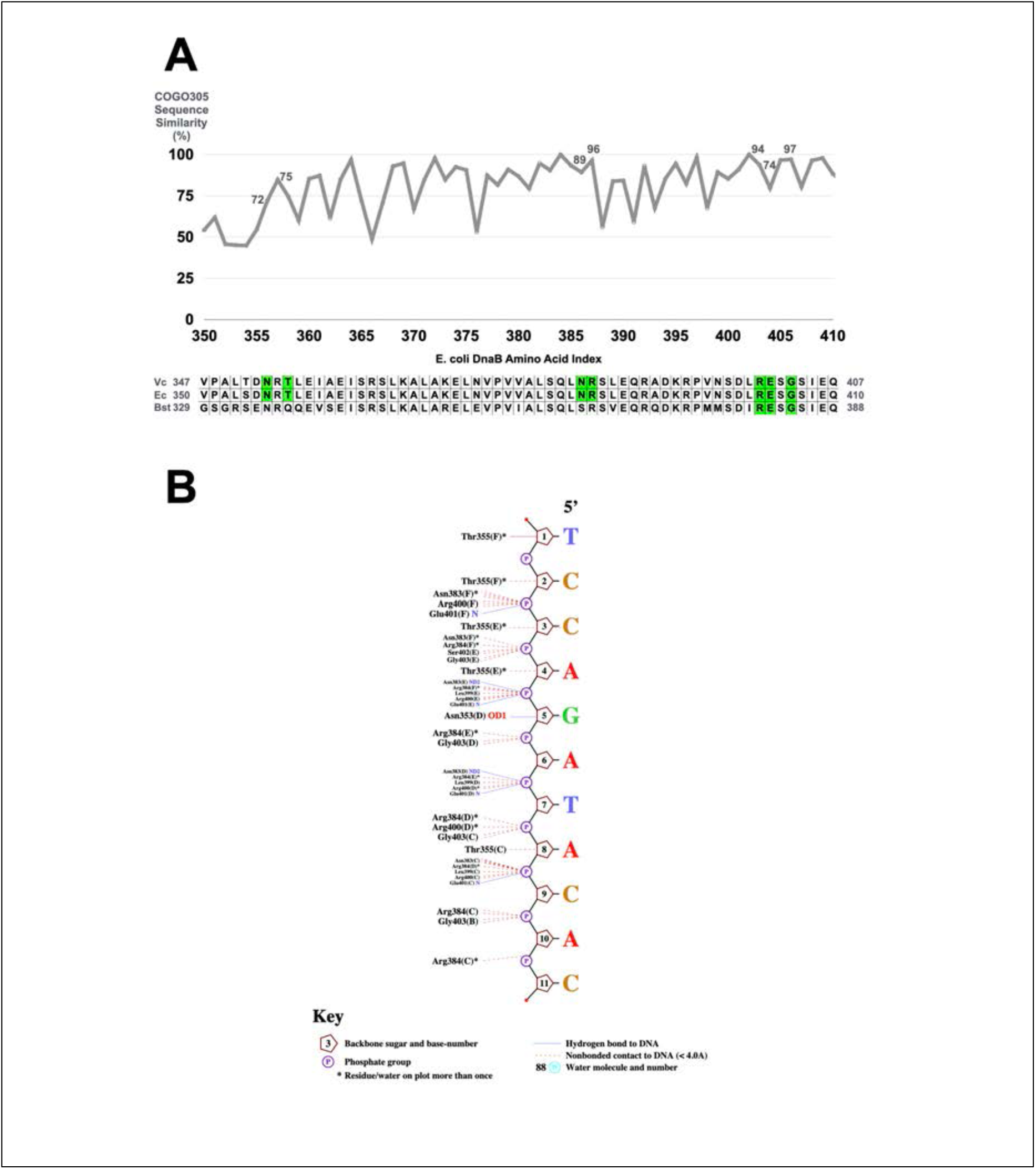
Protein–ssDNA interactions in the *Vc*DnaB-ssDNA complex. A) Sequence identity is plotted against the amino acid index of a portion of the sequence alignment derived from the DnaB COG0305 conserved protein domain family (100 sequences). Seven *Vc*DnaB residues are in contact with ssDNA, though not every subunit uses each residue. These are N353, T355, N383, R384, R400, E401, and G403 (highlighted in green). The sequence alignment teaches that these residues are conserved in the COG0305 DnaB-family. The conservation value for each contact residue is listed in the plot. The residues used by *E. coli* (PDB: 7T20 (17)) and Bst (PDB: 4ESV(12)) are also highlighted. B) *Vc*DnaB-ssDNA contacts are depicted schematically as drawn by Nucplot (37).

**Supplementary Figure 12.**
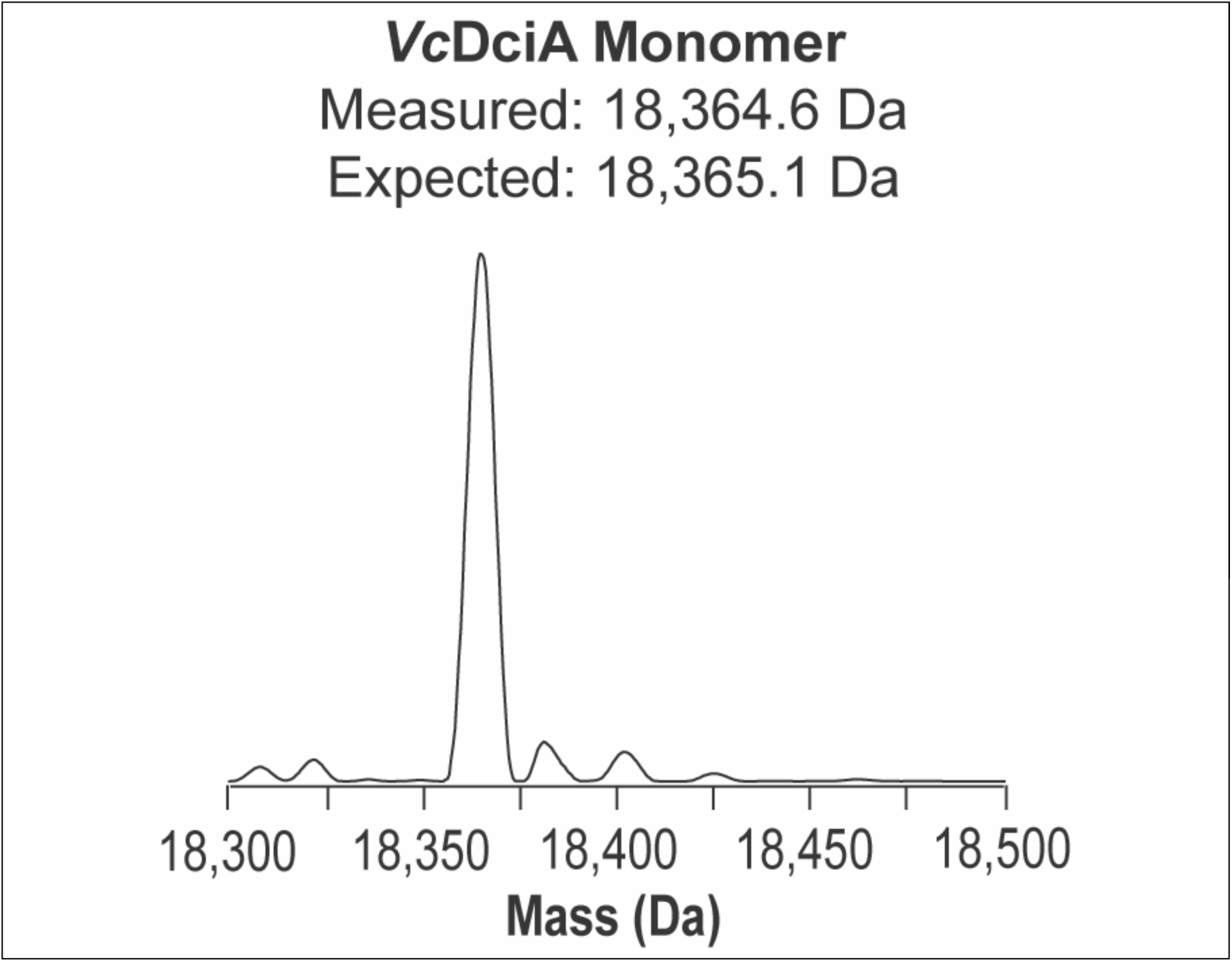
Isolated *Vc*DciA is a monomer. Deconvolved nMS spectra of *Vc*DnaB.

**Supplementary Figure 13.**
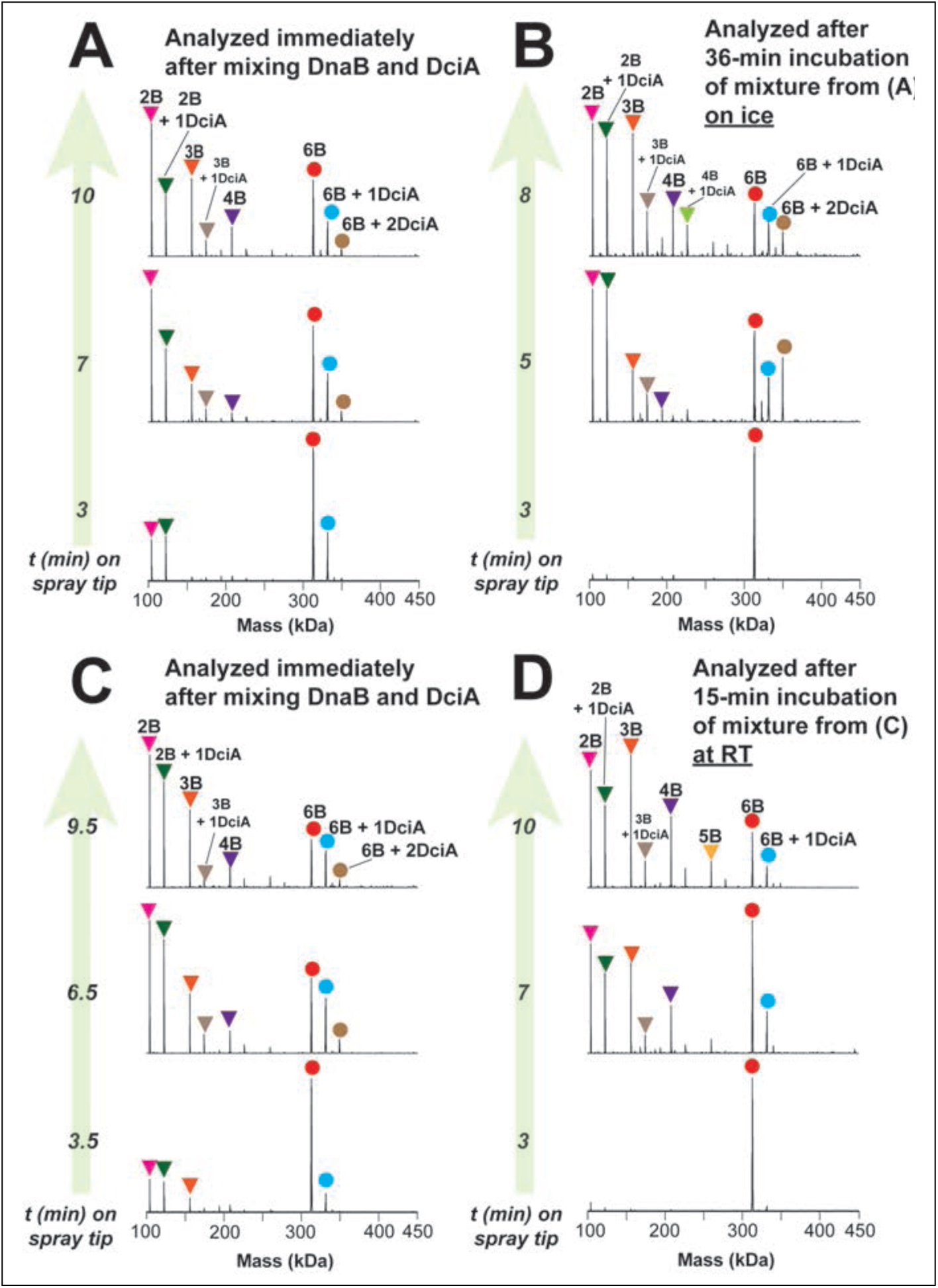
Disassembly of *Vc*DnaB upon incubation with *Vc*DciA during nMS analysis. *Vc*DnaB was mixed with *Vc*DciA, and an aliquot of the mixture was loaded immediately on the electrospray tip for nMS analysis (A and C), and the rest of the mix was either incubated on ice for 36 min (B) or at room temperature for 13 min (D) before nMS characterization. The deconvolved spectra at specific time points during the nMS analysis (∼100 scans averaged) and the corresponding peak assignments are shown for each sample. The overall delay time from loading the sample into the tip, setting up the electrospray, and initiating nMS data acquisition is typically 2 min. The sample mixture comprised 10 μM DnaB and 7 μM DciA in 10 μM ATP, 0.5 mM magnesium acetate, 500 mM ammonium acetate, and 0.01% Tween-20. The peaks for DnaB_6_ and DnaB_6_ + DciA_1_ predominate during the first 1-3 minutes of initiating the electrospray, and then peaks with increasing intensities for DnaB_2_, DnaB_2_ + DciA_1,_ and other DnaB subcomplexes appear, indicating the progressive disassembly of DnaB_6_. Analysis of the rest of the sample incubated on ice or at room temperature (RT) resulted in similar time-resolved mass spectrometry (MS) profiles, suggesting that DnaB_6_ dissociation only occurs during electrospray. Overall, the results indicate that DciA binding can destabilize DnaB_6,_ and this destabilization is intensified during the electrospray process, leading to DnaB_6_ disassembly.

**Supplementary Figure 14.**
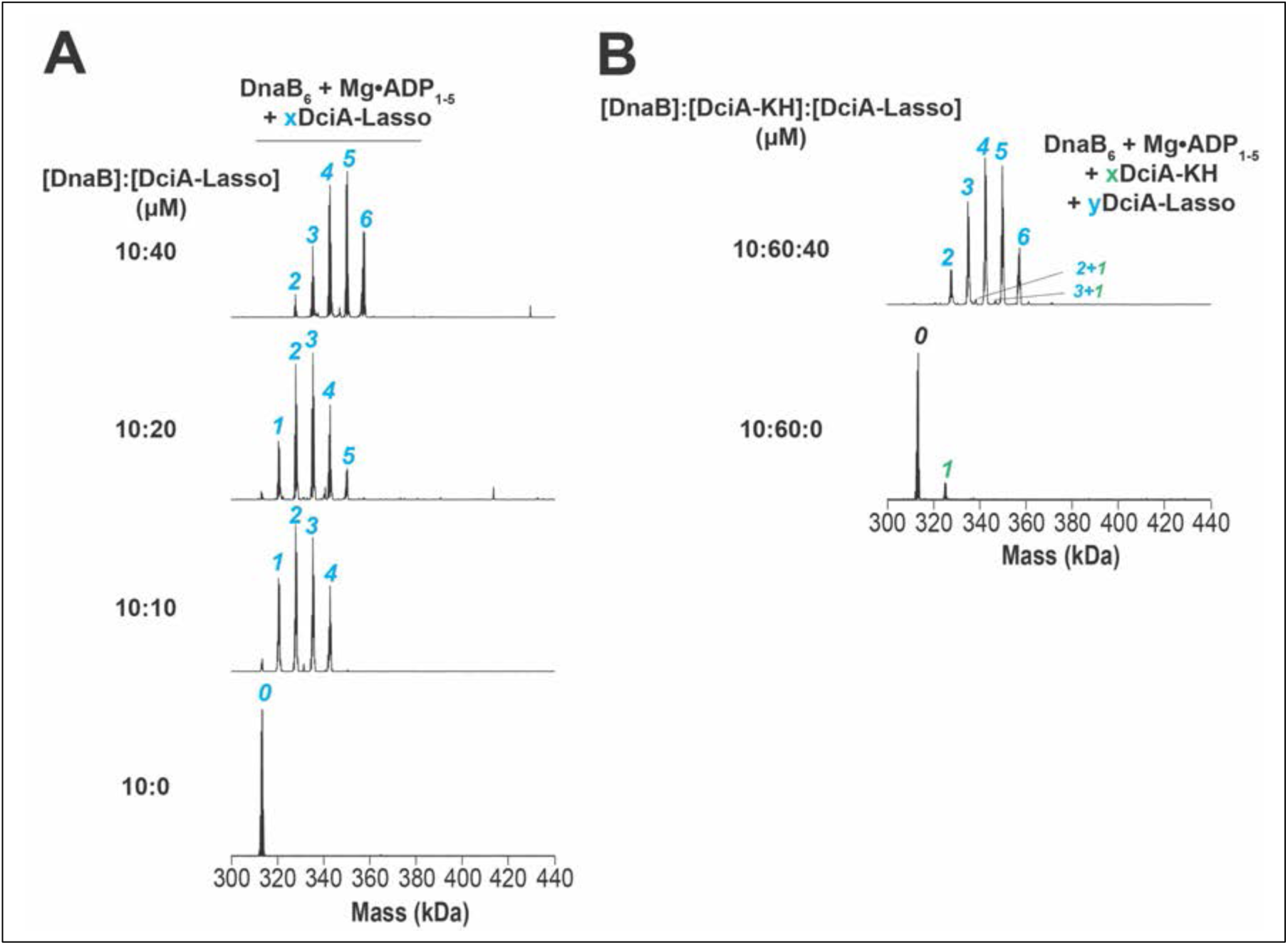
Stoichiometry of binding the *Vc*DciA KH domain, the *Vc*DciA Lasso, and ssDNA to *Vc*DnaB by nMS. Deconvolved nMS spectra of (**A**) DnaB titrated with increasing concentrations of DciA-Lasso, and (**B**) DnaB incubated with an excess of DciA’s KH domain in the absence or presence of DciA’s Lasso domain. The expected masses of *Vc*DnaB_6_ = 311.5 kDa, DciA-KH = 11.8 kDa, DciA Lasso = 7.5 kDa, and ssDNA = 32.5 kDa. The colored numbers and Roman letters ‘X’ and ‘Y’ indicate the number of DciA-Lasso domains bound (A) or DciA-KH and Lasso bound (B).

**Supplementary Figure 15.**
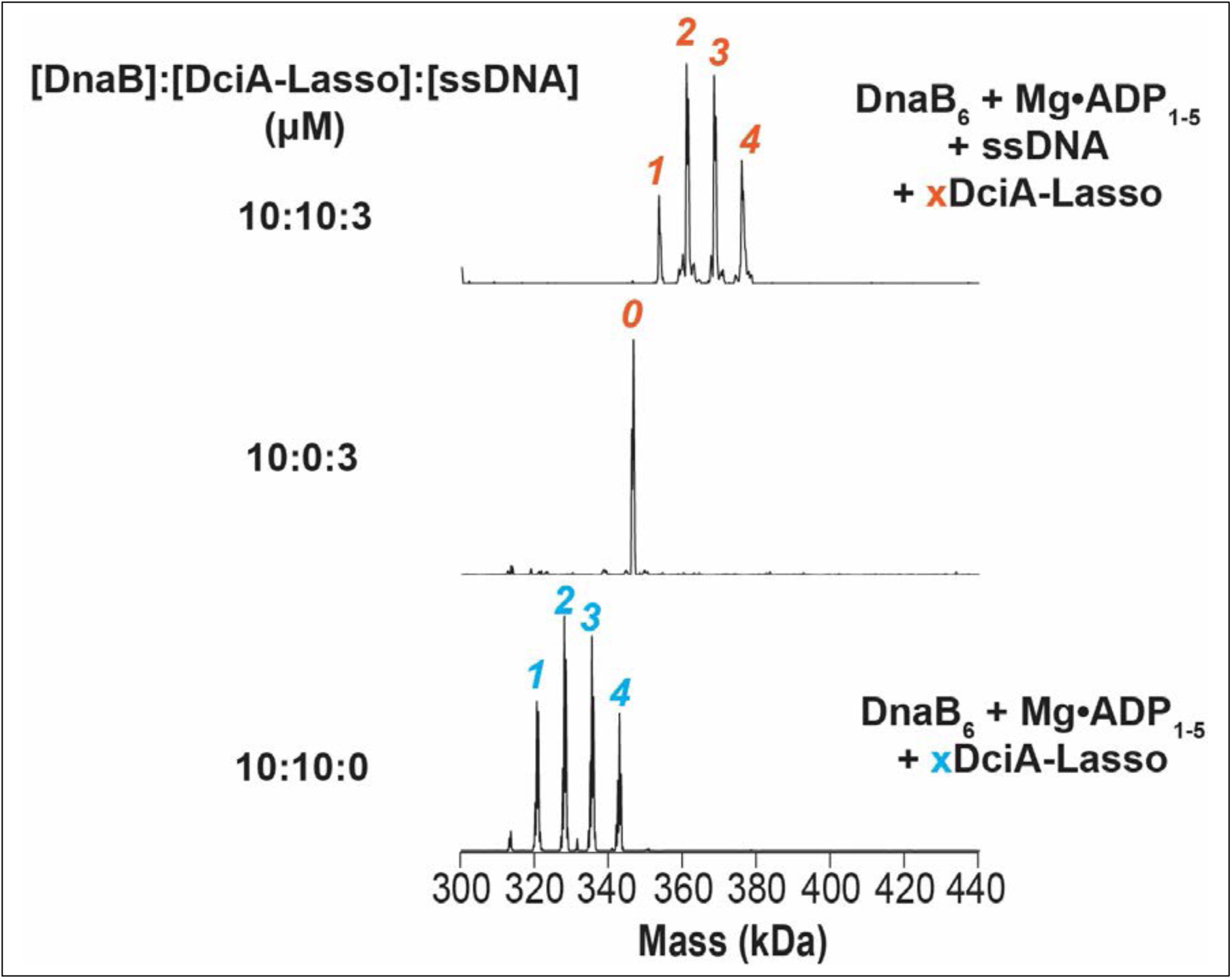
The *Vc* Lasso domain forms complexes with *Vc*DnaB and ssDNA. Up to 4 *Vc* Lasso domains are found in *Vc*DnaB complexes, whether ssDNA in the absence/presence of ssDNA. Deconvolved nMS spectra of DnaB and ssDNA (O4-*Vc*F) incubated with and without an equimolar concentration of DciA-Lasso. All samples are prepared in 10 µM ATP, 500 µM magnesium acetate, 500 mM ammonium acetate, and 0.01% Tween-20 before nMS analysis. Spectra were collected at the indicated DnaB, DciA-Lasso, and ssDNA ratios. The blue and red Roman letter ‘X’ indicates the number of copies of *Vc*DciA revealed by nMS for a given sample. The expected masses are DnaB_6_ = 311.5 kDa, DciA-KH = 11.8 kDa, DciA Lasso = 7.5 kDa, and ssDNA = 32.5 kDa.

**Supplementary Figure 16.**
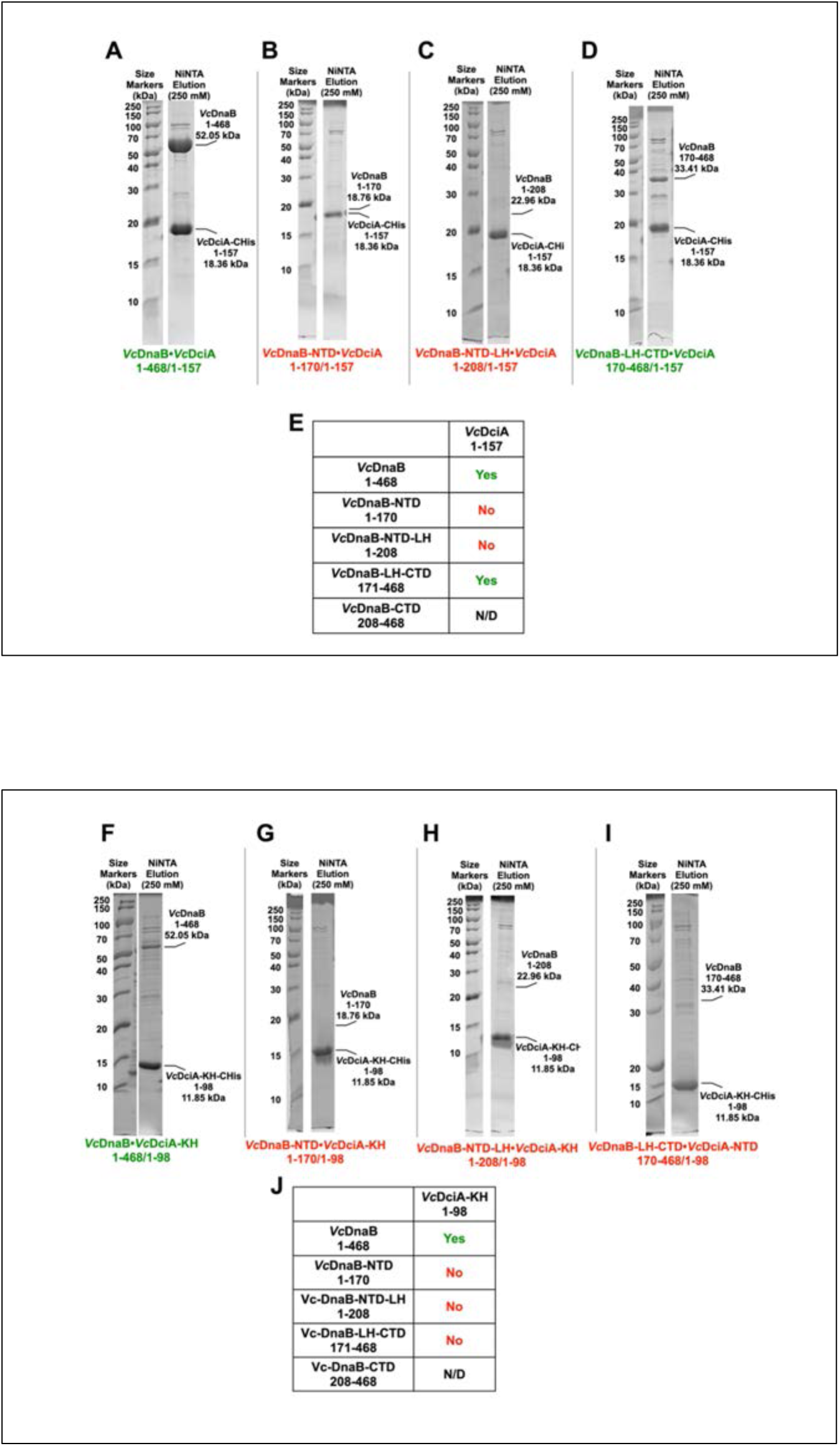
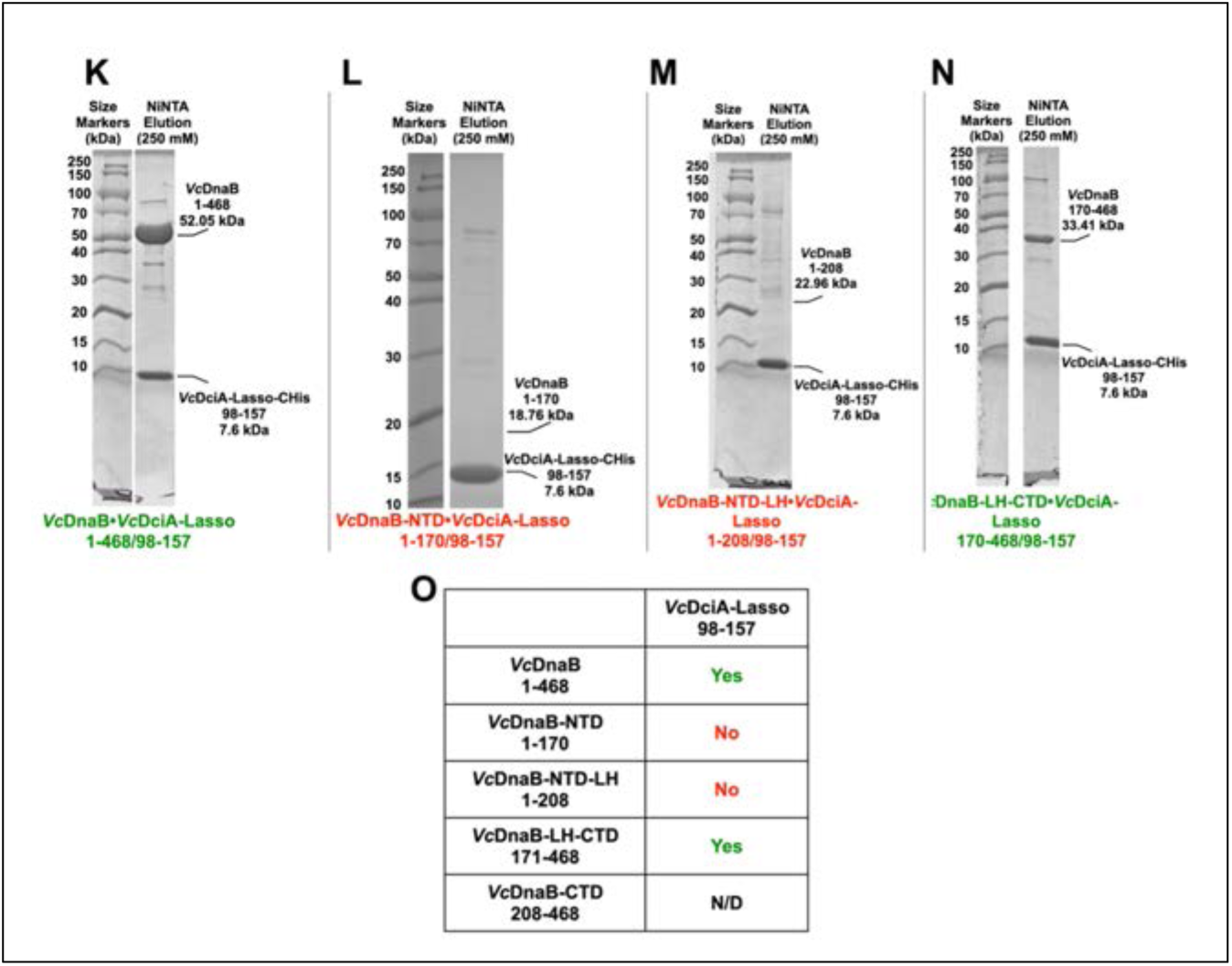
*Vc*DciA binds tightly to *Vc*DnaB, and the *Vc*DciA*-* Lasso domain prefers the LH-CTD domain. Composite image of 12 NiNTA ‘pull down’ assay between *Vc*DciA (full-length: 1:157; KH: 1:98; Lasso: 98:157) and DnaB (full-length: 1:468; NTD: 1:170; NTD-LH: 1:208; LH-CTD: 171:468; CTD: 208:468) constructs. A-D) Full-length *Vc*DciA and various *Vc*DnaB constructs. E) Summary of binding data from panels A-D; the *Vc*DnaB CTD (residues 208:468) could not be stably expressed. F-I) *Vc*DciA KH domain (1:98) and various *Vc*DnaB constructs. J) Summary of binding data from panels F-I. K-N) *Vc*DciA Lasso domain (98:157) and various *Vc*DnaB constructs. O) Summary of binding data from panels K-N. The green text indicated a NiNTA pull-down scored as interacting, while the red text indicated that no interaction was observed. These data show that full-length *Vc*DciA binds to full-length *Vc*DnaB and that the Lasso domain binds to the LH-CTD, likely at the DH-LH interface, as with other helicase loaders (4, 7–9, 13, 15).

**Supplementary Figure 17.**
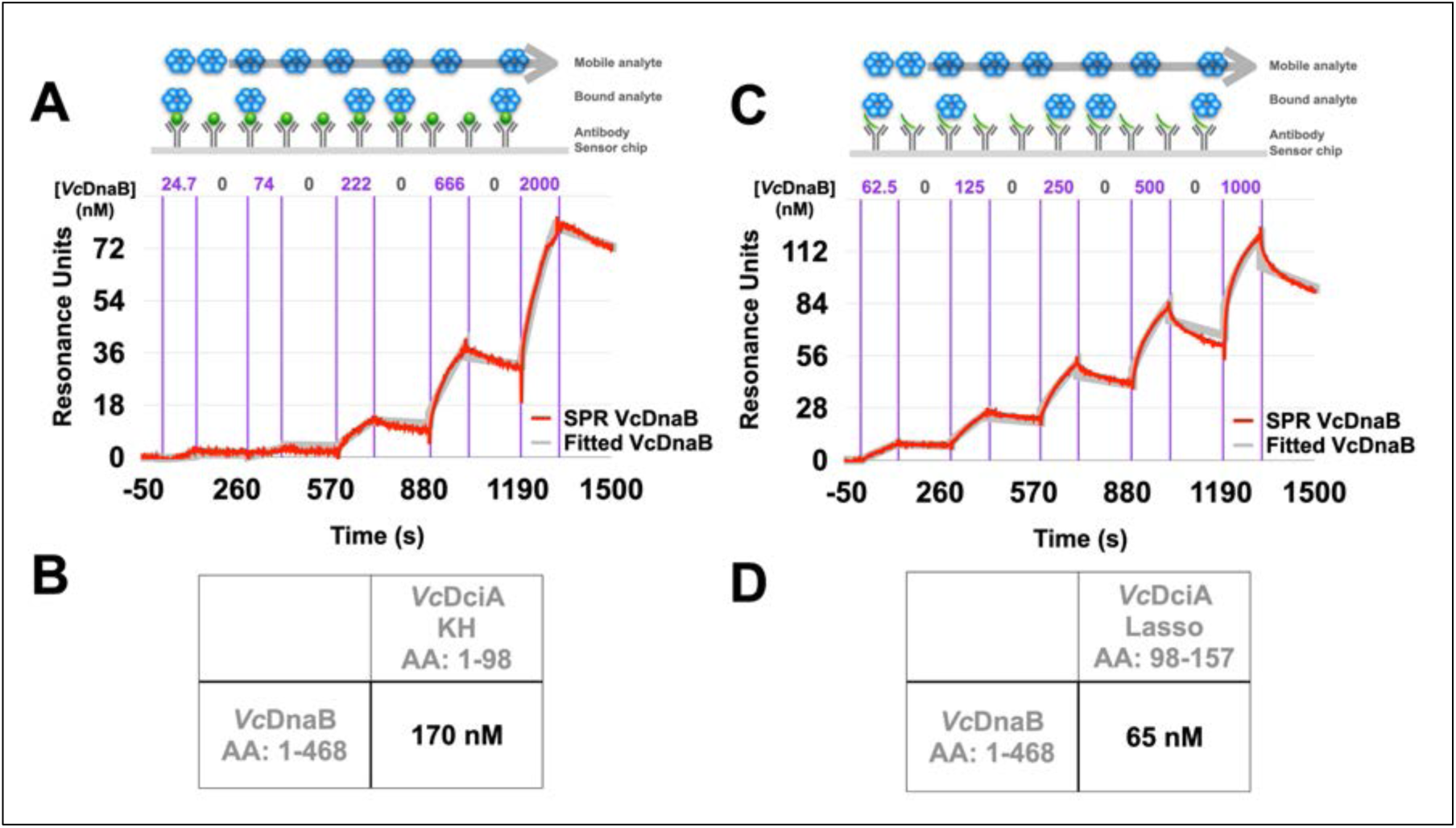
The *Vc*DciA KH and Lasso Domain Retain Affinity to *Vc*DnaB, albeit with a Higher K_D_. A) On and off rates with *Vc*DnaB at incremental concentrations (24.7, 74, 222, 666, 2000 nM) were measured by immobilizing cognate C-terminal His-tagged *Vc*DciA-KH to the SPR sensor chip. The single-cycle kinetic analyses through SPR revealed how DciA’s KH domain exhibited a decreased affinity to *Vc*DnaB. The *Vc*DciA KH domain displays a K_D_ = 170 nM (Chi^2^ = 1.82; R_max_ = 135.2), representing a ∼300-fold reduction compared to full-length (A, B). C) Same as panel A except that slightly different incremental concentrations (62.5, 125, 250, 500, 1000 nM) were used. D) The *Vc*DciA Lasso domain features a K_D_ of 65 nM (Chi^2^ = 5.89; R_max_ = 126.3). The 170 nM and 65 nM K_D_ measured for the KH and Lasso domains contrast with the 0.58 nM value measured for full-length DciA. The SPR experiments are diagrammed schematically above each trace. The experimental data were fitted as described in the methods. The raw SPR signal is shown in red, and the fitted model is depicted in grey.

**Supplementary Figure 18.**
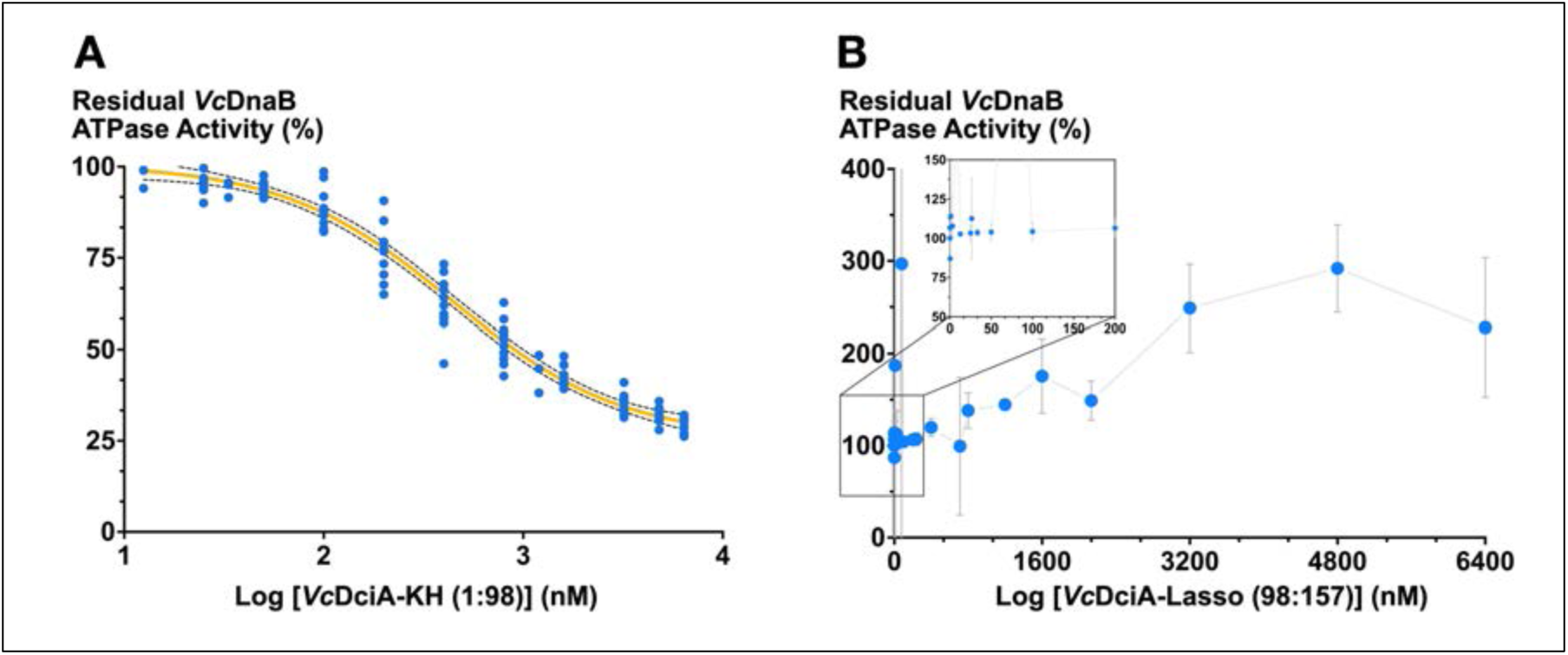
The *Vc*DciA KH, but not the Lasso, domain partially retains the ATPase suppression activity. A) The IC_50_ calculation described in Figure 7B was applied to the *Vc*DciA KH and Lasso domains. The *Vc*DciA KH domain represses *Vc*DnaB ATPase activity with an IC_50_ of 443 nM (R^2^ = 0.9693; 95% confidence interval: 380.5 to 520.0 nM), representing a 16-fold lower potency than the full-length loader. The blue dots represent individual measured ATPase rates plotted on a log scale. A non-linear fit (light-orange line), calculated using the Sigmoidal, 4PL model as implemented in PRISM, is superimposed on the experimental data. The dashed lines represent the 95% confidence interval (CI). These experiments encompassed four independent protein batches. B) The same analysis applied to the *Vc*DciA Lasso domain revealed no effect on *Vc*DnaB’s ATPase activity at low concentrations (0.11 – 200 nM). However, the titration of a significant excess of the Lasso domain (800 – 6400 nM) revealed a 2.8x stimulation of DnaB’s ATPase activity at 4800 nM *Vc*DciA. The measured ATPase rates (blue dots) in the inset are plotted on a linear scale and connected with a grey line.

**Supplementary Figure 19.**
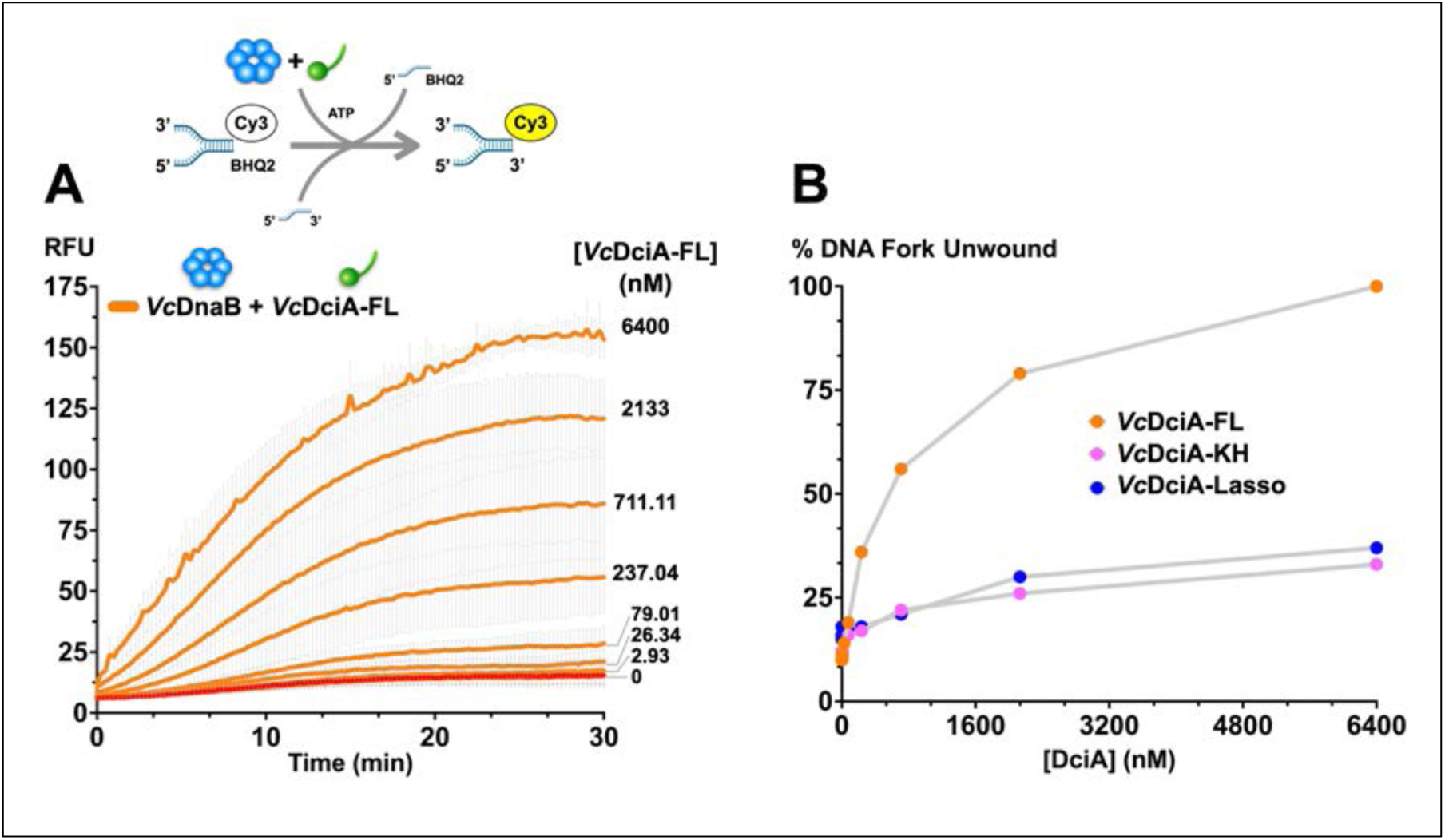
*Vc*DciA stimulates the DNA unwinding activity of *Vc*DnaB in a concentration-dependent manner. A) The unwinding experiments employed a FRET-based 5ʹ- and 3ʹ-tailed replication fork mimic, which is diagrammed schematically above. A time course of the evolution of fluorescence at 570 nm, due to relief of quenching at the Cy3/BHQ2 center during unwinding, is shown for unwinding experiments where the *Vc*DnaB concentration was held fixed at 200 nM, and the *Vc*DciA concentration varied from 0.11 to 6400 nM (orange curves). Shown in red is *Vc*DnaB’s unwinding when *Vc*DciA was absent. The data indicate a 10-fold increase in unwinding. B) The data in panel A are plotted as a function of %DNA unwound, where the highest RFU value was considered 100% unwound. By this measure, the *Vc*DciA-KH (pink) and *Vc*DciA-Lasso (blue) domains exhibit 30% of the activity of full-length VcDciA. Control measurements lacking *Vc*DciA showed no unwinding, as was observed with the bubble substrate (Figure 8 and Supplementary Figure 25).

**Supplementary Figure 20.**
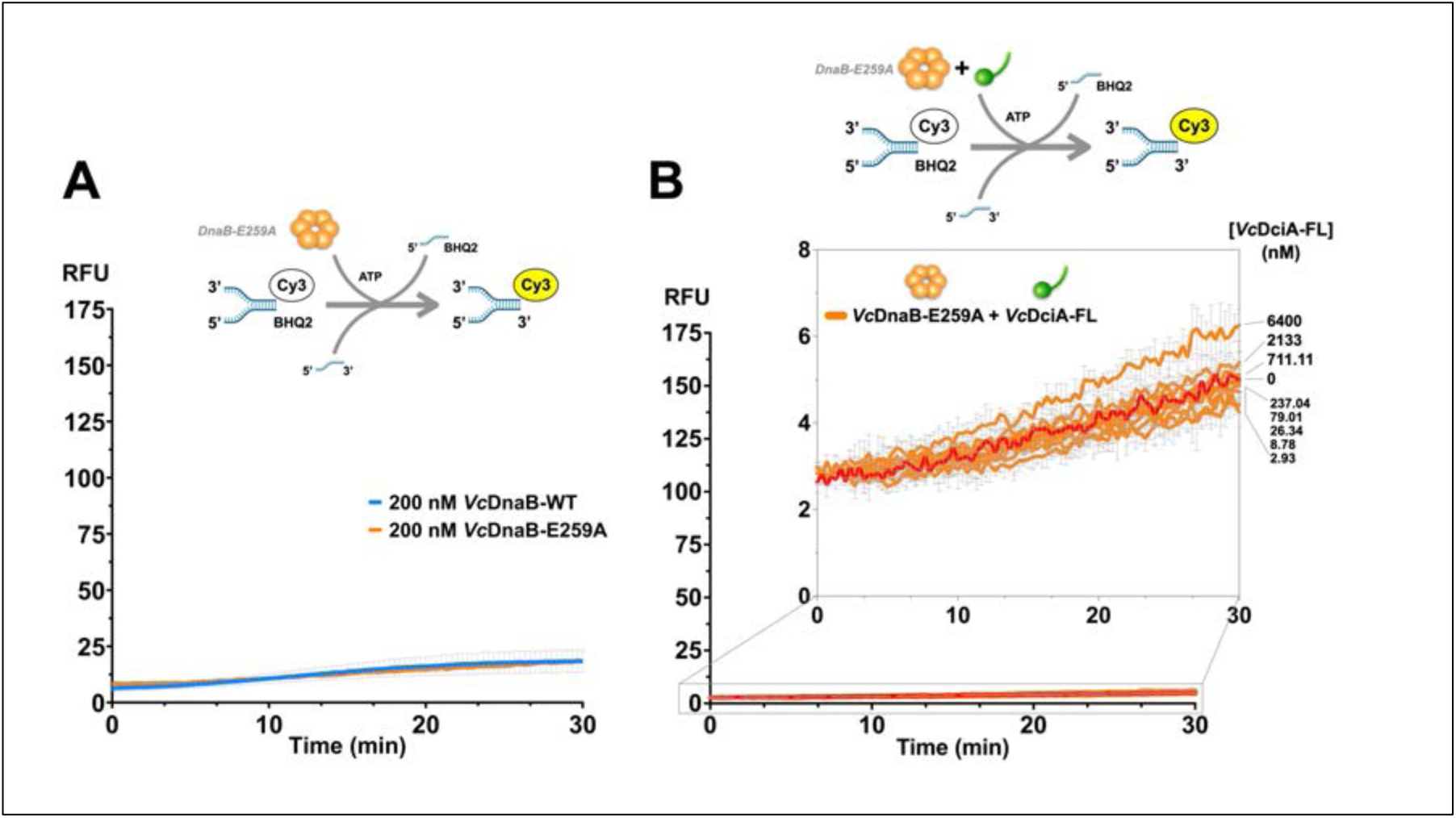
The *Vc*DnaB Walker B glutamate (E259A) mutation Significantly Reduces DNA unwinding activity and *Vc*DciA-stimulated unwinding. The FRET-based DNA unwinding experiments utilized a 5ʹ- and 3ʹ-tailed replication fork mimic, as diagrammed schematically above each panel. A) The time course of the 570 nm fluorescence signal due to the unwinding of the DNA substrate is plotted for wild-type (blue) and E259A mutant (orange) DnaB. In this experiment, the *Vc*DnaB concentration was held at 200 nM. The blue line represents the average of 27 replicates measured from 200 nM *Vc*DnaB-WT. Both wild-type and the E259A mutation show a slight basal fluorescence signal increase in the DNA fork unwinding assay. Although the E259A mutant behaves like wild-type in the fork unwinding assay, adding *Vc*DciA (panel B) does not stimulate it as seen for wild-type (Supplementary Figure 19). B) The time course of the 570 nm fluorescence signal arising from titration of *Vc*DciA (0.11-6400 nM) into an assay containing the Walker B *Vc*DnaB-E259A mutant. Essentially, no unwinding activity was recovered. The inset shows the same data but expands a section of the plot.

**Supplementary Figure 21.**
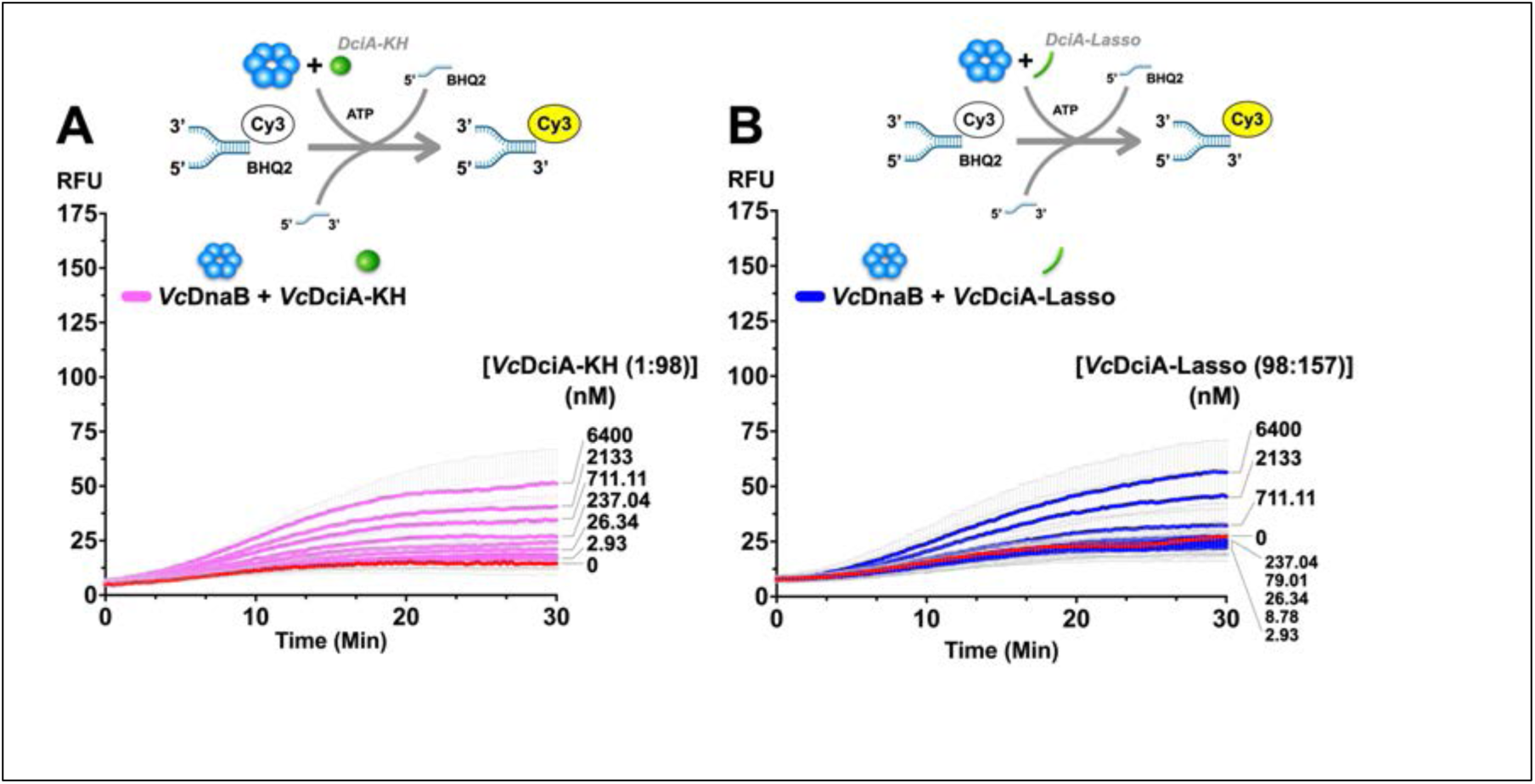
The Isolated KH and Lasso Domains Retain 30% of the Potency of Full-length VcDciA in Stimulating Unwinding by VcDciA. The FRET-based DNA unwinding experiments utilized a 5ʹ- and 3ʹ-tailed replication fork mimic, as diagrammed schematically above each panel. Titration of *Vc*DciA-KH (pink, A) or DciA-Lasso (blue) domains into the assay stimulated DnaB’s unwinding activity by only 30% compared to full-length DciA. The titration of the appropriate DciA-construct encompassed concentrations from 0.11 to 6400 nM. At most, the isolated KH and Lasso domains stimulated DNA unwinding by 3.3-fold compared to a sample lacking DciA. In contrast, full-length DciA produces a 10-fold increase (Supplementary Figure 19).

**Supplementary Figure 22.**
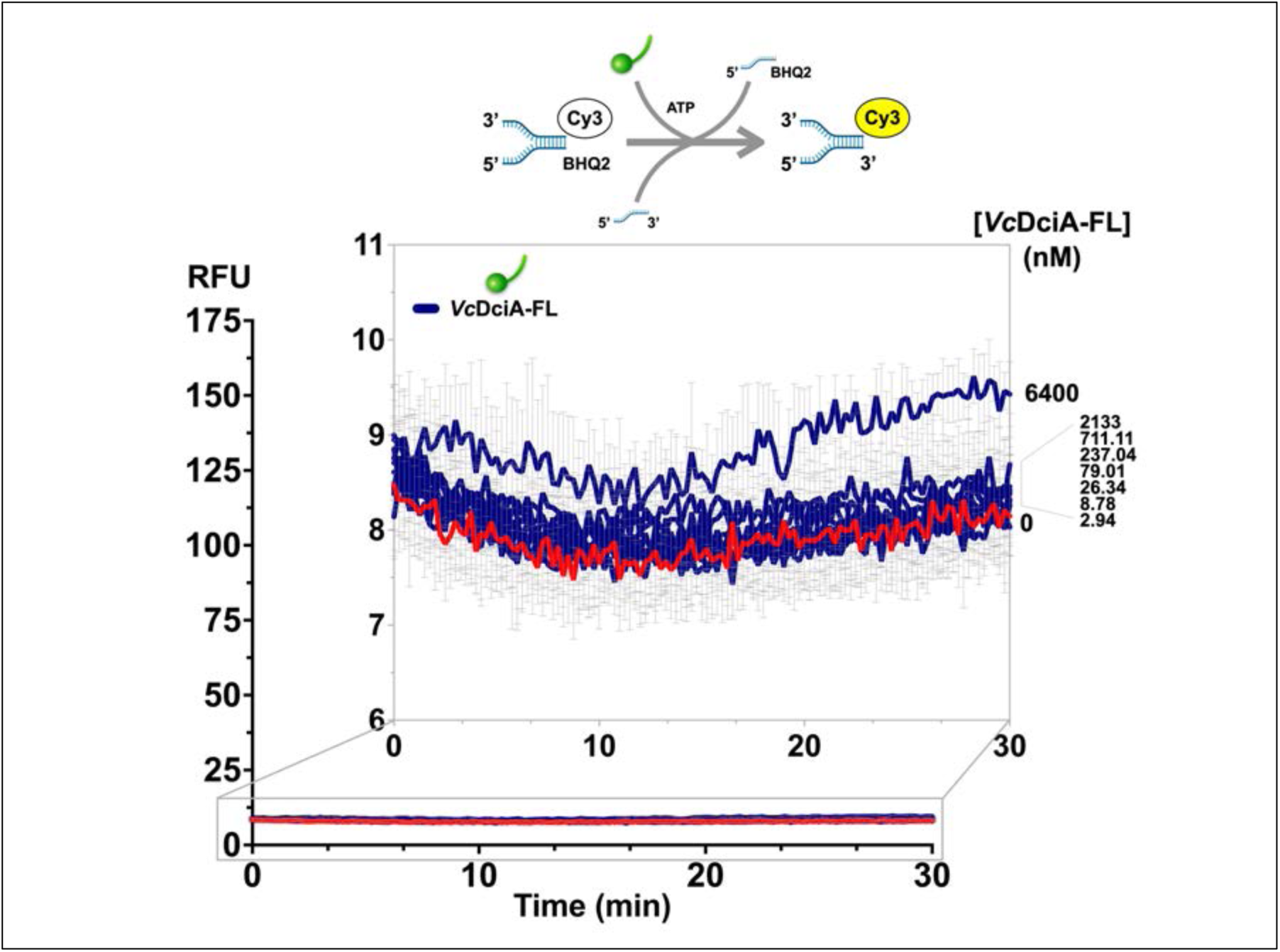
Isolated *Vc*DciA fails to unwind the 5ʹ- and 3ʹ-tailed replication fork mimic. Titration from 0-6400 nM (blue curves) of *Vc*DciA into an assay that lacks DnaB, as diagrammed schematically above, fails to show any 570 nm fluorescent signal arising from unwinding of the DNA. The inset shows a plot section expanded around RFU values of 6-11. The red curve describes fluorescent signals from an experiment that lacked DciA. These data suggest that DciA alone fails to unwind the fork DNA substrate, a finding that contrasts with the robust unwinding observed with the DNA bubble substrate. The reason for this difference in activity is unknown.

**Supplementary Figure 23.**
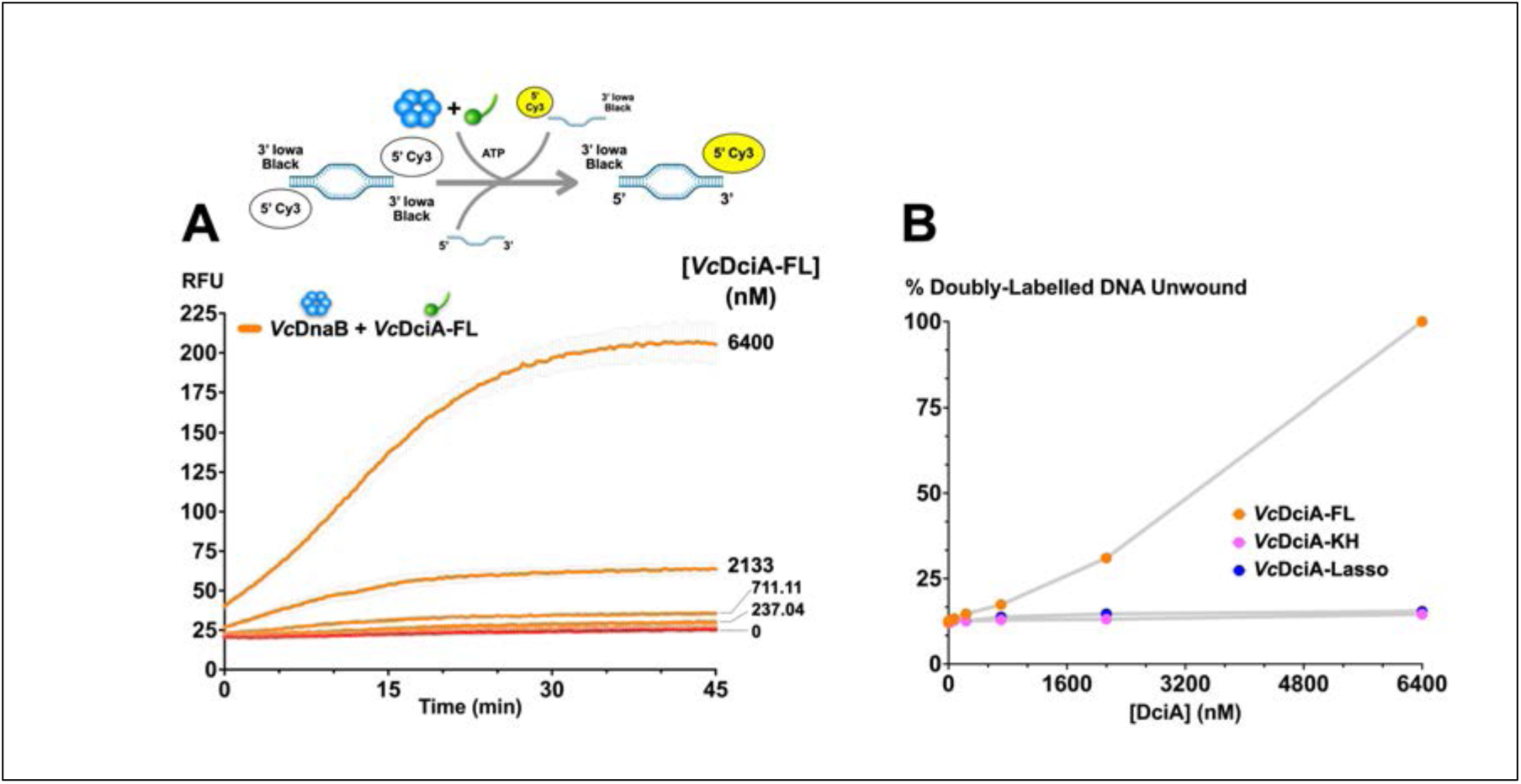
Measurement of *Vc*DciA-mediated loading of *Vc*DnaB using a doubly labeled DNA bubble. The assay diagrammed above was performed precisely as the singly labeled DNA bubble substrate in Figure 7, except that FRET labels were included at both ends of the DNA bubble. A) A time course (orange curves) of fluorescence at 570 nM was recorded from an experiment wherein *Vc*DciA was titrated from 0.11 to 6400 nM while the *Vc*DnaB concentration was fixed at 200 nM. The red curve shows the signal from an experiment that lacked *Vc*DciA. Surprisingly, the amplitude of the fluorescence signals did not increase twofold compared to the singly labeled, as was expected, given the presence of two fluorescence centers. Instead, the signal increased by 10%. B) Data from panel A are plotted as a percentage of DNA unwound as a function of *Vc*DciA concentration. These data show no loading activity by the *Vc*DciA-KH (pink) or Lasso domains (blue), even with a significant excess of each domain (Supplementary Figure 25).

**Supplementary Figure 24.**
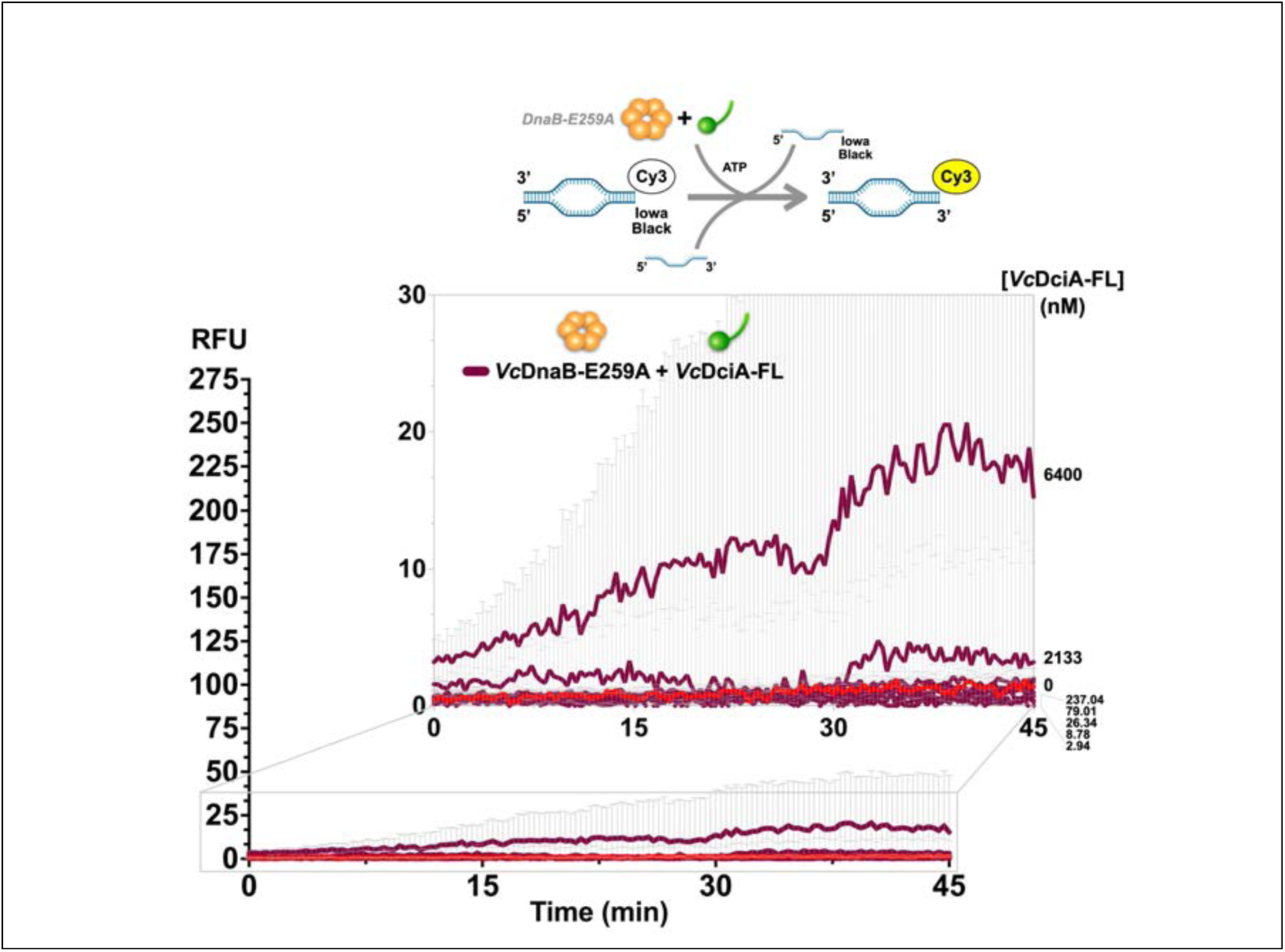
The *Vc*DnaB Walker B glutamate (E259A) mutation is deficient in loading onto a physiological bubble substrate. This experiment, diagrammed above, is similar to the one shown in Supplementary Figure 22, except that a DNA bubble substrate is used. The plot illustrates a point-by-point subtraction of the fluorescence signals generated by *Vc*DciA, as shown in Figure 7 B. The resulting curves (orange) show little loading activity above the basal rate measured without DciA (red). Only when DciA is present at 6400 nM is a curve with significant activity above the basal rate observed. However, the amplitude of this curve (∼15) was 30% of the amplitude seen with wild-type *Vc*DnaB. The inset shows the same data but expands a section of the plot.

**Supplementary Figure 25.**
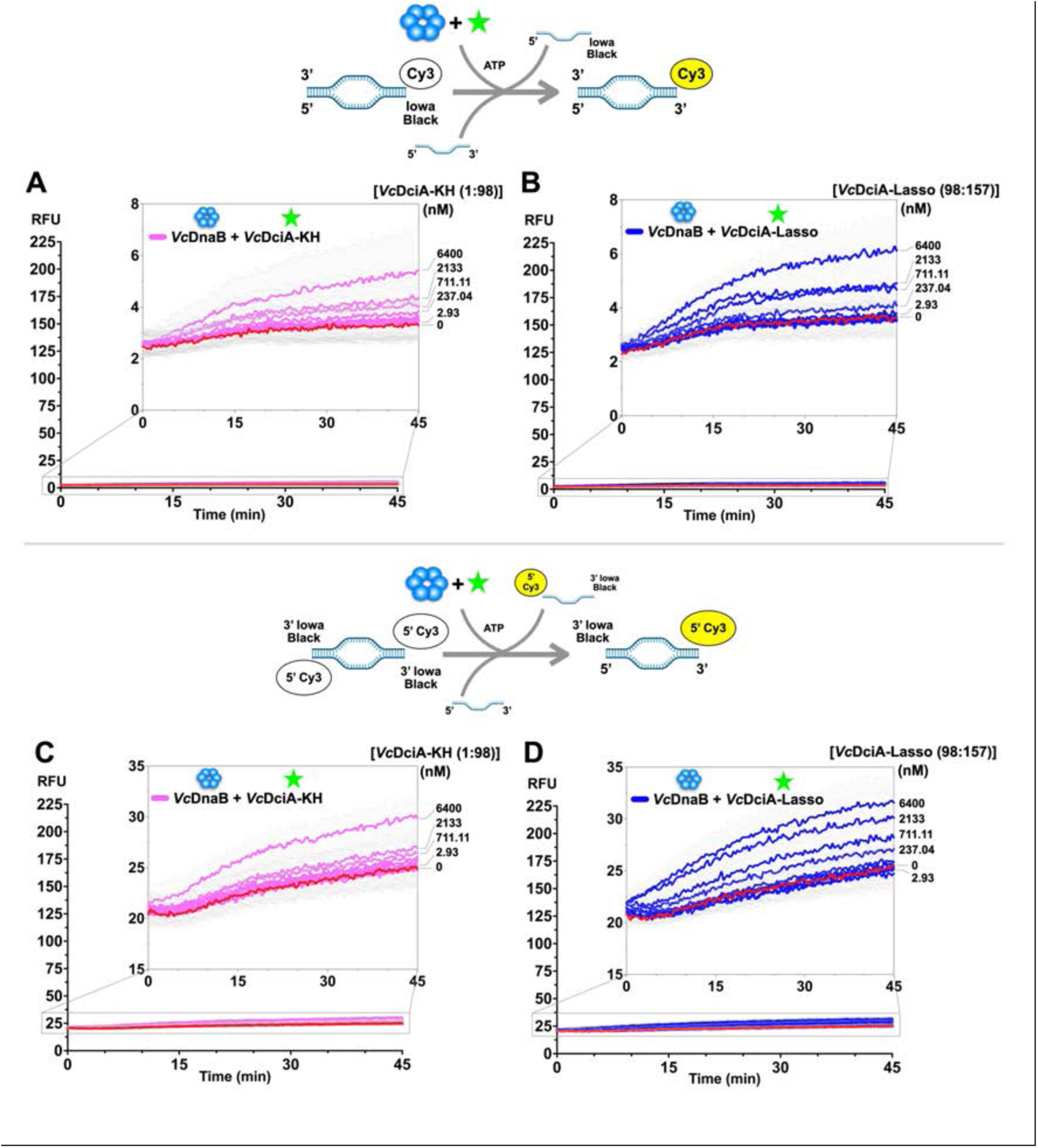
The isolated *Vc*DciA KH and Lasso domains are inactive in the bubble-loading assay. These measurements, diagrammed above each panel, represent controls for parallel studies with full-length *Vc*DciA. Panels A and B employ the singly labeled substrate, and panels C and D employ the doubly labeled substrate. Titration of the appropriate DciA construct was carried out as described in Figure 7. Measurements with the *Vc*DciA-KH domain are colored pink, and those with the *Vc*DciA-Lasso domain are colored blue. The red curves in each plot represent the basal rate of loading measured in the absence of *Vc*DciA. With the singly labeled substrate, a 1.5 to 2-fold increase in the fluorescence signal was seen at the highest concentration tested (6400 nM); in contrast, full-length *Vc*DciA produces a 10-fold increase (Figure 7). With the doubly labeled substrate, a ∼1.2-fold increase in the fluorescence signal was seen at the highest concentration tested (6400 nM); in contrast, full-length *Vc*DciA produces an 8-fold increase (Supplementary Figure 25). The lack of differentiation of the DciA titration-derived curves from the basal rate (red) leads us to conclude that each *Vc*DciA truncated construct is inactive in helicase loading.

**Supplementary Figure 26.**
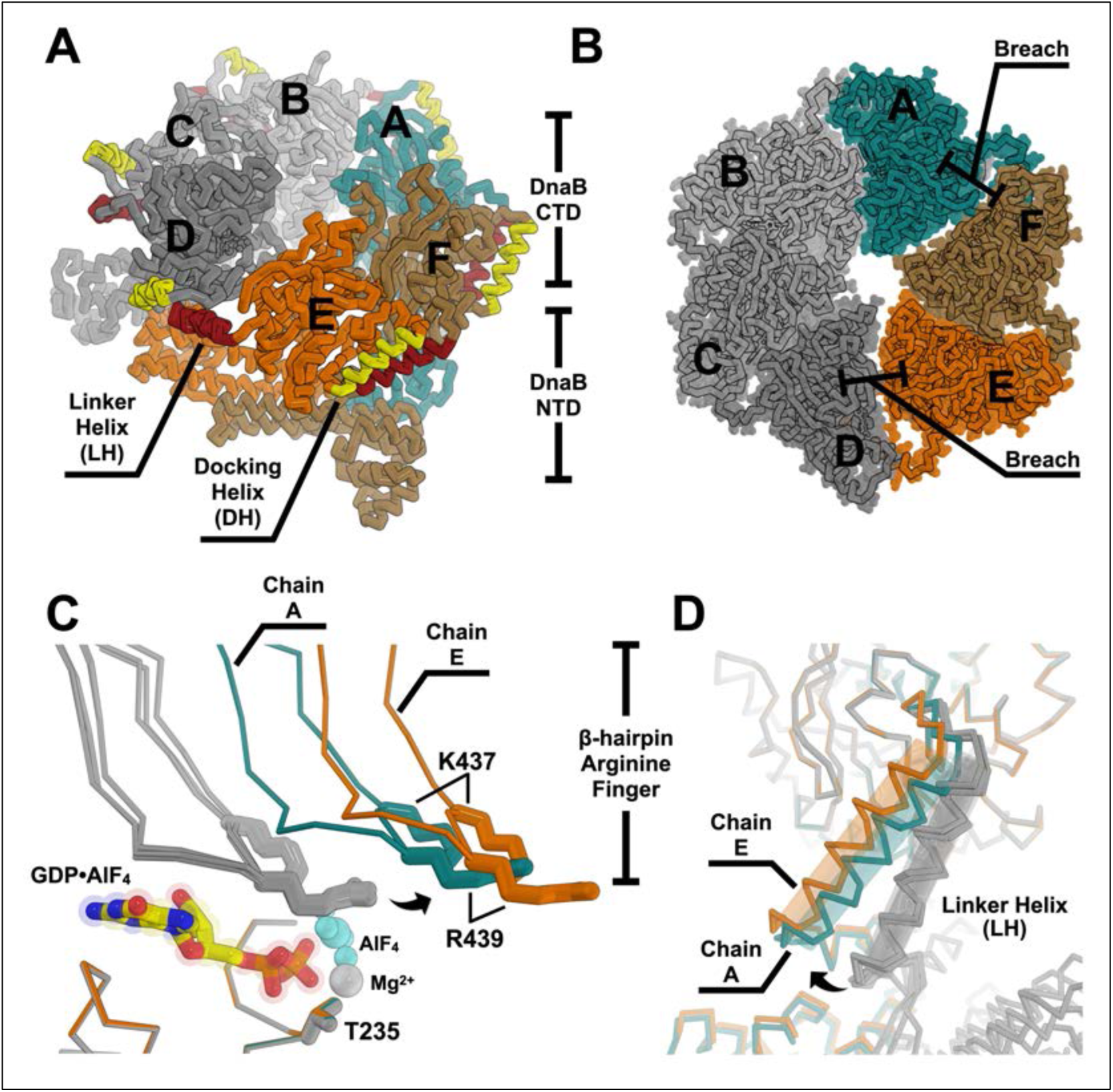
Crystal Structure of *Vc*DnaB-GDP/AlF_4_ (PDB: 6T66) and the disposition of β-hairpin arginine fingers and Linker Helix elements. A) *Vc*DnaB is depicted as a ribbon with chains labeled. The NTD and CTD layers are indicated. The six Docking-Helix (DH) and Linker-Helix (LH) elements are shown in yellow and red, respectively. Chains A, F, and E are shown in blue, sand, and orange, and the other chains are in alternating shades of gray. (A) The 6T66 DnaB structure depicts the two breaches in the CTD layer: the largest is between chains E (orange) and D (grey), and the narrowest is between chains A (deep teal) and F (sand). The protein surface is drawn with the PyMol solvent radius = 0.3 setting. C) The six CTD pairs (A-B, B-C, etc., residue 200:461) from 6T66 were superimposed (RMSD = 0.03 Å) on chain A, but the DH elements (residue 280:306) and the second chain were excluded from the calculation. Each *Vc*DnaB harbors a GDP•AlF_4_ species, and only one is shown in the stick representation, which has transparent spheres overlaid. The β-hairpin arginine fingers are shown as ribbons, and K437 and K439 are shown as sticks. The β-hairpins from four chains (B, C, D, F) follow each other closely (grey), although not included in the alignment. The corresponding elements from chain A (deep teal) and E (orange) have moved away by ∼9 Å and ∼13 Å, respectively. D) The superposition from panel C was interrogated for the disposition of the LH elements represented as transparent cylinders. The LH elements from chains B, C, D, and F (grey) superimpose closely and occupy the LH1 position as defined in the main text and Supplementary Figure 10. In contrast, those belonging to chain A (deep teal) and chain E (orange) drift away by ∼5-∼9 Å and occupy position LH2.

**Supplementary Figure 27.**
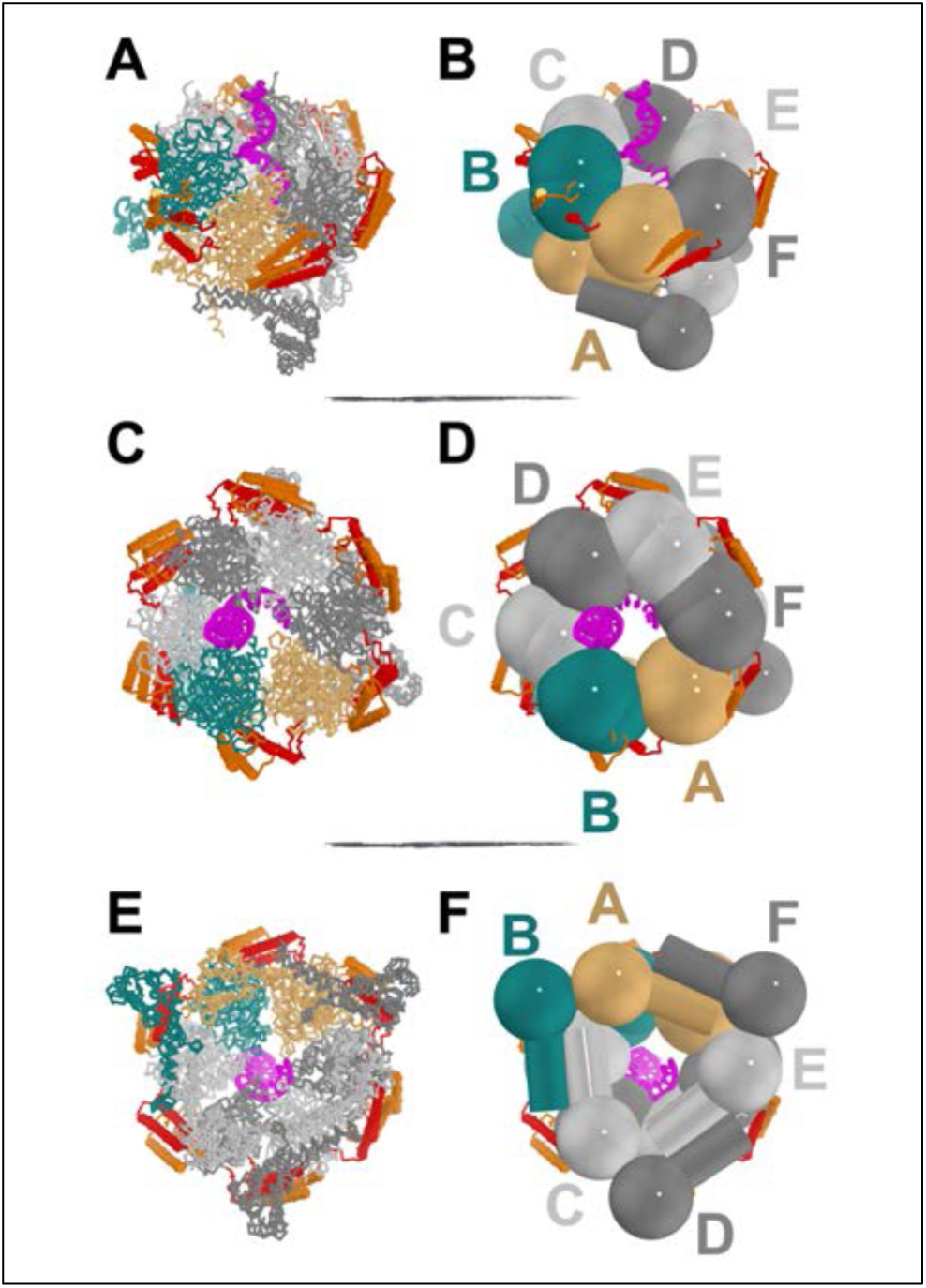
Distinct Configuration in DnaB in the VcDnaB-ssDNA-ATPγS complex and the DnaB-DciA complex (PDB: 8A3V). The DnaB-DciA crystal structure was superimposed on the chain A NTD (residue 1:170) of the current VcDnaB-ssDNA-ATPγS complex (RMSD: 0.85 Å; 139 Cα carbons). DnaB is drawn in a ribbon representation (A, C, and E) or a schematic using the NTD-cylinder/sphere and CTD-spheres design language (B, D, and F) in the identical poses. ssDNA, colored in magenta, is depicted in a cartoon and stick representation. The superimposed structures are colored identically. Chain A (light orange), chain B (deep teal), and other DnaB subunits (colored in alternating shades of gray) are labeled. The Docking Helix (DH) and the Linker Helix (LH) are depicted as cylinders and ribbons colored orange and red. Panels A/B and D/E are related by a 30° rotation about the X-axis, and panels D/E and E/F are related by 180°. The dilated configuration of the NTD tier enabled superposition even of chains not included in the calculation (panels E and F), as indicated by the close positions of the NTD sphere and cylinder proxies. In contrast, individual CTDs vary between the two structures in this superposition. The distinct conformations in the two DnaB structures can be seen in the positions of the DH and LH elements.

**Supplementary Movie 1. Migration of Subunits from the Bottom to the Top of the DnaB Spiral during Translocation.** This video shows the transition from the ATPγS conformation of the present *Vc*DnaB-ssDNA complex to the GDP-AlF_4_ configuration of the GDP•AlF4 4ESV DnaB ssDNA (12) complex. The two complexes are aligned by superimposing the ssDNA molecules of each structure, as detailed in the main text. The motions depicted in the video are described in the main text. The DnaB-ssDNA-ATPγS complex is drawn in the ribbon representation (left) or the sphere/cylinder/ribbon schematic (right). Both representations are colored according to their chain. The ssDNA is depicted in both cartoon and stick representations and is colored orange. To calculate the transition between structures, a version of the 4ESV coordinates was constructed that encompassed the *Vibrio cholerae* primary sequence. The two states were then submitted to PyMol to calculate the 90 intermediate structures between the ATPγS and the GDP-AlF_4_ conformers, as well as an additional 90 structures between the GDP-AlF_4_ and the ATPγS structures. Frames were rendered in PyMol and assembled in QuickTime Player.

## Supplementary Tables

**Supplementary Table 1.** Expression Plasmids Used in This Study.

| Expression Plasmid | Insert | Template | Primers | Plasmid vector |
| --- | --- | --- | --- | --- |
| pNG066-pET24a-VcDnaB-unt | VcDnaB-untagged | *V. cholerae* N16961 genomic DNA | VcDnaB-NdeI-F, VcDnaB-XhoI-R | pET24a |
| pNG067-pET24a-VcDnaB-NHis | VcDnaB-NHis | pNG066 | VcDnaB-pET24a-NHis-F, VcDnaB-pET24a-NHis-R | pET24a |
| pNG072-pET24a-VcDnaB-E259A-NHis | VcDnaB-E259A-NHis | pNG067 | VcDnaB-E259A-F, VcDnaB-E259A-R | pET24a |
| pNG114-pET24a-VcDnaB-1to170-unt | VcDnaB-NTD-untagged | pNG066 | VcDnaB_170-pET24a-F, VcDnaB_170-pET24a-R | pET24a |
| pNG131-pET24a-VcDnaB-1to208-unt | VcDnaB-NTD_LH-untagged | pNG066 | VcDnaB_208-pET24a-F, VcDnaB_208-pET24a-R | pET24a |
| pNG134-pET24a-VcDnaB-170to468-unt | VcDnaB-LH_CTD-untagged | pNG066 | pET24a-VcDnaB_170-F, pET24a-VcDnaB_170-R | pET24a |
| pNG057-pCDFDuet-1-VcDciA-CHis | VcDciA-CHis | *V. cholerae* N16961 genomic DNA | VcDciA-NdeI-F, VcDciA-XhoI-CHis-R | pCDFDuet-1 |
| pNG078-pCDFDuet-VcDciA-1to98-CHis | VcDciA-NTD-CHis | pNG057 | VcDciA_98-CHis_pCDF-F, VcDciA_98-CHis_pCDF-R | pCDFDuet-1 |
| pNG087-pCDFDuet-VcDciA-98to157-CHis | VcDciA-CTD-CHis | pNG057 | VcDciA_1-VcDciA_98-F, VcDciA_1-VcDciA_98-R | pCDFDuet-1 |

**Supplementary Table 2.**
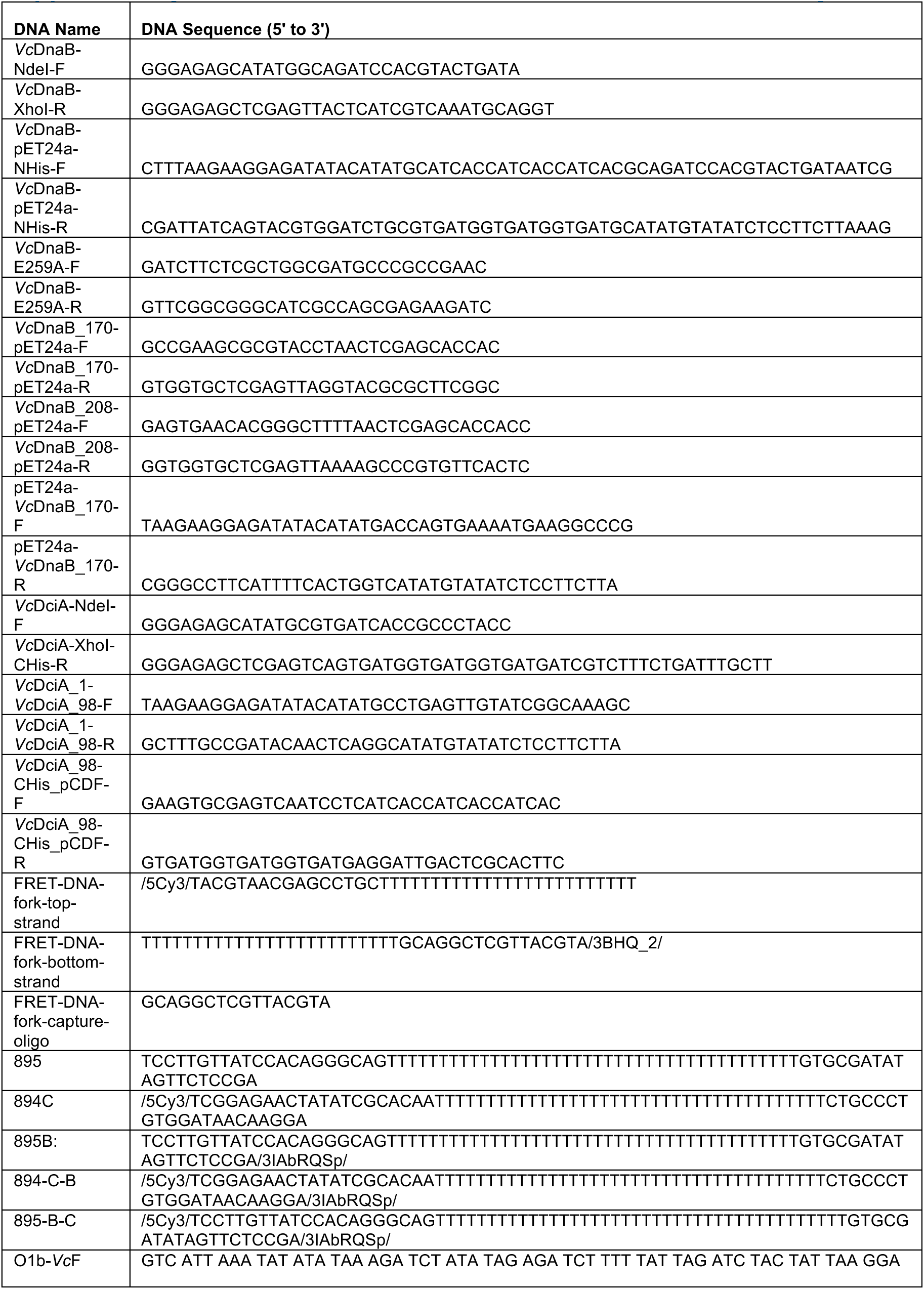
Primers and Other DNAs Used in This Study.

**Supplementary Table 3:**
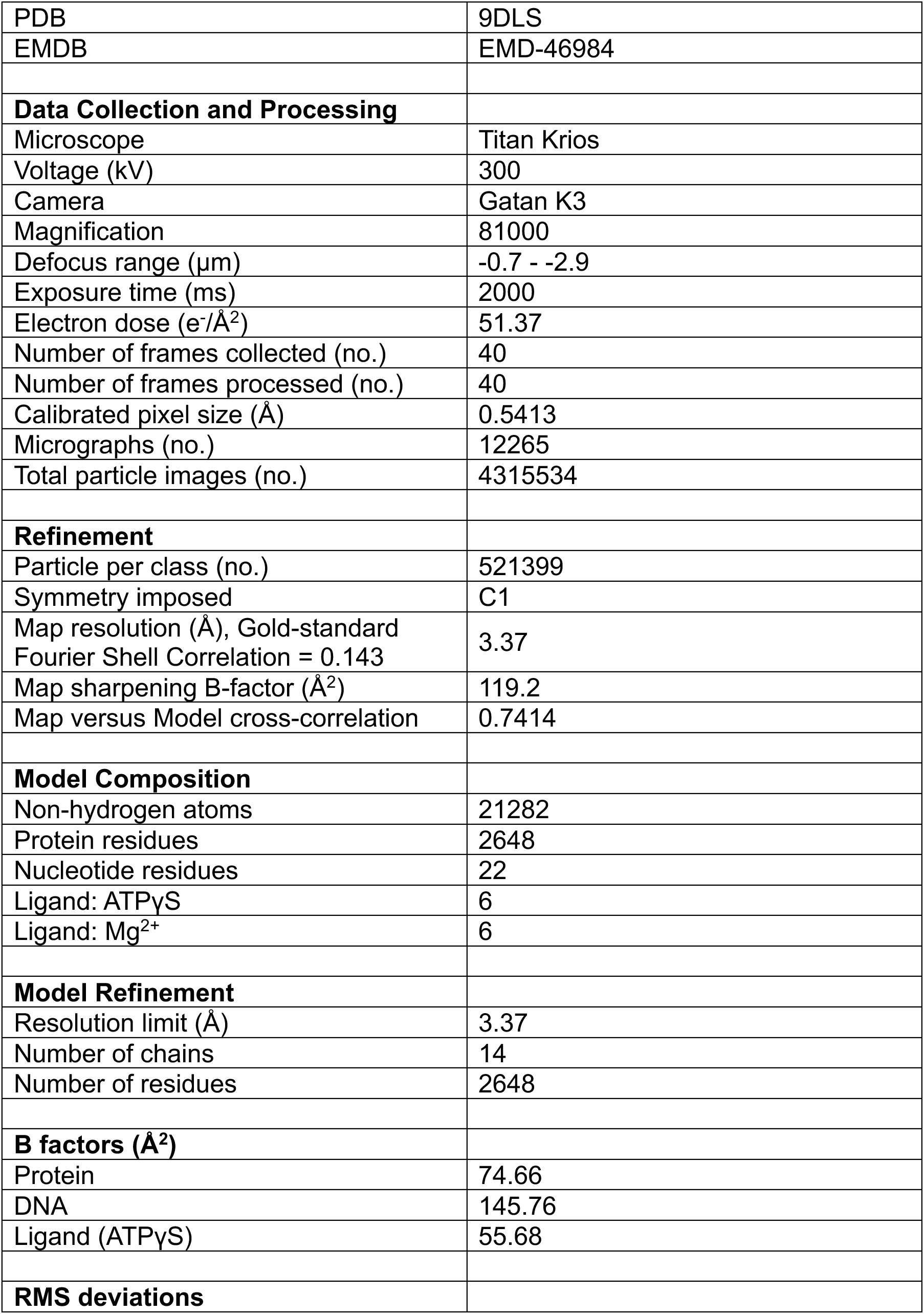
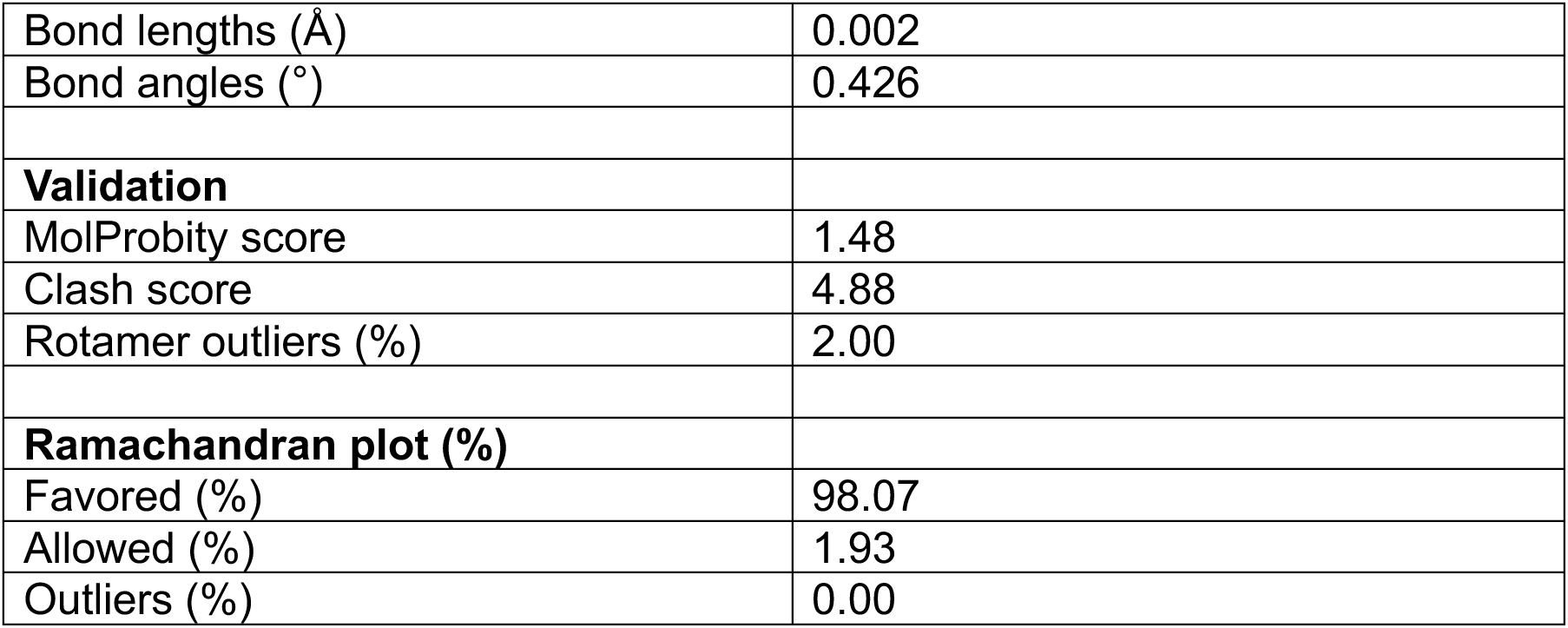
Cryogenic Electron Microscopy Data Measurement and PDB Model Refinement.

**Supplementary Table 4.** Contacts (<4 Å) in the VcDnaB-ssDNA complex.

| Model1 | Chain1 | Res1 | ResN1 | Atom1 | to | Model2 | Chain2 | Res2 | ResN2 | Atom2 | Dist |
| --- | --- | --- | --- | --- | --- | --- | --- | --- | --- | --- | --- |
| VcB-ssDNA | B | THR | 355 | OG1 | to | VcB-ssDNA | G | DC | 9 | O2 | 3.947 |
| VcB-ssDNA | B | ASN | 383 | OD1 | to | VcB-ssDNA | G | DC | 11 | C5' | 3.716 |
| VcB-ssDNA | B | ASN | 383 | ND2 | to | VcB-ssDNA | G | DC | 11 | O5' | 3.771 |
| VcB-ssDNA | B | ASN | 383 | OD1 | to | VcB-ssDNA | G | DC | 11 | O5' | 3.727 |
| VcB-ssDNA | B | ASN | 383 | CG | to | VcB-ssDNA | G | DC | 11 | OP1 | 3.107 |
| VcB-ssDNA | B | ASN | 383 | ND2 | to | VcB-ssDNA | G | DC | 11 | OP1 | 2.651 |
| VcB-ssDNA | B | ASN | 383 | OD1 | to | VcB-ssDNA | G | DC | 11 | OP1 | 2.814 |
| VcB-ssDNA | B | ASN | 383 | ND2 | to | VcB-ssDNA | G | DC | 11 | OP2 | 3.263 |
| VcB-ssDNA | B | ASN | 383 | CG | to | VcB-ssDNA | G | DC | 11 | P | 3.922 |
| VcB-ssDNA | B | ASN | 383 | ND2 | to | VcB-ssDNA | G | DC | 11 | P | 3.259 |
| VcB-ssDNA | B | ASN | 383 | OD1 | to | VcB-ssDNA | G | DC | 11 | P | 3.784 |
| VcB-ssDNA | B | ASN | 383 | CA | to | VcB-ssDNA | G | DA | 12 | OP2 | 3.423 |
| VcB-ssDNA | B | ASN | 383 | CB | to | VcB-ssDNA | G | DA | 12 | OP2 | 3.404 |
| VcB-ssDNA | B | ASN | 383 | CG | to | VcB-ssDNA | G | DA | 12 | OP2 | 3.126 |
| VcB-ssDNA | B | ASN | 383 | OD1 | to | VcB-ssDNA | G | DA | 12 | OP2 | 2.483 |
| VcB-ssDNA | B | ASN | 383 | OD1 | to | VcB-ssDNA | G | DA | 12 | P | 3.825 |
| VcB-ssDNA | B | ARG | 400 | CA | to | VcB-ssDNA | G | DC | 11 | OP1 | 3.351 |
| VcB-ssDNA | B | ARG | 400 | CD | to | VcB-ssDNA | G | DA | 12 | OP2 | 3.746 |
| VcB-ssDNA | B | ARG | 400 | NH1 | to | VcB-ssDNA | G | DA | 12 | OP2 | 3.77 |
| VcB-ssDNA | B | GLU | 401 | CA | to | VcB-ssDNA | G | DA | 10 | C4' | 3.997 |
| VcB-ssDNA | B | GLU | 401 | CA | to | VcB-ssDNA | G | DA | 10 | C5' | 3.812 |
| VcB-ssDNA | B | GLU | 401 | CA | to | VcB-ssDNA | G | DA | 10 | O3' | 3.636 |
| VcB-ssDNA | B | GLY | 403 | CA | to | VcB-ssDNA | G | DA | 10 | OP1 | 3.887 |
| VcB-ssDNA | C | THR | 355 | OG1 | to | VcB-ssDNA | G | DA | 8 | C5' | 3.886 |
| VcB-ssDNA | C | ASN | 383 | ND2 | to | VcB-ssDNA | G | DC | 9 | O5' | 3.47 |
| VcB-ssDNA | C | ASN | 383 | CG | to | VcB-ssDNA | G | DC | 9 | OP1 | 3.534 |
| VcB-ssDNA | C | ASN | 383 | ND2 | to | VcB-ssDNA | G | DC | 9 | OP1 | 3.136 |
| VcB-ssDNA | C | ASN | 383 | OD1 | to | VcB-ssDNA | G | DC | 9 | OP1 | 3.236 |
| VcB-ssDNA | C | ASN | 383 | ND2 | to | VcB-ssDNA | G | DC | 9 | OP2 | 2.885 |
| VcB-ssDNA | C | ASN | 383 | ND2 | to | VcB-ssDNA | G | DC | 9 | P | 3.222 |
| VcB-ssDNA | C | ARG | 384 | NH2 | to | VcB-ssDNA | G | DA | 10 | O3' | 3.553 |
| VcB-ssDNA | C | ARG | 384 | NH2 | to | VcB-ssDNA | G | DA | 10 | O5' | 3.969 |
| VcB-ssDNA | C | ARG | 384 | CB | to | VcB-ssDNA | G | DA | 10 | OP2 | 3.621 |
| VcB-ssDNA | C | ARG | 384 | NH2 | to | VcB-ssDNA | G | DC | 11 | OP1 | 3.334 |
| VcB-ssDNA | C | ARG | 384 | NH2 | to | VcB-ssDNA | G | DC | 11 | OP2 | 3.229 |
| VcB-ssDNA | C | ARG | 384 | NH2 | to | VcB-ssDNA | G | DC | 11 | P | 3.476 |
| VcB-ssDNA | C | ARG | 400 | NH2 | to | VcB-ssDNA | G | DC | 9 | O3' | 3.991 |
| VcB-ssDNA | C | ARG | 400 | CA | to | VcB-ssDNA | G | DC | 9 | OP1 | 3.163 |
| VcB-ssDNA | C | ARG | 400 | CB | to | VcB-ssDNA | G | DC | 9 | OP1 | 3.945 |
| VcB-ssDNA | C | GLU | 401 | CA | to | VcB-ssDNA | G | DA | 8 | O3' | 3.771 |
| VcB-ssDNA | C | GLU | 401 | CA | to | VcB-ssDNA | G | DC | 9 | OP1 | 3.548 |
| VcB-ssDNA | C | GLY | 403 | CA | to | VcB-ssDNA | G | DA | 8 | OP1 | 3.765 |
| VcB-ssDNA | D | ASN | 353 | CG | to | VcB-ssDNA | G | DG | 5 | N2 | 3.974 |
| VcB-ssDNA | D | ASN | 353 | OD1 | to | VcB-ssDNA | G | DG | 5 | N2 | 3.119 |
| VcB-ssDNA | D | ASN | 383 | ND2 | to | VcB-ssDNA | G | DT | 7 | O5' | 3.535 |
| VcB-ssDNA | D | ASN | 383 | CG | to | VcB-ssDNA | G | DT | 7 | OP1 | 3.881 |
| VcB-ssDNA | D | ASN | 383 | ND2 | to | VcB-ssDNA | G | DT | 7 | OP1 | 3.317 |
| VcB-ssDNA | D | ASN | 383 | ND2 | to | VcB-ssDNA | G | DT | 7 | OP2 | 2.826 |
| VcB-ssDNA | D | ASN | 383 | OD1 | to | VcB-ssDNA | G | DA | 8 | OP2 | 3.144 |
| VcB-ssDNA | D | ASN | 383 | ND2 | to | VcB-ssDNA | G | DT | 7 | P | 3.287 |
| VcB-ssDNA | D | ARG | 384 | NH1 | to | VcB-ssDNA | G | DC | 9 | OP1 | 3.975 |
| VcB-ssDNA | D | ARG | 384 | CB | to | VcB-ssDNA | G | DA | 8 | OP2 | 3.817 |
| VcB-ssDNA | D | ARG | 384 | NH1 | to | VcB-ssDNA | G | DC | 9 | OP2 | 3.673 |
| VcB-ssDNA | D | ARG | 400 | CA | to | VcB-ssDNA | G | DT | 7 | OP1 | 3.31 |
| VcB-ssDNA | D | ARG | 400 | NH1 | to | VcB-ssDNA | G | DA | 8 | OP1 | 3.523 |
| VcB-ssDNA | D | ARG | 400 | NH1 | to | VcB-ssDNA | G | DA | 8 | OP2 | 3.988 |
| VcB-ssDNA | D | GLU | 401 | CA | to | VcB-ssDNA | G | DA | 6 | C5' | 3.914 |
| VcB-ssDNA | D | GLU | 401 | CA | to | VcB-ssDNA | G | DA | 6 | O3' | 3.561 |
| VcB-ssDNA | D | GLU | 401 | CG | to | VcB-ssDNA | G | DA | 6 | O3' | 3.774 |
| VcB-ssDNA | D | GLU | 401 | CA | to | VcB-ssDNA | G | DT | 7 | OP1 | 3.923 |
| VcB-ssDNA | D | GLY | 403 | CA | to | VcB-ssDNA | G | DA | 6 | OP1 | 3.815 |
| VcB-ssDNA | E | THR | 355 | OG1 | to | VcB-ssDNA | G | DA | 4 | C5' | 3.779 |
| VcB-ssDNA | E | THR | 355 | OG1 | to | VcB-ssDNA | G | DC | 3 | O2 | 3.494 |
| VcB-ssDNA | E | THR | 355 | OG1 | to | VcB-ssDNA | G | DA | 4 | O4' | 3.947 |
| VcB-ssDNA | E | ASN | 383 | OD1 | to | VcB-ssDNA | G | DG | 5 | C3' | 3.914 |
| VcB-ssDNA | E | ASN | 383 | CG | to | VcB-ssDNA | G | DG | 5 | O5' | 3.967 |
| VcB-ssDNA | E | ASN | 383 | ND2 | to | VcB-ssDNA | G | DG | 5 | O5' | 3.376 |
| VcB-ssDNA | E | ASN | 383 | CG | to | VcB-ssDNA | G | DG | 5 | OP1 | 3.957 |
| VcB-ssDNA | E | ASN | 383 | ND2 | to | VcB-ssDNA | G | DG | 5 | OP1 | 3.414 |
| VcB-ssDNA | E | ASN | 383 | CG | to | VcB-ssDNA | G | DG | 5 | OP2 | 3.968 |
| VcB-ssDNA | E | ASN | 383 | ND2 | to | VcB-ssDNA | G | DG | 5 | OP2 | 2.769 |
| VcB-ssDNA | E | ASN | 383 | OD1 | to | VcB-ssDNA | G | DA | 6 | OP2 | 3.377 |
| VcB-ssDNA | E | ASN | 383 | ND2 | to | VcB-ssDNA | G | DG | 5 | P | 3.268 |
| VcB-ssDNA | E | ARG | 384 | CB | to | VcB-ssDNA | G | DA | 6 | OP2 | 3.604 |
| VcB-ssDNA | E | ARG | 384 | NH2 | to | VcB-ssDNA | G | DT | 7 | OP2 | 3.483 |
| VcB-ssDNA | E | ARG | 400 | CA | to | VcB-ssDNA | G | DG | 5 | OP1 | 3.17 |
| VcB-ssDNA | E | ARG | 400 | CB | to | VcB-ssDNA | G | DG | 5 | OP1 | 3.856 |
| VcB-ssDNA | E | GLU | 401 | CA | to | VcB-ssDNA | G | DA | 4 | C4' | 3.884 |
| VcB-ssDNA | E | GLU | 401 | CA | to | VcB-ssDNA | G | DA | 4 | C5' | 3.87 |
| VcB-ssDNA | E | GLU | 401 | CA | to | VcB-ssDNA | G | DA | 4 | O3' | 3.897 |
| VcB-ssDNA | E | GLY | 403 | CA | to | VcB-ssDNA | G | DA | 4 | OP1 | 3.625 |
| VcB-ssDNA | F | THR | 355 | OG1 | to | VcB-ssDNA | G | DC | 2 | C5' | 3.719 |
| VcB-ssDNA | F | THR | 355 | OG1 | to | VcB-ssDNA | G | DT | 1 | O4' | 3.647 |
| VcB-ssDNA | F | ASN | 383 | CG | to | VcB-ssDNA | G | DC | 3 | O5' | 3.901 |
| VcB-ssDNA | F | ASN | 383 | ND2 | to | VcB-ssDNA | G | DC | 3 | O5' | 3.271 |
| VcB-ssDNA | F | ASN | 383 | CG | to | VcB-ssDNA | G | DC | 3 | OP1 | 3.928 |
| VcB-ssDNA | F | ASN | 383 | ND2 | to | VcB-ssDNA | G | DC | 3 | OP1 | 3.269 |
| VcB-ssDNA | F | ASN | 383 | ND2 | to | VcB-ssDNA | G | DC | 3 | OP2 | 3.048 |
| VcB-ssDNA | F | ASN | 383 | OD1 | to | VcB-ssDNA | G | DA | 4 | OP2 | 3.653 |
| VcB-ssDNA | F | ASN | 383 | ND2 | to | VcB-ssDNA | G | DC | 3 | P | 3.276 |
| VcB-ssDNA | F | ARG | 384 | NH2 | to | VcB-ssDNA | G | DA | 4 | C3' | 3.931 |
| VcB-ssDNA | F | ARG | 384 | NH2 | to | VcB-ssDNA | G | DG | 5 | OP1 | 3.545 |
| VcB-ssDNA | F | ARG | 384 | CB | to | VcB-ssDNA | G | DA | 4 | OP2 | 3.528 |
| VcB-ssDNA | F | ARG | 384 | NH2 | to | VcB-ssDNA | G | DG | 5 | OP2 | 3.591 |
| VcB-ssDNA | F | ARG | 384 | NH2 | to | VcB-ssDNA | G | DG | 5 | P | 3.861 |
| VcB-ssDNA | F | ARG | 400 | CA | to | VcB-ssDNA | G | DC | 3 | OP1 | 3.216 |
| VcB-ssDNA | F | ARG | 400 | CB | to | VcB-ssDNA | G | DC | 3 | OP1 | 3.778 |
| VcB-ssDNA | F | GLU | 401 | CA | to | VcB-ssDNA | G | DC | 2 | C4' | 3.918 |
| VcB-ssDNA | F | GLU | 401 | CG | to | VcB-ssDNA | G | DC | 2 | C4' | 3.794 |
| VcB-ssDNA | F | GLU | 401 | CA | to | VcB-ssDNA | G | DC | 2 | C5' | 3.855 |
| VcB-ssDNA | F | GLU | 401 | OE2 | to | VcB-ssDNA | G | DC | 3 | C5' | 3.76 |
| VcB-ssDNA | F | GLU | 401 | CG | to | VcB-ssDNA | G | DC | 2 | O3' | 3.81 |

**Supplementary Table 5.**
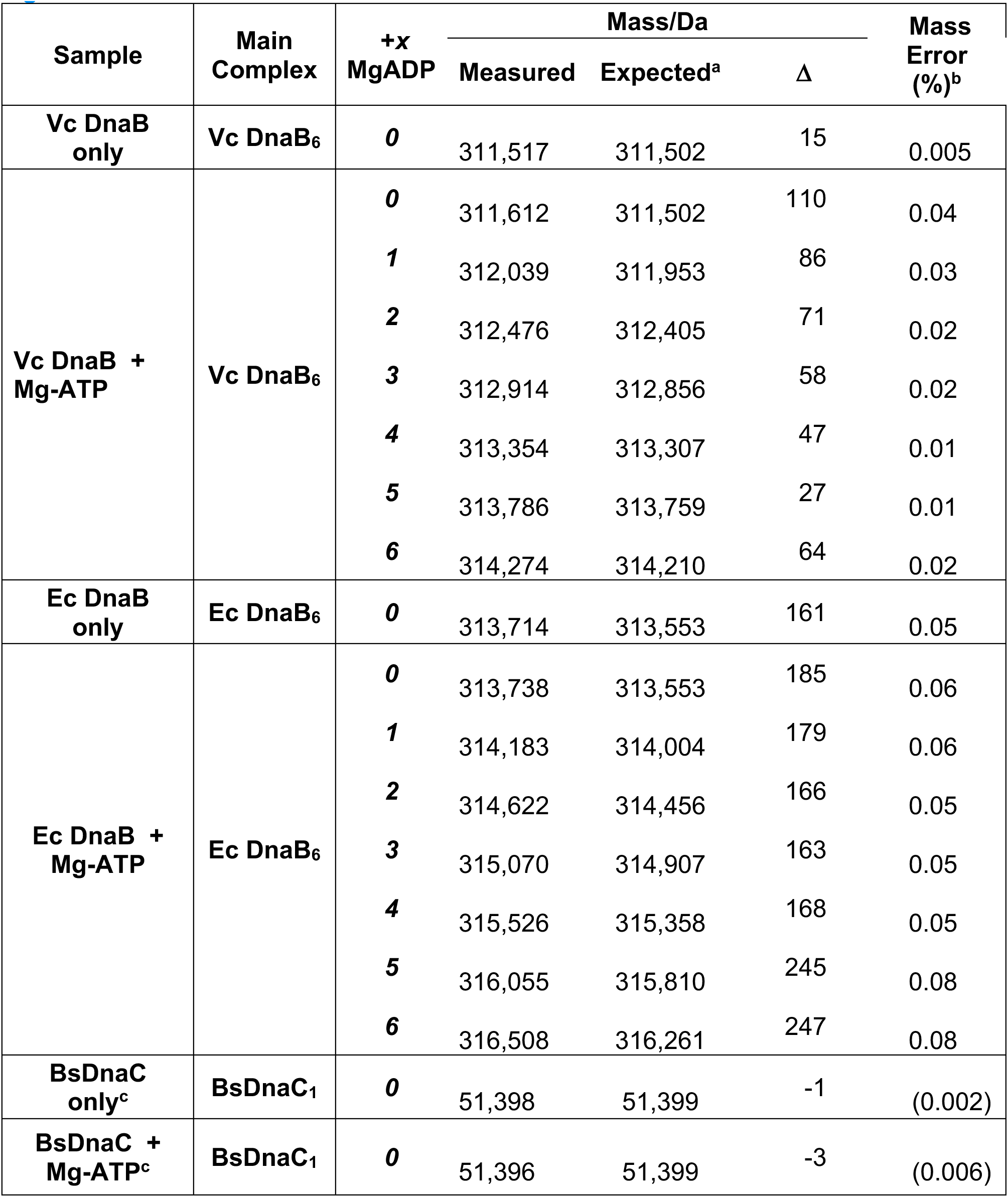
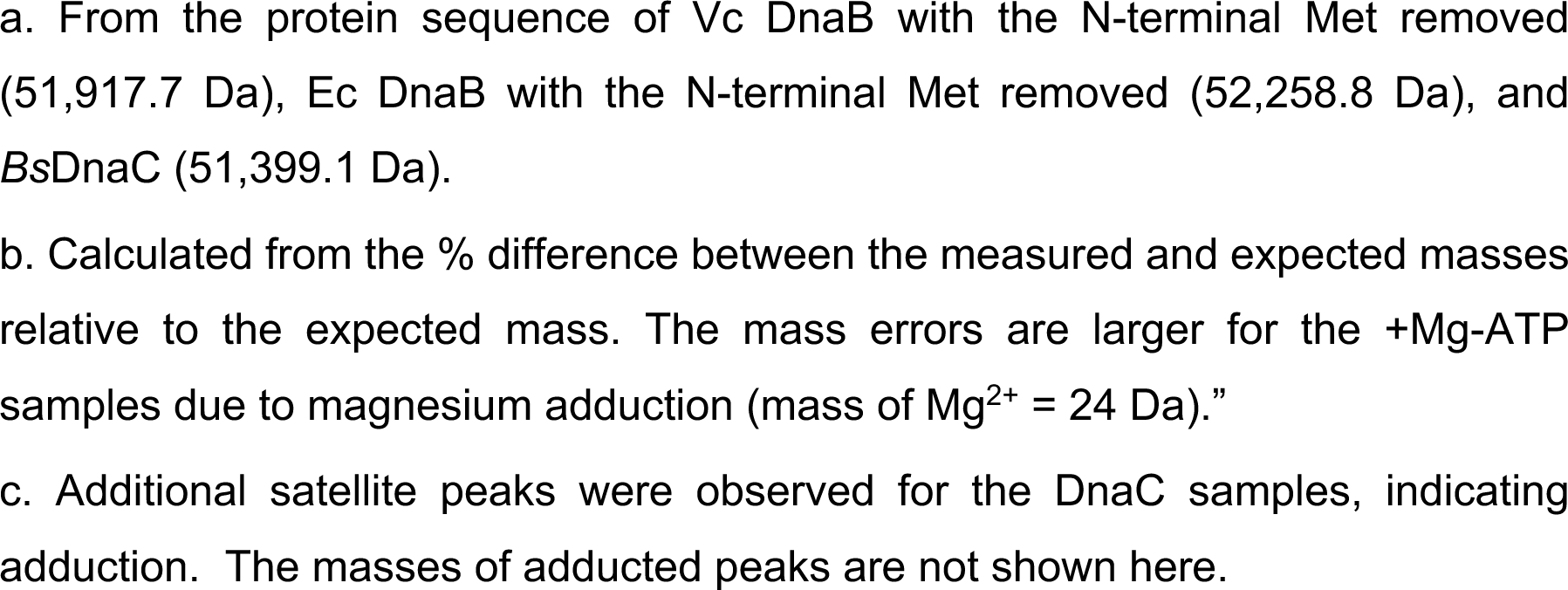
native mass spectrometry mass measurements for predominant DnaB assemblies from three bacterial species with and without Mg-ATP.

| Sample | Main Complex | +x<br>MgADP | Mass/Da |  |  | Mass Error (%)<sup>b</sup> |
| --- | --- | --- | --- | --- | --- | --- |
| | | | Measured | Expected<sup>a</sup> | $\Delta$ | |
| **Vc DnaB only** | **Vc DnaB<sub>6</sub>** | **0** | 311,517 | 311,502 | 15 | 0.005 |
| **Vc DnaB + Mg-ATP** | **Vc DnaB<sub>6</sub>** | **0** | 311,612 | 311,502 | 110 | 0.04 |
|  |  | **1** | 312,039 | 311,953 | 86 | 0.03 |
|  |  | **2** | 312,476 | 312,405 | 71 | 0.02 |
|  |  | **3** | 312,914 | 312,856 | 58 | 0.02 |
|  |  | **4** | 313,354 | 313,307 | 47 | 0.01 |
|  |  | **5** | 313,786 | 313,759 | 27 | 0.01 |
|  |  | **6** | 314,274 | 314,210 | 64 | 0.02 |
| **Ec DnaB only** | **Ec DnaB<sub>6</sub>** | **0** | 313,714 | 313,553 | 161 | 0.05 |
| **Ec DnaB + Mg-ATP** | **Ec DnaB<sub>6</sub>** | **0** | 313,738 | 313,553 | 185 | 0.06 |
|  |  | **1** | 314,183 | 314,004 | 179 | 0.06 |
|  |  | **2** | 314,622 | 314,456 | 166 | 0.05 |
|  |  | **3** | 315,070 | 314,907 | 163 | 0.05 |
|  |  | **4** | 315,526 | 315,358 | 168 | 0.05 |
|  |  | **5** | 316,055 | 315,810 | 245 | 0.08 |
|  |  | **6** | 316,508 | 316,261 | 247 | 0.08 |
| **BsDnaC only<sup>c</sup>** | **BsDnaC<sub>1</sub>** | **0** | 51,398 | 51,399 | -1 | (0.002) |
| **BsDnaC + Mg-ATP<sup>c</sup>** | **BsDnaC<sub>1</sub>** | **0** | 51,396 | 51,399 | -3 | (0.006) |

a. From the protein sequence of Vc DnaB with the N-terminal Met removed (51,917.7 Da), Ec DnaB with the N-terminal Met removed (52,258.8 Da), and *BsDnaC* (51,399.1 Da). b. Calculated from the % difference between the measured and expected masses relative to the expected mass. The mass errors are larger for the +Mg-ATP samples due to magnesium adduction (mass of $\text{Mg}^{2+}$ = 24 Da)." c. Additional satellite peaks were observed for the DnaC samples, indicating adduction. The masses of adducted peaks are not shown here.

